# Protective pan-betacoronavirus neutralizing antibodies by vaccination

**DOI:** 10.64898/2026.08.06.743418

**Authors:** Panpan Zhou, Ziqi Feng, Wan-ting He, Yuxin Zhu, Meng Yuan, Xuduo Li, Yuexiu Zhang, Lina Vo, Tazio Capozzola, Sean Callaghan, Nitesh Mishra, Gabriel Avillion, Katharina Dueker, Bo Liang, Rohan Roy Chowdhury, Rebecca Nedellec, Wen-Hsin Lee, Joel D. Allen, Agnes Walsh, Mariane Melo, Eileen T. McAnarney, Nisha Asok Kumar, William Rinaldi, Melissa Ferguson, Max M. Crispin, Andrew B. Ward, Darrell J. Irvine, Mohamad-Gabriel Alameh, Drew Weissman, Ralph S. Baric, Lisa E. Gralinski, Ian A. Wilson, Dennis R. Burton, Raiees Andrabi

**Author notes:** Corresponding author. (I.A.W.); (D.R.B.); (R.A.). Shanghai Institute of Virology, Shanghai Jiao Tong University School of Medicine, Shanghai 200025, China (Panpan Zhou). These authors contributed equally to this work (Panpan Zhou, Ziqi Feng, Wan-ting He).

## Abstract

The continued emergence of betacoronaviruses underscores the urgent need for vaccines that provide broadly protective immunity. Here, we present an epitope-focused vaccine strategy targeting the conserved S2 stem-helix region of the spike fusion machinery, a broadly neutralizing antibody-(bnAb) epitope shared across betacoronaviruses yet partially occluded on the native spike. Immunization of non-human primates with engineered S2 stem-helix nanoparticle immunogens, alone or followed by a SARS-CoV-2 BA.1 spike mRNA boost, elicited broadly cross-reactive antibody responses against sarbecoviruses, merbecoviruses, and embecoviruses and neutralized SARS-CoV-2, multiple variants, other sarbecoviruses, and MERS-CoV. Vaccine-elicited monoclonal antibodies displayed broad *in-vitro* neutralizing activity and protected against both SARS-CoV-2 and MERS-CoV *in-vivo*. Structural analyses revealed conserved features between rhesus and human stem-helix bnAbs, supporting the translational potential. Overall, our findings provide proof-of-concept that epitope-focused nanoparticle immunogens can target partially occluded, immunoquiescent bnAb epitopes, laying the groundwork for pan-betacoronavirus vaccines that provide broad protection and strengthen pandemic preparedness.

**ONE SENTENCE SUMMARY:** Epitope-focused S2 stem-helix nanoparticle immunogens elicit protective broadly neutralizing antibodies (bnAbs) against diverse betacoronaviruses in non-human primates, establishing a framework for development of pan-betacoronavirus vaccines.

## Introduction

The persistent threat posed by zoonotic spillovers of betacoronaviruses, exemplified by SARS-CoV-1 in 2002, MERS-CoV in 2012, and SARS-CoV-2 in 2019 (*1–4*), underscores the urgent need for next-generation vaccines capable of providing broad protection across diverse lineages (*5–7*). Current vaccines, although highly effective against specific strains of each betacoronavirus, largely rely on inducing neutralizing antibodies (nAbs) against the receptor-binding domain (RBD) of the spike protein (*8–12*). However, the RBD shows considerable sequence variation among betacoronaviruses. Furthermore, for a given betacoronavirus, most nAb sites on the RBD can tolerate substitutions with limited fitness costs thereby permitting immune escape by emerging variants (*13–15*). One approach to pan-betacoronavirus vaccine design has been to generate immunogens incorporating RBDs from multiple betacoronaviruses (*16–19*), but the breadth of such mosaic strategies is intrinsically constrained by the specific RBDs represented in the formulation of the immunogen. To overcome this limitation, an alternative strategy is to design immunogens that elicit broadly neutralizing antibodies (bnAbs) against conserved and functionally constrained epitopes on the viral spike.

The S2 subunit of the betacoronavirus spike protein harbors such conserved and functionally constrained epitopes (*20–25*). In particular, the S2 stem-helix region within the spike fusion machinery has emerged as a promising target of protective bnAbs capable of cross-reacting with diverse betacoronaviruses, including sarbecoviruses and merbecoviruses (*25–32*). However, structural and immunological studies reveal that this epitope region is largely buried in both pre-and post-fusion spike conformations, rendering it poorly accessible and immunologically subdominant during natural infection or conventional vaccination (*25, 27, 29*). Although recent vaccine design efforts have sought to redirect B cell responses to this site (*21, 26, 28, 33–35*), none have consistently and reliably induced cross-betacoronavirus bnAbs. These challenges underscore the need for precision vaccine strategies designed to better expose the stem-helix epitope and focus immune responses on this conserved yet partially occluded site of vulnerability.

In this study, we present an epitope-focused vaccine strategy targeting the spike S2 stem-helix bnAb site. We show that immunization of non-human primates with engineered nanoparticle immunogens displaying S2 stem-helix bnAb site, administered alone or combined with SARS-CoV-2 BA.1 spike mRNA vaccine boost, elicited robust antibody responses that neutralized SARS-CoV-2 (including Omicron variants), diverse sarbecoviruses, and MERS-CoV. Monoclonal antibodies isolated from vaccinated animals conferred *in vivo* protection against both SARS-CoV-2 and MERS-CoV. High-resolution structural analyses of the rhesus bnAbs revealed convergence in structure and paratope with human S2 stem-helix bnAbs, supporting the translational potential of this vaccine approach. Together, these findings establish a proof-of-concept for pan-betacoronavirus vaccine design based on epitope-focused nanoparticle immunogens and highlight the broader potential of this approach for targeting partially occluded, immunoquiescent conserved bnAb sites on glycoproteins of highly immune-evasive RNA viruses.

## Results

### Design of S2 stem-helix bnAb site-targeted nanoparticle immunogens

The spike stem-helix region, which is critical to the viral fusion machinery, has emerged as a promising target for broad betacoronavirus vaccine strategies. Our recent work demonstrated that bnAbs directed against this region confer protection against multiple pathogenic betacoronaviruses, including SARS-CoV-1, SARS-CoV-2, and MERS-CoV (*25*). However, natural infection or conventional vaccination rarely elicit such bnAbs, likely because the stem-helix epitope site remains poorly accessible on the native spike (*25, 27, 29*). To overcome this challenge, we adopted an epitope-focused vaccine design approach to present the stem-helix bnAb epitope region in a highly accessible and more immunogenic context, thereby enhancing its ability to engage and stimulate B cell responses to this otherwise immunologically subdominant region.

A 25-amino acid or 27-amino acid (MERS-CoV) peptide encompassing the S2 stem-helix region of betacoronavirus spike proteins from multiple subgenera, including SARS-CoV-2 from sarbecovirus, MERS-CoV from merbecovirus, as well as HCoV-OC43, HCoV-HKU1, and murine hepatitis virus (MHV) from embecovirus, was grafted onto a self-assembling ferritin nanoparticle derived from *Helicobacter pylori* (Fig. 1A-B, S1A) (*36–38*). This 25 or 27-AA peptide region contains the complete epitopes of most reported S2 stem-helix bnAbs (*27*) and, since ferritin assembles into a 24-mer nanoparticle, it supports multivalent display of the conserved bnAb epitope on each nanoparticle (Stem-FR; Fig. 1B, S1A). To further increase epitope density and enhance immunogenicity, we engineered variants in which each ferritin subunit displays three tandem copies (3R) of the S2 stem-helix peptide, separated by flexible glycine-serine (GS) linkers (Stem-3R-FR), resulting in presentation of 72 copies of the stem-helix peptide epitope per nanoparticle (Fig. 1B, S1B). In parallel, we designed a mosaic nanoparticle in which each ferritin subunit displays three tandem stem-helix peptides derived from SARS-CoV-2 (sarbecovirus), MERS-CoV (merbecovirus), and HCoV-OC43 (embecovirus), respectively (Mosaic-Stem-3-FR; Fig. 1B, S1B). This design presents 24 copies of each subgenus-specific stem-helix region, for a total of 72 copies per nanoparticle, while maintaining equal representation from the three distinct betacoronavirus subgenera.

**Figure 1.**
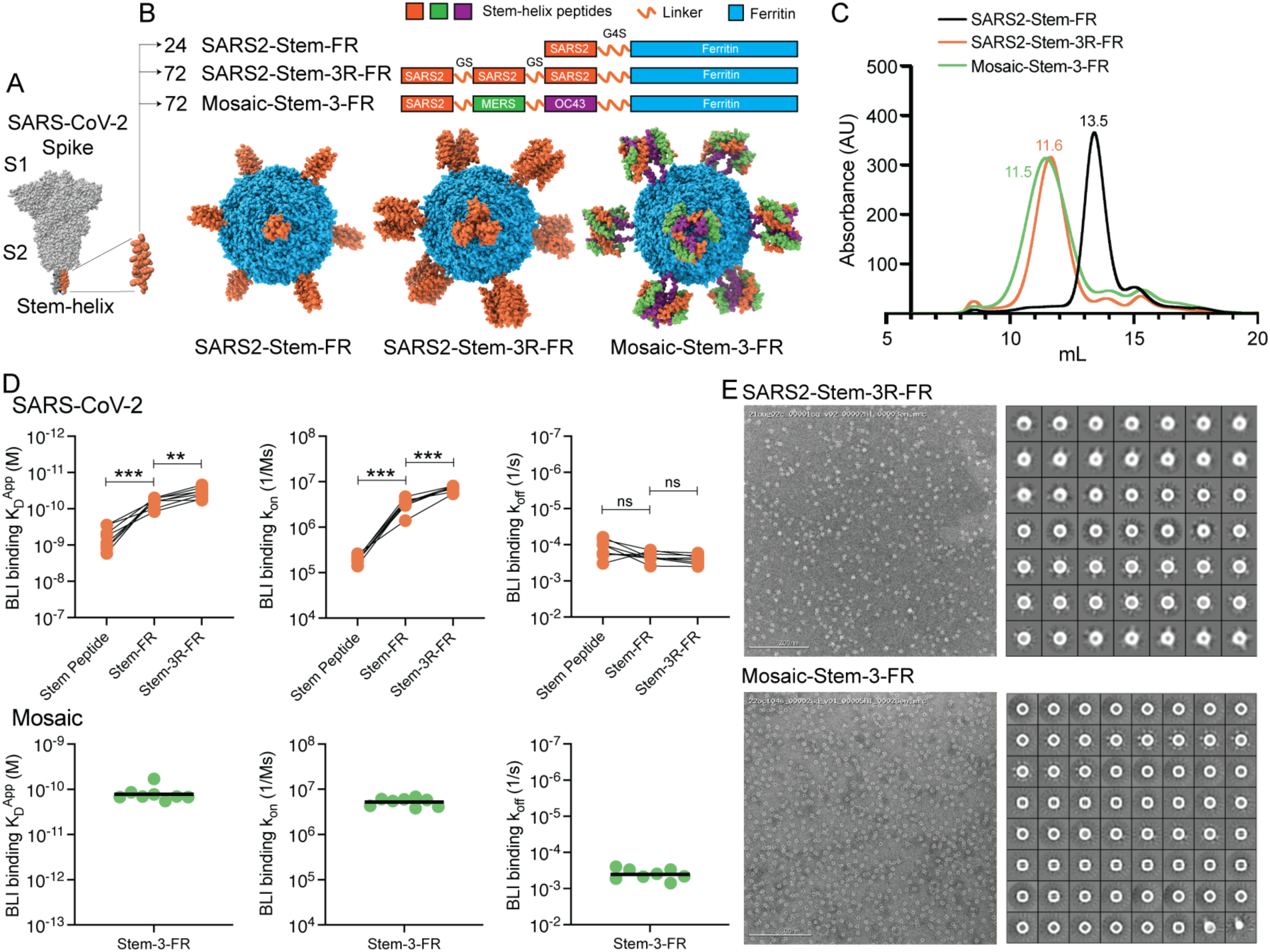
Design and characterization of S2 stem-helix ferritin nanoparticles. (**A**) Architecture of the SARS-CoV-2 spike protein showing the S1 and S2 subunits. The conserved S2 stem-helix region at the base of the spike is highlighted in orange. (**B**) Schematics and structural models of ferritin-based stem-helix nanoparticles. Each ferritin subunit was fused at its N-terminus to either one (SARS2-Stem-FR) or three tandem repeats (SARS2-Stem-3R-FR) of the 25-amino-acid SARS-CoV-2 S2 stem-helix peptide via G4S linkers. SARS2-Stem-FR displayed 24 copies of the peptide per nanoparticle, SARS2-Stem-3R-FR displayed 72 copies, and Mosaic-Stem-3-FR presented 24 copies each of SARS-CoV-2 (orange), MERS-CoV (green), and HCoV-OC43 (purple) stem-helix peptides on the ferritin scaffold (blue). For visualization, the structural models depict the stem-helix peptides in a trimeric arrangement; however, on the nanoparticles the peptides are expected to be displayed as individual monomers rather than trimers. (**C**) Size-exclusion chromatography of SARS2-Stem-FR (black), SARS2-Stem-3R-FR (orange), and Mosaic-Stem-3-FR (green) on a Superose 6 Increase 10/300 GL column. Absorbance monitored at 280nm. (**D**) BioLayer Interferometry analysis of broadly neutralizing S2 stem-helix antibody binding to monomeric peptide, SARS2-Stem-FR, SARS2-Stem-3R-FR, and Mosaic-Stem-3-FR. Apparent dissociation constants (K_D_^App^), association rates (k_on_), and dissociation rates (k_off_) are shown. For Mosaic-Stem-3-FR, horizontal bars represent geometric mean values. (**E**) Representative negative-stain electron micrographs (left) and corresponding 2D class averages (right) of SARS2-Stem-3R-FR and Mosaic-Stem-3-FR. Scale bars, 200 nm. *P* values were calculated by a Mann-Whitney test. ns, not significant; **p < 0.01; ***p < 0.001.

The nanoparticles were expressed in HEK293F mammalian cells and purified by *Galanthus nivalis* lectin (GNL) affinity chromatography, taking advantage of conserved glycosylation sites present on both the S2 stem-helix peptide and the ferritin scaffold (Fig. S1A-C). With the exception of MERS-Stem-3R-FR and OC43-Stem-3R-FR, size-exclusion chromatography (SEC) profiles of the stem-helix nanoparticle immunogens exhibited single, homogeneous peaks consistent with the predicted nanoparticle sizes (Fig. 1C, S1D-E), indicating efficient self-assembly of most constructs.

The antigenicity of the assembled nanoparticles was evaluated using a panel of S2 stem-helix bnAbs. All successfully assembled stem-helix nanoparticles displayed the desired antigenic properties, demonstrating that this epitope-focused nanoparticle design strategy can be broadly applied across diverse betacoronavirus stem-helix sequences (Fig. 1D, S2A-B). Multivalent presentation of the stem-helix peptide substantially enhanced apparent binding affinities to multiple stem-helix binding antibodies compared with the corresponding synthetic monomeric peptides (Fig. 1D, S2C). For example, SARS2-Stem-FR bound S2 stem-helix antibodies with apparent equilibrium dissociation constants (K_D_^app^) in the 10^-10^–10^-11^ M range, compared with 10^-9^–10^-10^ M for the synthetic monomeric SARS-CoV-2 stem-helix peptide. The higher-valency SARS2-Stem-3R-FR design further increased apparent affinity, primarily by enhancing association rates (k_on_) while maintaining similar dissociation rates (k_off_) (Fig. 1D). Notably, the Mosaic-Stem-3-FR nanoparticle exhibited comparable binding kinetics to stem-helix bnAbs, validating the overall design strategy. Based on expression yield, antigenic profile, assembly characteristics, and representation of distinct betacoronavirus subgenera, SARS2-Stem-3R-FR and Mosaic-Stem-3-FR were selected for subsequent immunogenicity and functional studies.

To validate the structural integrity and uniformity of the S2 stem-helix ferritin nanoparticles, we performed negative stain electron microscopy (ns-EM) on SARS2-Stem-3R-FR and Mosaic-Stem-3-FR constructs. Ns-EM analysis revealed uniformly sized, well-folded, self-assembling nanoparticles, consistent with multivalent epitope display and structural homogeneity (Fig. 1E). Differential scanning calorimetry (DSC) demonstrated excellent thermal stability, with melting temperatures (T_m_) of approximately 81.0±0.8°C for SARS2-Stem-3R-FR and 81.6±0.2°C for Mosaic-Stem-3-FR, indicating robust structural stability suitable for vaccine applications (Fig. S1F). Site-specific glycan analysis by mass spectrometry revealed substantial glycosylation site occupancy across ferritin protomers, with predominantly complex-type glycans consistent with proper mammalian glycan processing (Fig. S1C).

Together, these comprehensive biophysical characterizations established that the S2 stem-helix nanoparticles possessed the structural uniformity, thermal stability, and desired antigenic profile, supporting their development as pan-betacoronavirus vaccine candidates.

### Immunogenicity evaluation of S2 stem-helix nanoparticle in rhesus macaques

Rhesus macaques (RMs) share key B cell immunogenetic features and affinity maturation pathways with humans, making them a valuable outbred preclinical model for predicting vaccine responses. Although we previously showed that RMs are not an ideal model for evaluating SARS-CoV-2 RBD-based vaccines because their antibody repertoires are intrinsically predisposed to generate cross-neutralizing RBD bnAbs through germline-encoded features (*39*), our recent findings demonstrate that they are well suited for evaluating S2 stem-helix–targeted vaccine strategies. Specifically, vaccine-elicited rhesus S2 stem-helix bnAbs closely resemble the public clonotype S2 stem-helix bnAbs identified in humans (*28*). To evaluate the immunogenicity of the designed S2 stem-helix nanoparticles, two RM cohorts (n = 6 per group) were subcutaneously immunized with 100 µg of nanoparticle protein formulated with saponin/MPLA nanoparticle adjuvant (SMNP) (*40*). Group 1 received SARS2-Stem-3R-FR, and Group 2 received Mosaic-Stem-3-FR (Fig. 2A). Both groups received homologous nanoparticle boosts (B1) at week 4, followed by a heterologous boost (B2) at week 12 with 100 µg of SARS-CoV-2 Omicron BA.1 spike mRNA–lipid nanoparticles (LNPs) to recall and further mature S2 stem-helix–specific B cell responses. Plasma and peripheral blood mononuclear cells (PBMCs) were collected longitudinally (plasma: weeks -1, 2, 6, 12, and 14; PBMCs: weeks 6 and 14) to monitor antibody and B cell responses (Fig. 2A).

**Figure 2.**
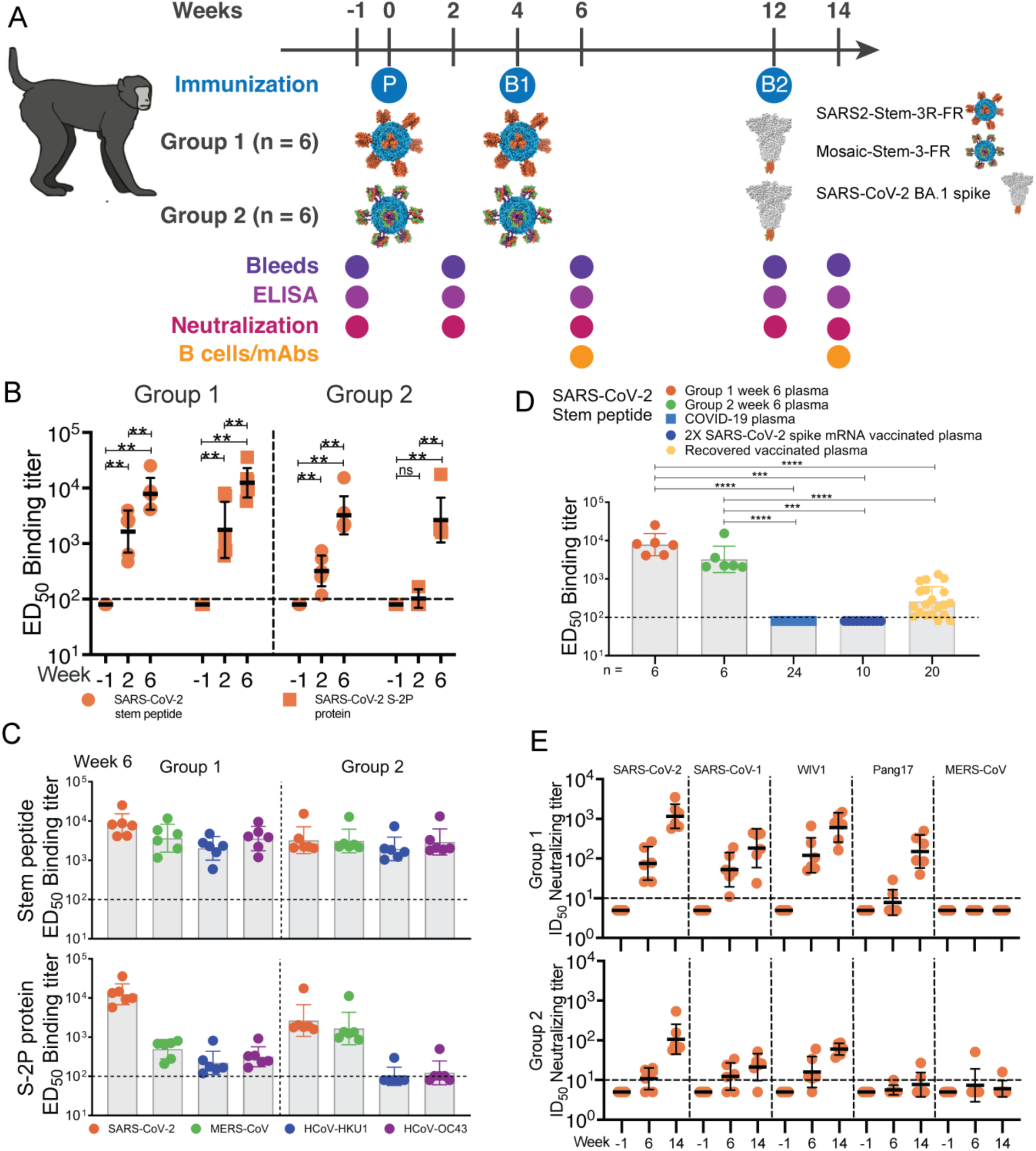
Immunogenicity evaluation of S2 stem-helix nanoparticles in rhesus macaques. (**A**) Immunization regimen of S2 stem-helix nanoparticles in rhesus macaques. Two cohorts of six rhesus macaques each received 100 µg of SARS2-Stem-3R-FR (Group 1) or Mosaic-Stem-3-FR (Group 2) formulated with 350 µg SMNP adjuvant. Vaccinations were administered subcutaneously at week 0 (prime, P) and week 4 (homologous boost, B1), followed by a 100 µg intramuscular boost (B2) with Omicron BA.1 spike mRNA-LNP at week 12. Plasma samples were collected at weeks −1, 2, 6, 12, and 14, and PBMCs at weeks 6 and 14. (**B**) ED_50_ ELISA binding titers of plasma at weeks −1, 2 and 6 against the SARS-CoV-2 S2 stem-helix peptide and S-2P spike protein. (**C**) ED_50_ binding titers of plasma at week 6 to the stem-helix peptides and S-2P spike proteins from SARS-CoV-2, MERS-CoV, HCoV-HKU1, and HCoV-OC43. (**D**) S2 stem-helix–specific ED_50_ binding titers in vaccinated macaques (week 6), COVID-19 convalescent donors (n = 24), two-dose spike mRNA vaccine recipients (n = 10), and recovered-vaccinated individuals (n = 20). (**E**) Neutralization breadth and potency of plasma at weeks −1, 6 and 14 against diverse betacoronavirus pseudoviruses, including SARS-CoV-2, SARS-CoV-1, WIV1, Pang17, and MERS-CoV. Geometric mean values ± SD wre shown in panels B–E. ED_50_, half-maximal effective dilution; ID_50_, 50% inhibitory dilution; ED_50_ values <100 and ID_50_ values <10, below the dashed line, were considered negative for binding and neutralization, respectively. ED_50_ values <100 and ID_50_ values <10 were assigned a value of 80 and 5 for visualization, respectively. *P* values were calculated by a Mann-Whitney test. ns, not significant; \*\**P* < 0.01; \*\*\**P* < 0.001; \*\*\*\**P* < 0.0001.

Both immunization groups developed robust antibody responses against SARS-CoV-2 S2 stem-helix peptide by two weeks after the prime, as measured by ELISA binding (Fig. 2B, S3). Antibody titers increased significantly following the homologous boost, with peak responses observed at week 6. Group 1 achieved higher geometric mean ED_50_ binding titers compared to Group 2, although all animals in both cohorts mounted consistent S2-directed antibody responses. In addition to recognizing the SARS-CoV-2 stem-helix peptide, plasma collected at weeks 2 and 6 from both groups exhibited broad cross-reactive binding to the corresponding stem-helix peptides from other human betacoronaviruses, including MERS-CoV, HCoV-HKU1, and HCoV-OC43 (Fig. 2C, S3). In contrast, plasma from 24 SARS-CoV-2 convalescent individuals and 10 SARS-CoV-2 spike mRNA vaccine recipients exhibited no detectable binding to the stem-helix peptides from SARS-CoV-2 and other human betacoronaviruses tested (Fig. 2D, S4), highlighting the immunologically subdominant nature of this conserved bnAb epitope in natural infection or conventional vaccination. Consistent with previous reports (*41–45*), plasma from 20 individuals with hybrid immunity (infection plus vaccination; “recovered vaccinated”) exhibited detectable S2 stem-helix reactivity; however, these responses remained significantly weaker than those elicited by our epitope-focused nanoparticle immunization strategy (Fig. 2D, S3, S4). Together, these findings underscored the potent immunofocusing capacity of our epitope-focused nanoparticle approach in directing antibody responses toward a highly conserved yet naturally subdominant bnAb site.

**Figure 3.**
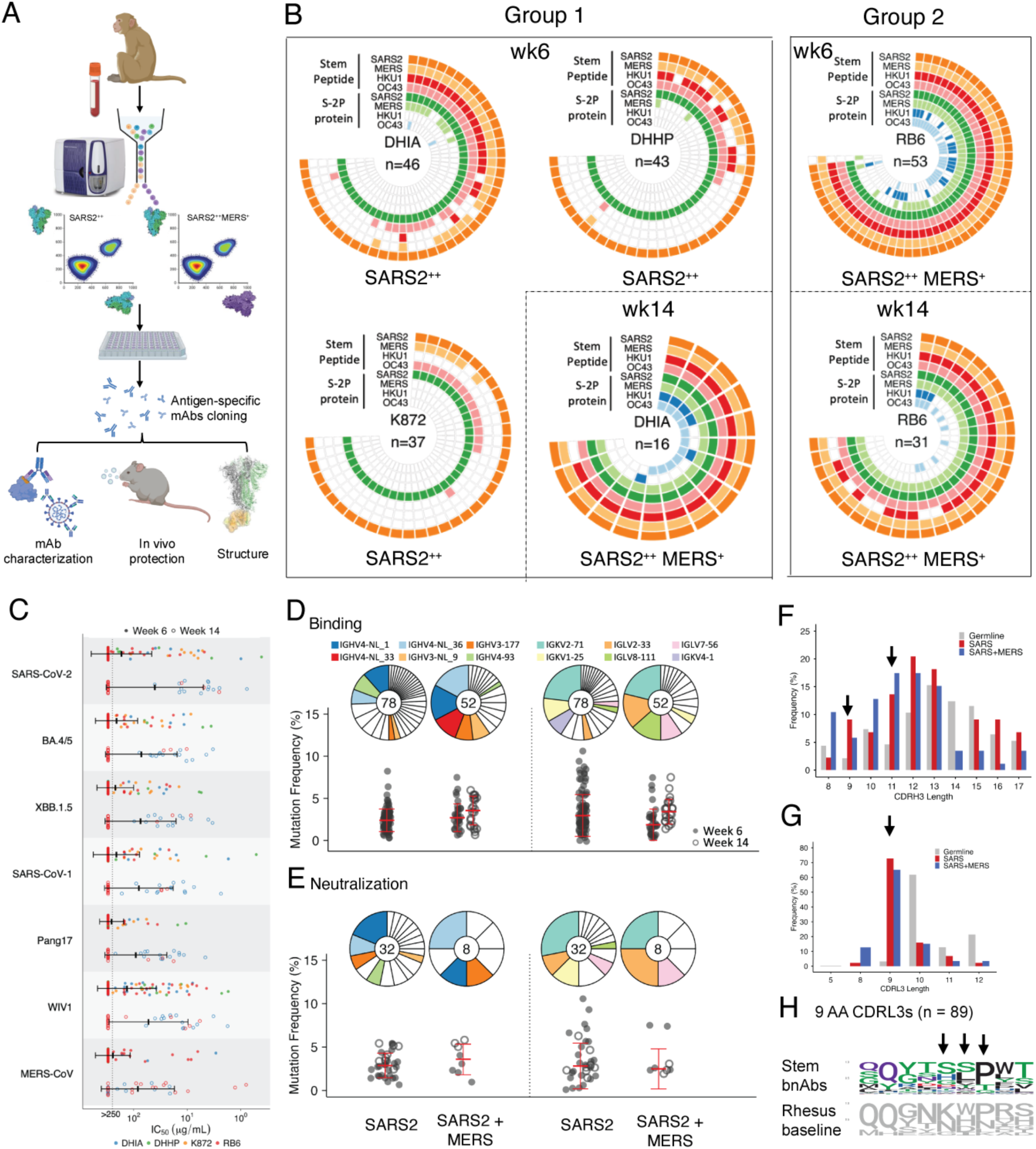
Isolation and characterization of antibodies from S2 stem-helix nanoparticle–vaccinated rhesus macaques. (**A**) Schematic overview of the methods used to isolate antigen-specific, IgG class-switched B cells from PBMCs of vaccinated macaques. SARS2^++^: B cells sorted using SARS-CoV-2 S-2P protein alone; SARS2^++^MERS^+^: dual-positive B cells sorted using both SARS-CoV-2 and MERS-CoV S-2P proteins. (**B**) Binding specificity of monoclonal antibodies (mAbs) isolated from Group 1 animals (DHIA, DHHP, K872) and the Group 2 representative (RB6). Each mAb was represented as a radial vector, with colored antigen rings indicating binding to specific antigens (stem peptide: SARS2, orange; MERS, light orange; HKU1, red; OC43, pink; S-2P protein: SARS2, green; MERS, light green; HKU1, blue; OC43, light blue). Antibodies are ordered clockwise by decreasing average OD_405nm_ binding values across all tested antigens (also shown in Table S1). OD_405 nm_ values < 0.8 were considered negative for binding, and corresponding radial vectors were shown in white. SARS2, SARS-CoV-2; MERS, MERS-CoV; HKU1, HCoV-HKU1; and OC43, HCoV-OC43. (**C**) Neutralization activity of S2 stem-helix mAbs from DHIA (blue), DHHP (green), K872 (orange), and RB6 (red) after homologous nanoparticle boost (week 6, closed dots) and SARS-CoV-2 BA.1 spike boost (week 14, open dots) immunizations against SARS-CoV-2, its Omicron variants BA.4/5 and XBB.1.5, SARS-CoV-1, Pang17, WIV1 and MERS-CoV. Geometric mean IC_50_ values ± SD are shown. IC_50_ values >250 μg/mL were considered not-neutralizing, and . IC_50_ values >250 μg/mL were assigned a value of 300 for visualization. IC_50_, 50% inhibitory concentration. (**D-E**) Germline gene usage distribution and somatic hypermutation level (mutation frequency) for isolated S2 stem-helix mAbs grouped by binding (**D**) and neutralization specificity (**E**) after homologous nanoparticle boost (week 6, closed dots) and SARS-CoV-2 BA.1 spike boost (week 14, open dots). Dot plots show the percentage of nucleotide somatic hypermutations (SHMs) in the heavy and light chain variable regions. Median values ± SD are shown. Pie chart segments are arranged counterclockwise by V gene usage frequency, from highest to lowest. SARS2: SARS-CoV-2-specific binders or neutralizers, binding to or neutralizing SARS-CoV-2 but not MERS-CoV; SARS2+MERS: cross-binding or cross-neutralizing antibodies to both SARS-CoV-2 and MERS-CoV. (**F-G**) CDRH3 loop (**F**) and CDRL3 loop (**G**) length distributions of isolated S2 stem-helix mAbs binding SARS-CoV-2 spike specifically (SARS2, red, n = 78) or cross-reactive to SARS-CoV-2 and MERS-CoV spike (SARS2+MERS, blue, n = 52) compared to rhesus baseline germline reference (grey). (**H**) Amino acid enrichments of 9 AA-long CDRL3-bearing stem-helix mAbs (n = 89) compared to rhesus baseline germline reference. Enriched residues (corresponding to an SSP motif) are indicated by arrows. The 130 unique mAbs shown in panels D, F, and G are selected from the 131 antibody lineages shown in Table S1 (one mAb per lineage). RB6-Stem10.1 is excluded from this analysis because it only displays binding to the MERS-CoV S-2P protein and the stem-helix peptide. Panels E and H present analyses of selected subsets of these 130 unique mAbs.

**Figure 4.**
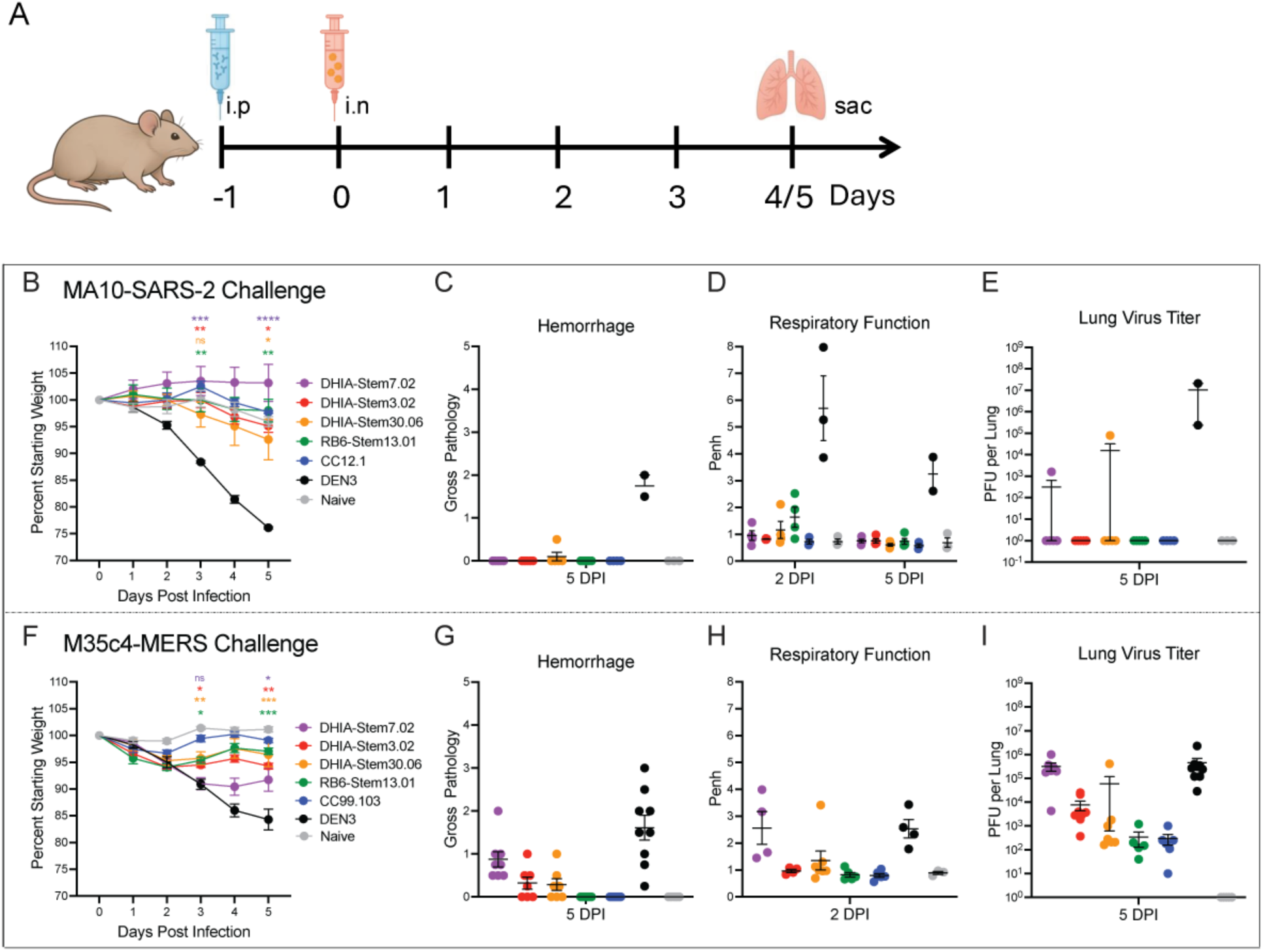
Prophylactic efficacy of rhesus S2 stem-helix bnAbs against SARS-CoV-2 and MERS-CoV challenge in aged mice. (**A**) Experimental scheme for prophylaxis of rhesus S2 stem-helix bnAbs against SARS-CoV-2 and MERS-CoV challenge in aged mice. Four rhesus S2 stem-helix bnAbs (DHIA-Stem7.02, DHIA-Stem3.02, DHIA-Stem30.06 and RB6-Stem13.01) along with SARS-CoV-2 human RBD-targeting nAb CC12.1, human S2 stem-helix bnAb CC99.103 and a negative control antibody DEN3, were administered intraperitoneally (i.p.) to groups of aged mice at a dose of 300μg per mouse. Twelve hours post-infusion, mice were challenged intranasally (i.n.) with either mouse-adapted SARS-CoV-2 (MA10) or MERS-CoV (M35c4). (**B, F**) Percent weight change following viral challenge in rhesus bnAbs (DHIA-Stem7.02, DHIA-Stem3.02, DHIA-Stem30.06 and RB6-Stem13.01), CC12.1, CC99.103 or DEN3-treated animals. Percent weight change was calculated from day 0 starting weight for all animals. Weight loss was monitored daily and expressed as arithmetic mean ± SEM. As a control, mice in naive group were exposed to PBS in the absence of virus. Statistical significance was calculated with Dunnett’s multiple comparisons test between each experimental group and the DEN3 control Ab group. *p < 0.05; **p < 0.01; ***p < 0.001; ****p < 0.0001; ns, not significant (p > 0.05). A one-way ANOVA was used. (**C, G**) Gross lung hemorrhage scores at day 5 post-infection assessed at necropsy in mice from naive groups and the groups treated with rhesus bnAb, CC12.1, CC99.103, or DEN3. Data are shown as arithmetic mean values ± SEM. (**D, H**) Pulmonary function (Penh score) at day 2 or 5 post-infection measured by whole-body plethysmography in rhesus bnAb-, CC12.1-, CC99.103- or DEN3-treated mice and mice in naive group. Data are shown as arithmetic mean values ± SEM. (**E, I**) Lung viral titers (PFU per lung) determined by plaque assay at day 5 post-infection in all groups.

Beyond binding to S2 stem-helix peptides, plasma collected at weeks 2 and 6 also recognized recombinant prefusion-stabilized spike proteins (S-2P; containing two proline substitutions) from all five human betacoronaviruses. Group 2 exhibited stronger binding to the MERS-CoV spike, whereas Group 1 demonstrated broader cross-reactive binding across heterologous betacoronavirus spikes (Fig. 2C, S3). These binding profiles indicate that the nanoparticle immunogens successfully elicited S2 stem-helix–focused antibody responses capable of recognizing this conserved epitope in the context of the native-like trimeric spike glycoprotein. Consistent with these binding data, post-prime plasma from both groups neutralized SARS-CoV-2 and related sarbecoviruses as early as two weeks after the initial immunization (Fig. 2E, S5).

Neutralizing activity against most sarbecovirus pseudoviruses increased in magnitude and became more consistent across animals following the homologous boost, with peak responses observed at week 6. Group 1 elicited more potent and broadly consistent sarbecovirus neutralizing activity than Group 2, whereas MERS-CoV neutralization remained minimal in both groups and was detected in only a single animal from Group 2 (Fig. 2E, S5).

To recall and further mature S2-directed memory responses, both groups received an mRNA-LNP boost encoding the native full-length SARS-CoV-2 Omicron BA.1 spike with a transmembrane domain at week 12. Unlike most licensed vaccines, our BA.1 spike mRNA construct omitted the S2-stabilizing 2P mutations based on prior evidence that these substitutions reduced stem-helix epitope accessibility and that S2 stem-helix bnAbs bound more efficiently to membrane-anchored native spike than to stabilized soluble spike (*27*). As expected, the heterologous BA.1 boost markedly enhanced cross-reactive S2 stem-helix binding and elicited potent, broadly neutralizing serum antibody responses compared with pre-boost levels at week 12, demonstrating efficient recall of S2-focused B cell memory (Fig. S3, S5). The increased breadth and potency were unlikely to be driven primarily by antibodies targeting other spike epitopes, such as the BA.1 RBD, because Group 2 sera remained substantially less potent than Group 1 after the BA.1 boost, particularly against BA.4/5, XBB.1.5, and Pang17. Together, these results indicate that boosting promoted affinity maturation toward the native stem-helix epitope and enhanced recognition of authentic spike proteins.

Overall, these findings represented, to our knowledge, the first successful demonstration of serum bnAb responses induced by epitope-focused nanoparticle immunogens targeting a conserved yet largely occluded epitope on a class I viral fusion glycoprotein. This work established the transformative potential of rational vaccine design for redirecting immune responses toward immunologically subdominant yet highly sought-after sites on viruses that remained refractory to conventional vaccination approaches.

### Monoclonal antibodies isolated from memory B cells of S2 stem-helix nanoparticle immunized macaques

#### Broad betacoronavirus spike binding

To define the nature of the antibodies elicited by S2 stem-helix nanoparticle vaccination, we used flow cytometry to isolate antigen-specific, IgG class-switched B cells from PBMCs. Based on plasma neutralization profiles (Fig. S5), we selected three Group 1 animals (DHIA, DHHP, K872) vaccinated with SARS2-Stem-3R-FR and one Group 2 animal (RB6) vaccinated with Mosaic-Stem-3-FR for single B cell isolation. Antigen-specific B cells were isolated from all four animals at week 6 (two weeks after the homologous nanoparticle boost) and from one representative animal per group at week 14 (two weeks after the BA.1 spike mRNA boost), using SARS-CoV-2 S-2P protein (SARS2^++^) alone or in combination with MERS-CoV S-2P protein (SARS2^++^MERS^+^) as sorting baits (Fig. 3A). Heavy and light chains were amplified and sequenced from each single antigen-sorted B cells.

In total, we isolated 226 S2 stem-helix mAbs across both time points from the four animals. Week 6 yielded 179 mAbs: 46 from DHIA, 43 from DHHP, and 37 from K872 (Group 1), plus 53 from RB6 (Group 2). An additional 47 mAbs were obtained at week 14: 16 from DHIA and 31 from RB6 (Fig. 3B and Table S1). Among these antibodies, 131 were encoded by unique immunoglobulin germline VDJ gene combinations, with several forming expanded lineages containing two or more clonal members.

At week 6, all mAbs (126/126, OD_405nm_ > 0.8) from Group 1 macaques (SARS2-Stem-3R-FR–vaccinated) exhibited binding to the SARS-CoV-2 S2 stem-helix peptide, with more than half (70/126) also cross-reacting with the stem-helix peptides from the other human betacoronaviruses tested (Fig. 3B and Table S1). A subset of DHIA-derived mAbs (21/46), and to a lesser extent DHHP- and K872-derived (7/43 and 0/37, respectively), simultaneously bound SARS-CoV-2, MERS-CoV, HCoV-OC43 and HCoV-HKU1 S2 stem-helix peptides. All Group 1 mAbs at week 6 recognized the SARS-CoV-2 spike, yet most showed minimal cross-binding to other human betacoronavirus spikes, except for a few DHIA-derived mAbs (9/46) that engaged the MERS-CoV spike (Fig. 3B and Table S1), suggesting that SARS2-Stem-3R-FR primarily elicits stem-helix-specific B cells that required further maturation to efficiently recognize diverse spikes.

In contrast, mAbs isolated at week 6 from the single Group 2 macaque (RB6; Mosaic-Stem-3-FR–vaccinated) displayed substantially broader reactivity. Nearly all mAbs (52/53) cross-reacted with stem-helix peptides from all four human betacoronaviruses tested, while most also bound both SARS-CoV-2 and MERS-CoV spike proteins (39/53). In addition, a majority (34/53) recognized either the HCoV-HKU1 or HCoV-OC43 spike proteins (Fig. 3B and Table S1). This broader specificity likely reflected the dual-antigen B-cell sorting strategy, which used both SARS-CoV-2 and MERS-CoV S-2P proteins to enrich for cross-reactive memory B cells from the RB6.

Collectively, these results confirmed that epitope-focused stem-helix nanoparticle immunogens, unlike natural infection or conventional spike immunization, directed B cell responses toward the S2 stem-helix epitope and generated diverse lineages of specific mAbs with broad betacoronavirus binding potential.

#### Broad betacoronavirus neutralization

We next evaluated the neutralization breadth of the purified rhesus S2 stem-helix mAbs, representing 122 unique antibody lineages, against a panel of betacoronaviruses that included clade 1a sarbecoviruses (SARS-CoV-1 and WIV1), clade 1b sarbecoviruses (SARS-CoV-2, Omicron BA.4/5 and XBB.1.5 variants, and Pang17), and the merbecovirus MERS-CoV (Fig. 3C and Table S2). Overall, 40 of the 122 unique antibody lineages (32.8%) neutralized SARS-CoV-2, with each macaque yielding at least five distinct SARS-CoV-2-neutralizing lineages, demonstrating efficient recruitment and expansion of diverse S2 stem-helix bnAb precursor B cells (Table S2). More than half of these SARS-CoV-2-neutralizing lineages (23/40) also neutralized all of the other sarbecoviruses and SARS-CoV-2 variants tested, highlighting their broad sarbecovirus activity.

Antibodies isolated at week 14 were generally more potent than those recovered at week 6, especially within the same mAb lineages recovered at both time points (Fig. 3C and Table S2), consistent with ongoing affinity maturation following immunization. This maturation was further demonstrated in macaque RB6 (Group 2), from which both week 6 and week 14 antibodies were recovered from SARS-CoV-2^++^MERS-CoV^+^ cross-reactive B cells, the geometric mean IC_50_ values of RB6 neutralizing antibodies improved from 55.2, 52.7, 60.0, 67.1, 30.8, and 44.0 μg/mL at week 6 to 5.4, 17.8, 18.0, 27.8, 17.3, and 30.2 μg/mL at week 14 against SARS-CoV-2, BA.4/5, XBB.1.5, SARS-CoV-1, WIV1, and Pang17, respectively. Consistent with the detectable MERS-CoV plasma neutralizing activity observed in RB6 (Fig. S5), nearly half of its antibody lineages (18/44) also neutralized MERS-CoV (Table S2), including one mAb, RB6-Stem13.01, which cross-neutralized MERS-CoV and all tested sarbecoviruses.

In contrast, although macaque DHIA (Group 1) exhibited no detectable MERS-CoV serum neutralization, several isolated mAb lineages (5/35) potently neutralized MERS-CoV. In particular, DHIA-Stem7.02, isolated at week 6 following immunization with SARS2-Stem-3R-FR nanoparticle alone, displayed potent neutralizing activity against both sarbecoviruses and MERS-CoV (Table S2), demonstrating that broadly neutralizing B-cell lineages can be present even when their activity was not apparent at the serum level.

Collectively, these findings demonstrate that our stem-helix nanoparticle immunogens efficiently recruited and expanded broadly cross-reactive memory B-cell lineages capable of pan-betacoronavirus neutralization. However, these lineages often remained subdominant within the overall serum antibody response, presumably by failing to differentiate efficiently into antibody-secreting plasma cells and thereby limiting their contribution to serum neutralizing activity. These results therefore highlighted the need for further optimization of immunogen design and immunization strategies to preferentially amplify these rare broadly neutralizing B-cell lineages and achieve consistent pan-betacoronavirus serum neutralization.

#### Immunogenetic properties

To investigate whether antibody genetic features were associated with binding breadth, we analyzed variable gene usage and somatic hypermutation (SHM) in 130 unique mAbs using IgDiscover (*46*). Antibodies were grouped according to their binding specificity as either SARS-CoV-2 spike-specific (binding to SARS-CoV-2 but not MERS-CoV spike, n = 78) or cross-reactive with both SARS-CoV-2 and MERS-CoV spikes (n = 52) (Fig. 3D). Both groups exhibited preferential usage of specific IGHV and IGLV germline gene families. Among heavy-chain genes, IGHV4-NL_1 (11.5% vs. 15.4%; SARS-CoV-2-specific vs. SARS-CoV-2/MERS-CoV cross-reactive) and IGHV4-NL_36 (6.4% vs. 17.3%) were the most frequently utilized (Fig. 3D and Fig. S6A). Light-chain repertoires were similarly dominated by IGKV2-71 (23.1% vs. 21.2%) (Fig. 3D and Fig. S6A). Similar V-gene usage patterns were observed when antibodies were grouped according to stem-helix peptide binding specificity. Notably, substantially more antibodies exhibited cross-reactive binding to both SARS-CoV-2 and MERS-CoV stem-helix peptides (86/130) than to the corresponding full-length spike proteins (52/130) (Fig. 3D, S6B and Table S1), indicating that many vaccine-elicited antibodies recognize conserved stem-helix epitopes but had not yet acquired sufficient affinity or structural accommodation to engage the native spike proteins of divergent betacoronaviruses.

SARS-CoV-2 spike-specific mAbs (n = 78) exhibited relatively modest levels of SHM, with median V_H_ and V_L_ mutation frequencies of 2.39% and 2.96% at week 6, respectively (Fig. 3D and Table S1). Similarly, mAbs cross-reactive with both SARS-CoV-2 and MERS-CoV spikes also showed modest but progressive increases in SHM following BA.1 spike mRNA boost, with median V_H_ mutation frequencies increasing from 2.70% at week 6 to 3.55% at week 14, and median V_L_ mutation frequencies increasing from 1.86% to 3.41%. Comparable increases in SHM were observed for antibodies grouped by stem-helix peptide binding specificity (Fig. S6B). Together, these findings indicated that the BA.1 spike mRNA boost appeared to drive continued affinity maturation and clonal selection of vaccine-induced stem-helix-specific B-cell responses, although overall SHM levels remained relatively low, consistent with human S2 stem-helix bnAbs previously described that achieved breadth with limited mutations (*25*).

We next assessed whether immunoglobulin gene usage correlated with neutralization breadth. Antibodies were classified as either SARS-CoV-2-specific neutralizers (neutralizing SARS-CoV-2 but not MERS-CoV; *n* = 32) or cross-neutralizers against both SARS-CoV-2 and MERS-CoV (*n* = 8) (Fig. 3E). Gene usage among neutralizing mAbs closely mirrored that observed in the broader binding groups, with frequent utilization of IGHV4-NL_1 (18.8% vs. 12.5%; SARS-CoV-2-specific vs. SARS-CoV-2/MERS-CoV cross-neutralizers), IGHV4-NL_36 (9.4% vs. 25.0%), and IGKV2-71 (28.1% vs. 25.0%). Although somatic hypermutation (SHM) levels remained relatively low across all neutralizing antibodies, cross-neutralizers exhibited a modestly higher median VH mutation frequency than SARS-CoV-2-specific neutralizers (3.58% vs. 2.88%), whereas their median VL mutation frequency was slightly lower (2.48% vs. 2.82%). These findings suggest that increased VH somatic hypermutation may contribute to broader betacoronavirus neutralization.

Analysis of CDRH3 loop lengths in rhesus S2 stem-helix mAbs revealed a 3- to 4-fold enrichment of 9- and 11-residue loops, relative to the baseline macaque repertoire (Fig. 3F), closely mirroring the CDRH3 length distribution of human S2 stem-helix bnAbs (*25*). Similarly, CDRL3 loops were strongly enriched for a 9-residue length (approximately 20-fold enrichment; Fig. 3G) and frequently encoded a conserved VJ-derived “SSP” motif (Fig. 3H), resembling the “SSPPxF” motif characteristic of human S2 stem-helix bnAbs, in which the CDRL3-encoded PP residues are critical for stem-helix recognition (*25*).

Together, these findings demonstrate that rhesus S2 stem-helix mAbs are derived from a restricted set of germline V genes, require only modest affinity maturation to achieve broad neutralizing activity, and display convergent CDR3 features that parallel those of human stem-helix bnAbs associated with broad betacoronavirus recognition.

#### Epitope specificities

Among the 131 unique rhesus antibody lineages identified, we selected 29 representative antibodies for detailed characterization, including 3 stem-helix peptide broad binders, 5 MERS-CoV-only neutralizers, 17 sarbecovirus-only neutralizers, and 4 pan-betacoronavirus bnAbs. Compared with S-2P proteins, these rhesus antibodies generally exhibited stronger binding to cell surface-expressed full-length native spike proteins, particularly those from HCoV-HKU1 and HCoV-OC43 (Table S3), mirroring the preference for membrane-expressed native spike reported for human S2 stem-helix bnAbs (*25*).

To define their epitopes, we performed alanine-scanning mutagenesis of the SARS-CoV-2 stem-helix peptide in combination with antibody competition assays. These analyses identified a conserved core set of S2 stem-helix residues targeted by the rhesus antibodies (Table S4). Consistent with prior mapping of human S2 stem-helix bnAbs, substitutions at the highly conserved residues F1148, L1152, and F1156 markedly reduced antibody binding, indicating that these residues constitute key determinants of epitope recognition across betacoronaviruses. In addition, mutations at E1151 and Y1155 impaired binding for a subset of rhesus antibodies, suggesting some diversity in fine epitope specificity within the antibody set. Competition-binding analyses further demonstrated substantial epitope overlap between rhesus and human S2 stem-helix antibody: with the exception of several broad binders and MERS-CoV-only neutralizers, most rhesus antibodies effectively competed with epitope-defined human S2 stem-helix bnAbs (Table S5).

Collectively, these findings demonstrate that S2 stem-helix nanoparticle-elicited rhesus antibodies target a highly conserved epitope and shared a common mode of recognition with human S2 stem-helix bnAbs, reinforcing the translational relevance of our vaccine approach.

#### Protection against challenge with both SARS-CoV-2 and MERS-CoV

To evaluate the protective efficacy of S2 stem-helix bnAbs elicited by our nanoparticle immunogens, aged mice received prophylactic intraperitoneal injections of individual mAbs (300 µg per mouse) and were challenged 12 hours later with mouse-adapted SARS-CoV-2 (MA10) or MERS-CoV (M35c4) (Fig. 4A). Four of the most potent bnAbs, DHIA-Stem7.02 (isolated at week 6 post homologous S2 stem-helix nanoparticle boost), DHIA-Stem3.02, DHIA-Stem30.06 and RB6-Stem13.01 (all isolated at week 14 following BA.1 spike mRNA boost), were tested alongside the SARS-CoV-2 RBD antibody CC12.1, human stem-helix bnAb CC99.103 and a negative control antibody (DEN3) in groups of five-ten mice each. Animals were monitored daily for weight loss, and euthanized on day 5 for gross lung pathology and viral titer analysis. Pulmonary function was assessed via whole body plethysmography at day 2 (SARS-CoV-2 and MERS-CoV) and day 5 (SARS-CoV-2) post infection. Compared with DEN3-treated controls, rhesus bnAb-treated mice exhibited significantly reduced weight loss, minimal pulmonary hemorrhage, preserved respiratory mechanics, and substantially lower lung viral loads in both SARS-CoV-2 and MERS-CoV challenge models (Fig. 4B-I).

Taken together, these results demonstrate that S2 stem-helix nanoparticle-induced bnAbs confer robust in vivo protection against both SARS-CoV-2 and MERS-CoV, providing proof-of-concept that vaccine-elicited stem-helix bnAbs can mitigate severe betacoronavirus disease.

#### Structural basis for rhesus S2 stem-helix bnAb recognition

To provide molecular insights into the interaction of rhesus bnAbs with their S2 stem-helix targets, compare the modes of interaction of the different bnAbs, and better understand their abilities to broadly neutralize, we determined high-resolution crystal structures of eight rhesus mAb Fab-peptide complexes. The Fabs were derived from two MERS-CoV-only neutralizers (DHIA-Stem15.02 and RB6-Stem7.01), two sarbecovirus-only neutralizers (DHHP-Stem20.01 and RB6-Stem20.01), and four pan-betacoronavirus bnAbs (DHIA-Stem7.02, DHIA-Stem3.02, DHIA-Stem30.06, and RB6-Stem13.01). Crystal structures were solved with S2 stem-helix peptides from SARS-CoV-2 (six complexes at 1.15–2.28 Å resolution) and from MERS-CoV and HCoV-OC43 (1.68 Å and 2.77 Å, respectively) (Fig. 5A-D, S7A-D, and Table S6). In all eight structures, the S2 stem-helix peptides adopted a helical conformation and made extensive contacts with the antibodies. Heavy- and light-chain CDR loops formed a hydrophobic groove that accommodated the S2 stem-helix region (Fig. S8), burying 34–52% of the stem-helix surface via interactions with CDR H1, H2, H3, L1, and L3 (and all six CDRs in some antibodies) (Fig. 5A-D, S7A-E). Structural superposition of the eight antibody-S2 stem-helix peptide structures onto pre- and post-fusion SARS-CoV-2 spike protein showed steric clashes in both conformations (*47*) (Fig. S9A-H), consistent with a possible neutralization mechanism in which these antibodies bind a transient fusion-intermediate state and disrupt the formation of the six-helix bundle required for membrane fusion and viral entry (*48–51*).

**Figure 5.**
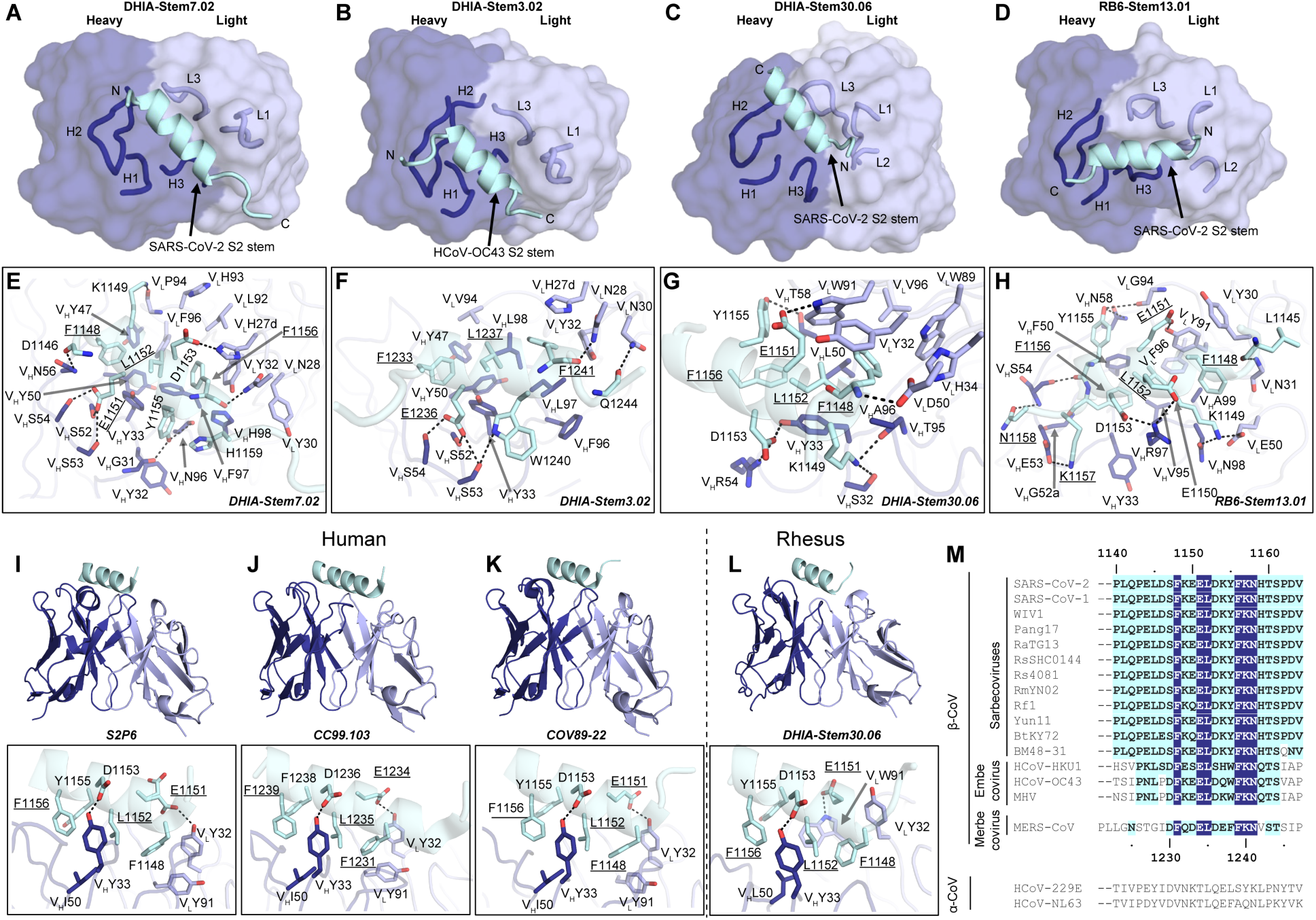
Crystal structures and detailed atomic interactions of rhesus antibodies in complex with S2 stem-helix peptides. Heavy and light chains of antibodies are shown in blue and lavender, respectively, and the S2 stem-helix peptides in cyan. Numbering of peptides is based on each corresponding betacoronavirus. Kabat numbering was applied to the antibodies. Hydrogen bonds and salt bridges are represented by black dashed lines. Conserved identical residues among betacoronaviruses are underlined. (**A-D**) Overall views of the crystal structures of DHIA-Stem7.02 mAb-SARS-CoV-2 S2 stem-helix peptide, DHIA-Stem3.02 mAb-HCoV-OC43 S2 stem-helix peptide, DHIA-Stem30.06 mAb-SARS-CoV-2 S2 stem-helix peptide, and RB6-Stem13.01 mAb-SARS-CoV-2 S2 stem-helix peptide complexes at 1.80 Å, 2.77 Å, 1.81 Å, and 1.15 Å resolutions, respectively. CDRH1, H2, H3, L1, L2, L3 are shown in backbone tube representation and labeled. (**E-H**) Detailed atomic interactions between the S2 stem-helix peptides and rhesus antibodies: (**E**) DHIA-Stem7.02 with SARS-CoV-2 S2 stem-helix peptide. There are 9 H-bonds; hydrophobic and aromatic paratope residues included V_H_ Y33, Y47, Y50, F97, H98 and V_L_ H27d, Y30, Y32, L92, F96. (**F**) DHIA-Stem3.02 with HCoV-OC43 S2 stem-helix peptide. There are 6 H-bonds; hydrophobic and aromatic paratope residues included V_H_ Y33, Y47, Y50, F96, L97, L98, and V_L_ H27d, Y32, V94. (**G**) DHIA-Stem30.06 with SARS-CoV-2 S2 stem-helix peptide. There are 6 H-bonds and 1 salt bridge; hydrophobic and aromatic paratope residues included V_H_ Y33, L50, A96, and V_L_ Y32, H34, W89, W91, V96. (**H**) RB6-Stem13.01 with SARS-CoV-2 S2 stem-helix peptide. There are 8 H-bonds and 4 salt bridges; hydrophobic and aromatic paratope residues included V_H_ Y33, F50, V95, and V_L_ Y30, Y91, F96. (**I-L**) Structural alignment of DHIA-Stem30.06 with three IGHV1-46/IGKV3-20 antibodies in complex with S2 stem-helix peptides. All of the bound peptides are in the same orientation. For clarity, only the variable domains of Fabs are shown. (**I**) human antibody S2P6 with SARS-CoV-2 S2 stem-helix peptide (PDB 7RNJ). (**J**) human antibody CC99.103 with MERS-CoV S2 stem-helix peptide (PDB 8DGV). (**K**) human antibody COV89-22 with SARS-CoV-2 S2 stem-helix peptide (PDB 8DTX). (**L**) rhesus antibody DHIA-Stem30.06 with SARS-CoV-2 S2 stem-helix peptide (this study). Detailed interactions of conserved residues in antibodies and S2 stem-helix peptides are shown on the right. (**M**) Sequence alignment of the S2 stem region of betacoronaviruses/alphacoronaviruses. Conserved identical residues in betacoronaviruses are highlighted with deep blue boxes, and similar residues are shown in cyan boxes (amino acids that scored greater than or equal to 0 in the BLOSUM62 alignment score matrix are counted as similar here (*75*).

The sequence of the S2 stem-helix peptide is highly conserved across betacoronaviruses (Fig. 5M), and these conserved residues contribute extensively to the recognition by all eight rhesus antibodies (Fig. 5E-H, S7F-J), explaining their broad cross-reactivity (Table S2, S3). In each structure, hydrophobic paratope residues interact with the hydrophobic face of the SARS-CoV-2, MERS-CoV, or HCoV-OC43 S2 stem-helix peptides represented by residues F1148/1231/1233, L1152/1235/1237, Y1155/F1238/W1240, and F1156/1239/1241 (the numbers correspond to SARS-CoV-2, MERS-CoV, and HCoV-OC43 numbering, respectively) (Fig. 5E-H, S7F-J, S8). Antibody paratope residues also formed numerous H-bonds and salt bridges with the SARS-CoV-2, HCoV-OC43, and MERS-CoV S2 stem-helix peptides, further stabilizing the interaction (Fig. 5E-H, S7F-J).

We next compared the eight rhesus antibody structures with published structures of ten human S2 stem-helix bnAbs and two rhesus S2 stem-helix antibodies (*25, 27–31*) (Fig. S9I-R). Remarkably, rhesus antibodies RB6-Stem20.01 and DHIA-Stem30.06 approached the S2 stem-helix region at nearly identical angles to those of the human IGHV1-46/IGKV3-20 public antibodies S2P6, CC99.103, CC68.109, COV30-14, and COV89-22(*49, 52, 53*) (Fig. 5I-L, S10A-C). Despite this structural convergence, their germline origins are distinct: heavy-chain sequence identity to human IGHV1-46 was only 41.8% and 45.9% for RB6-Stem20.01 and DHIA-Stem30.06, respectively, and light-chain identity to human IGKV3-20 was only 47.4% and 49.5%, respectively (Fig. S10D-H). In contrast to the divergent frameworks, sequence and structure comparisons revealed conservation of key paratope residues across rhesus and human antibodies. For example, for rhesus RB6-Stem20.01, DHIA-Stem30.06, and human IGHV1-46/IGKV3-20 public antibodies, V_H_ Y33, I/L50, and V_L_ Y32, Y91, and V/F/Y96 interact with S2 stem-helix hydrophobic residues F1148, L1152, Y1155, and F1156 (SARS-CoV-2 numbering) or F1231, L1235, F1238, and F1239 (MERS-CoV numbering) (Fig. 5I-L, S10). These conserved paratope residues appear critical for recognition of the S2 stem-helix region and enable similar antibody binding angles despite relatively low overall sequence identity.

Additional structural comparisons revealed analogous convergence of binding mode within macaque antibodies. The binding angle and epitope of DHIA-Stem7.02 with the SARS-CoV-2 S2 stem-helix peptide were nearly identical to those of DHIA-Stem3.02 with the HCoV-OC43 S2 stem-helix peptide (Fig. S11A), consistent with highly similar paratope residues and interactions between the two antibodies (Fig. S11B-G). Likewise, RB6-Stem13.01 bound the SARS-CoV-2 and MERS-CoV S2 stem-helix peptides at nearly identical angles and dispositions, with four aromatic and aliphatic residues of the two peptides adopting the same side-chain rotamers and forming similar hydrophilic and hydrophobic interactions with RB6-Stem13.01 (Fig. S11H-K). The high structural similarity confirms that S2 stem-helix antibodies recognize a conserved site across betacoronaviruses.

Overall, these findings show that epitope-focused S2 stem-helix nanoparticle immunization elicits antibodies with convergent genetic features, conserved binding modes, and shared recognition of a structurally conserved S2 stem-helix epitope across betacoronaviruses, closely paralleling human public stem-helix bnAb responses (*25*).

## Discussion

The development of vaccines that protect against future zoonotic betacoronaviruses remains a major unmet challenge. Here, we show that an epitope-focused nanoparticle vaccine targeting the conserved S2 stem-helix bnAb site can overcome the immunological subdominance of this poorly accessible site and elicit broadly neutralizing antibody responses in non-human primates. Our study demonstrates that a rationally engineered epitope-focused nanoparticle immunogen can consistently induce serum neutralization directed to the S2 stem-helix together with monoclonal antibodies with specificity spanning sarbecoviruses and merbecoviruses. These vaccine-induced antibodies protect against SARS-CoV-2 and MERS-CoV *in vivo*—two pathogenic human betacoronaviruses—and exhibit immunogenetic and structural convergence with human S2 stem-helix bnAbs. Together, these findings provide proof-of-concept that antibody-guided epitope-focused design can yield immunogens that elicit protective bnAbs against conserved, structurally occluded vulnerable sites on complex viral glycoproteins in an outbred non-human primate model, representing a major advance toward the development of a pan-betacoronavirus vaccine.

Coronavirus infection and conventional spike-based vaccines predominantly focus antibody responses toward immunodominant but highly variable regions, such as the RBD (*13, 54–57*), resulting in responses with limited cross-strain reactivity. By multivalently displaying the S2 stem-helix on ferritin nanoparticles, our approach focused B cell responses toward this relatively immunoquiescent but highly conserved region of the spike protein. The robust stem-helix antibody responses elicited by S2 stem-helix nanoparticle vaccination in macaques, compared with the minimal responses observed after natural infection, conventional vaccination, or hybrid immunity in humans, highlights the power of epitope-focused vaccine design to reshape immunodominance and selectively amplify antibody responses against conserved occluded viral glycoprotein epitopes.

The structural convergence of vaccine-elicited antibodies further supports this epitope-targeted vaccine strategy. Two rhesus antibodies, RB6-Stem20.01 and DHIA-Stem30.06, recognize the S2 stem-helix using binding orientations and conserved paratope features nearly identical to those of the human public IGHV1-46/IGKV3-20 bnAb class despite limited overall sequence identity. Additional rhesus antibodies recognize the same conserved epitope through related binding modes, demonstrating that epitope-focused vaccination can reproducibly elicit convergent antibody solutions across species.

The isolation of a large number of bnAbs demonstrates that stem-helix nanoparticle immunization efficiently recruits and expands cross-reactive B cell lineages. However, several animals generated potent pan-betacoronavirus bnAb lineages despite limited or undetectable serum MERS-CoV neutralization, revealing a disconnect between the presence of desirable memory B cells and their contribution to circulating antibody responses. Thus, future vaccine development must focus not only on engaging bnAb precursors but also on promoting their preferential expansion, maturation, and differentiation into durable plasma cell responses.

Although our study establishes that serum neutralization against the S2 stem-helix can be achieved, improving the consistency and magnitude of pan-betacoronavirus serum neutralization remains a key objective. Several observations provide a roadmap for improving this vaccine strategy. First, the enhanced potency following BA.1 mRNA boosting suggests that sequential immunization strategies incorporating increasingly native-like antigen presentations, or heterologous boosts with divergent betacoronaviruses (e.g., from MERS-CoV or other betacoronaviruses) may further drive maturation toward broader activity. Second, additional improvements in epitope density, nanoparticle geometry, and antigen presentation may increase recruitment of favorable precursors while limiting competing off-target responses. Third, combining S2 stem-helix targeting with other conserved neutralizing epitopes, including the fusion peptide and conserved RBD sites (*20, 58*), may further enhance breadth and durability.

Complete protection against respiratory viruses will likely require coordinated systemic and mucosal immunity (*59–62*). Future studies should explore combinations of epitope-focused immunogens with platforms capable of inducing durable mucosal responses, including intranasal vaccination following systemic priming (prime-and-pull) or intranasal vector-based boosts after systemic priming to establish durable mucosal B- and T-cell memory (*61–66*). Incorporating mRNA delivery platforms may further improve immunogenicity and persistence, while heterologous mucosal boosts could complement broad systemic neutralization by reducing viral acquisition and transmission (*62, 63, 67, 68*). Finally, understanding how pre-existing immunity to seasonal human betacoronaviruses shapes the recruitment and maturation of S2 stem-helix-specific B cell responses will be critical for clinical translation, as prior coronavirus exposures may either facilitate or divert responses toward this conserved epitope.

In summary, our study demonstrates that rational epitope-focused nanoparticle vaccination is a powerful strategy for overcoming viral immunodominance and targeting conserved, relatively occluded bnAb epitopes on viral glycoproteins (*69*). The implications of this approach extend well beyond coronaviruses. Many high-value vaccine targets, including the influenza HA stem and HIV-1 Env fusion elements, are similarly constrained by epitope masking and limited accessibility, prompting extensive efforts to enhance their immunogenicity through structure-based vaccine design (*70–74*). Some of these targets also face the additional challenge of substantial sequence variability—for example, the HIV-1 fusion peptide (*70*). In contrast, the exceptional conservation of the S2 stem-helix epitope across betacoronaviruses appears sufficient to support the development of a broadly protective, pan-betacoronavirus vaccine.

## Acknowledgements

This work was supported by National Institutes of Health-(NIH), National Institute of Allergy and Infectious Diseases-(NIAID) awards, R01AI170928 (R.A.), R01 AI190286 (I.A.W., M.Y.), and the Gates Foundation INV-004923 (A.B.W., I.A.W., D.R.B.). We thank Henry Tien for technical support with the crystallization robot. We are grateful to the staff of the National Synchrotron Light Source II (NSLS-II) beamlines 17-ID-1 and 17-ID-2. NSLS-II is a U.S. Department of Energy Office of Science User Facility. This research used resources of the National Synchrotron Light Source II, a U.S. Department of Energy (DOE) Office of Science User Facility operated for the DOE Office of Science by Brookhaven National Laboratory under Contract No. DE-SC0012704. Beamline 5.0.3 of the Advanced Light Source, a U.S. DOE Office of Science User Facility under Contract No. DE-AC02-05CH11231, is supported in part by the ALS-ENABLE program funded by the National Institutes of Health, National Institute of General Medical Sciences, grant P30 GM124169-01. Use of the Stanford Synchrotron Radiation Lightsource, SLAC National Accelerator Laboratory, is supported by the U.S. Department of Energy, Office of Science, Office of Basic Energy Sciences under Contract No. DE-AC02-76SF00515. The SSRL Structural Molecular Biology Program is supported by the DOE Office of Biological and Environmental Research, and by the National Institutes of Health, National Institute of General Medical Sciences (P30GM133894). The contents of this publication are solely the responsibility of the authors and do not necessarily represent the official views of NIGMS or NIH.

## Author contributions

P.Z., Z.F., W.-t.H., I.A.W., D.R.B., and R.A. conceived and designed the study. P.Z., W.-t.H., T.C., and G.A. conducted animal immunization studies and processed plasma samples. W.R. and M.F. recruited donors and collected and processed plasma samples. P.Z., W.-t.H., X.L., Y. Zhang, L.V., T.C., S.C., N.M., G.A., K.D., B.L., R.R.C., and R.N. performed BLI assays, ELISAs, virus preparations, neutralization assays, and isolation and characterization of monoclonal antibodies. P.Z., X.L., L.V., T.C., and G.A. prepared the monoclonal antibody, Fab, spike, and stem-helix ferritin nanoparticle proteins. P.Z. and W.-t.H. performed cell surface binding assays. P.Z., Z.F., M.Y., and T.C. conducted DSC assays. P.Z., Y. Zhu, and R.A. performed the immunogenetic analysis of the antibodies. Z.F., M.Y., and I.A.W. determined the crystal structures and analyzed the structural data. W.-H.L. and A.B.W. conducted negative stain electron microscopy studies. J.D.A. and M.M.C. performed glycan analysis A.W. and M.M. prepared the SMNP adjuvant. M.- G.A. and D.W. prepared the BA.1 spike mRNA vaccine. E.T.M., N.A.K., R.S.B. and L.E.G. performed the murine challenge study. P.Z., Z.F., W.-t.H., M.Y., Y. Zhang, S.C., N.M., K.D., B.L., R.R.C., W.-H.L., J.D.A., M.M.C., A.B.W., D.J.I., M.-G.A., D.W., R.S.B., L.E.G., I.A.W., D.R.B., and R.A. designed the experiments and/or analyzed the data. P.Z., Z.F., W.-t.H., I.A.W., D.R.B., and R.A. wrote the paper, and all authors reviewed and edited the paper.

## Competing interests

P.Z., Z.F., W.-t.H., M.Y., S.C., D.W., I.A.W., D.R.B., and R.A. are listed as inventors on a pending patent application describing the S2 stem-helix ferritin nanoparticle vaccines reported in this study.

## Data and materials availability

The data supporting the findings of this study are available within the paper and its supplementary information files or from the corresponding author upon reasonable request. Antibody sequences have been deposited in GenBank under accession numbers XXX-XXX. The X-ray coordinates and structure factors have been deposited to the RCSB Protein Data Bank under accession codes: 37GT, 37GS, 37GV, 37GW, 37GX, 37GY, 37IA, 37HA, and 37HB.

## Supplementary Materials

### MATERIAL AND METHODS

#### Cell lines

FreeStyle™ 293-F Cells (Thermo Fisher Scientific Cat# R79007) and Expi293F cells (Gibco Cat# A14527) were cultured in FreeStyl 293 Expression Medium (Gibco Cat# 12338018) and Expi293 Expression Medium (Gibco Cat# A1435101), respectively, in a shaker at 120 rpm with 8% CO_2_ at 37°C. Adherent HEK293T cells, HeLa-ACE2 and HeLa-DPP4 cells were grown in Dulbecco’s Modified Eagle Medium (DMEM), which contained 10% heat-inactivated FBS, 1% penicillin-streptomycin and 4mM L-Glutamine, and maintained in an incubator with 5% CO_2_ at 37°C. The stable hACE2 and hDPP4-expressing HeLa cell lines were generated using the protocol previously described (*54*). Briefly, HeLa cells were transfected by pBOB-hACE2 or hDPP4 plasmid with lentiviral packaging plasmids including pMDL, pREV, and pVSV-G (Addgene Cat# 12251, Cat# 12253, Cat# 8454) by Lipofectamine 2000 reagent (Thermo Fisher Scientific Cat# 11668019) to generate related lentivirus, then HeLa cells were infected by the lentiviruses.

#### Plasmids, proteins, and peptides

To generate soluble spike ectodomain proteins of human betacoronaviruses including SARS-CoV-1 (residues 1-1190; GenBank: AAP13567), SARS-CoV-2 (residues 1-1208; GenBank: MN908947), MERS-CoV (residues 1-1291; GenBank: APB87319.1), HCoV-HKU1 (residue 1-1295; GenBank: YP_173238.1) and HCoV-OC43 (residues 1-1300; GenBank: AAX84792.1), DNA fragments synthesized by GeneArt were cloned into phCMV3 vector (Genlantis Cat# P003300) by HiFi DNA assembly (New England Biolabs Cat# E2621L) according to the manufacturer’s instructions. To produce stable spike trimer spike proteins in a pre-fusion state (*76, 77*), double proline substitutions (2P) were introduced into the spike S2 subunit,

K968P/V969P for SARS-CoV-1, K986P/V987P for SARS-CoV-2, V1060P/L1061P for MERS-CoV, A1071P/L1072P for HCoV-HKU1 and A1078P/L1079P for HCoV-OC43. A “GSAS” linker was used to replace the furin cleavage site in SARS-CoV-1 (residues 664–667), SARS-CoV-2 (residues 682–685), MERS-CoV (residues 748–751), HCoV-HKU1 (residues 756-760) and HCoV-OC43 (residues 762–766); T4 fibritin (foldon) was incorporated at the C-terminus of the spike proteins to help form stable trimers. An HRV-3C protease cleavage site, 6x His-tag for protein purification, and AviTag for biotinylation, which were spaced by GS-linkers, were added at the C-terminus of spike after T4 fibritin.

To generate the stem-helix ferritin nanoparticles shwon in Figure S1, DNA fragments encoding different nanoparticle constructs were synthesized by GeneArt and cloned into the phCMV3 expression vector (Genlantis, Cat# P003300). In all constructs, the stem-helix peptide sequences were inserted downstream of the human tissue plasminogen activator (tPA) signal peptide (MDAMKRGLCCVLLLCGAVFVSPSQEIHARFRRGAR) and connected through a G4S linker to a bullfrog ferritin N-terminal extension , which was fused to the N-terminus of the *Helicobacter pylori* ferritin scaffold containing three previously reported mutations (N19Q, C31S, and S111C) (*36*). The Stem-FR nanoparticle contained a single stem-helix peptide, whereas the Stem-3R-FR and Mosaic-Stem-3-FR nanoparticles contained three tandem copies of the stem-helix peptide, with the three peptide copies separated by GS linkers.

To generate pseudoviruses of sarbecoviruses and MERS-CoV, codon-optimized DNA fragments of full-length spikes without the ER retrieval signals were synthesized at GeneArt. In addition to SARS-CoV-1, SARS-CoV-2 and MERS-CoV, spike DNA fragments of Pang17 (residues 1-1249, GenBank: QIA48632.1), WIV1 (residues 1-1238, GenBank: KF367457) and SHC014 (residue 1-1238, GenBank: AGZ48806.1) were constructed into the phCMV3 vector.

To obtain stem-helix peptides for ELISA and BLI assay, N-terminal biotinylated peptides corresponding to the stem-helix of SARS-CoV-2 (PLQPELDSFKEELDKYFKNHTSPDV), MERS-CoV (PLLGNSTGIDFQDELDEFFKNVSTSIP), HCoV-HKU1 (HSVPKLSDFESELSHWFKNQTSIAP) and HCoV-OC43 (TSIPNLPDFKEELDQWFKNQTSVAP) were synthesized at GenScript. The stem-helix peptides for crystallization were without biotinylation.

#### Human samples

Plasma from convalescent COVID-19 donors, two-dose SARS-CoV-2 spike mRNA vaccinated donors, and COVID-19 recovered-vaccinated donors were provided through the “Collection of Biospecimens from Persons Under Investigation for 2019-Novel Coronavirus Infection to Understand Viral Shedding and Immune Response Study” (UCSD IRB#200236), as previously reported (*42*). The study protocol was approved by the UCSD Human Research Protection Program. Convalescent donor samples were collected from individuals with a confirmed COVID-19 diagnosis regardless of gender, race, ethnicity, disease severity, or other underlying medical conditions. All human donors were assessed for medical decision-making capacity using a standardized, approved assessment and provided written informed consent prior to enrollment in the study.

#### Animal models

All 12 Indian rhesus macaques used in this study were housed at Alpha Genesis, Inc. in compliance with the guidelines set by the Association for Assessment and Accreditation of Laboratory Animal Care (AAALAC). All experimental procedures were approved by the Institutional Animal Care and Use Committees (IACUC) of Alpha Genesis, Inc. (protocol 22-16). Macaques were sedated when taking blood samples and received care in accordance with AAALAC guidelines and best practice standards.

#### Expression and purification of soluble recombinant human betacoronavirus spike proteins

For expression of soluble recombinant human betacoronavirus spike proteins, after mixing the filtered 350 μg plasmids in 15mL Opti-MEM™ with 1.8 mL 40K PEI (1mg/mL, Kyfora, Cat# 24765-1) in 15mL Opti-MEM (Thermo Fisher Scientific Cat# 31985070), the mixture was incubated for 30 minutes at room temperature and then transferred into 1L FreeStyle293-F cells at a density of 1X10^6^ cells/mL. Four days after transfection, the supernatant was harvested at 4000rpm for 30 min and filtered through a 0.22μm membrane. HisPur Ni-NTA Resin (Thermo Fisher Scientific Cat# 88221) could specifically bind to the His tag to purify spike proteins. After washing the resin with 25 mM imidazole in PBS (pH 7.4) for at least 3 bed volumes, the protein was eluted by 25 mL 250 mM imidazole in PBS buffer (pH 7.4) at slow gravity speed (∼4 sec/drop), then concentrated by 100KDa Amicon tubes (Millipore Cat# UFC9100). The protein was pooled and concentrated again for future use after being further purified by size-exclusion chromatography (SEC) using Superdex 200 Increase 10/300 GL column (Cytiva Cat# 28990944).

#### Expression and purification of stem-helix ferritin nanoparticle

For the expression of stem-helix ferritin nanoparticle, 1L FreeStyle293-F cells at a density of 2X10^6^ cells/mL were transfected by the mixture of filtered 1mg plasmids in 15mL Opti-MEM™ and 5 mL 40K PEI (1mg/ml, Kyfora, Cat# 24765-1) in 15mL Opti-MEM (Thermo Fisher Scientific Cat# 31985070). Four days later, the supernatant was harvested at 4000rpm for 30 min and filtered through a 0.22μm membrane. Stem-helixferritin nanoparticles were purified from the supernatant by the *Galanthus nivalis* lectin (GNL) agarose (Vector Labs Cat# AL-1243-5). The bound proteins were washed by PBS for at least 3 bed volumes, and then eluted by 30mL 1 M methyl-α- D-mannopyranoside in PBS. After concentration by 50KDa Amicon tubes (Millipore Cat# UFC9050), the proteins were further purified by SEC using Superose 6 Increase 10/300 GL column (Cytiva Cat# 29091596), and then pooled and concentrated again for future use.

#### ELISA assay for stem-helix peptides or recombinant spike proteins

For peptide enzyme-linked immunosorbent assay (ELISA), each well of 96-well half-area high binding plates (Corning Cat# 3690) was coated by 50 μL streptavidin (Jackson Immuno Research Labs Cat# 016-000-084) at 2 μg/mL in PBS overnight at 4°C. For recombinant spike protein ELISA, mouse anti-His antibody (Thermo Fisher Scientific Cat# MA1-21315-1MG) was coated onto the plates at the same concentration and amount of streptavidin (50ul of 2μg/ml). The plates were washed by 0.05% PBST for 3 times, and then blocked by 3% BSA for 2h at 37°C. After removing 3% BSA, 50 μL biotinylated peptide or His-tagged recombinant spike proteins at 2 μg/mL in 1% BSA was applied to plates and incubated at room temperature (RT). One hour later, the plates were washed three times by 0.05% PBST before adding serially diluted plasma samples or antibodies. The plates with plasma samples or antibodies were then incubated for 1h at RT. After another wash 3 times, 50 μL alkaline phosphatase (AP)-conjugated goat anti-human IgG Fc secondary antibody (Jackson ImmunoResearch Cat# 109-055-008) in 1:1000 dilution was added onto the plates and incubated for 1h at RT. After the final wash for 3 times, 50 μL phosphatase substrate (Sigma-Aldrich Cat# S0942-200TAB) dissolved in staining buffer (0.1 M glycine, 1 mM MgCl_2_ and1 mM ZnCl_2_, pH 10.4) was added into each well. Absorption at 405 nm was measured. Asymmetrical dose-response model of the Richard version in GraphPad Prism 8 (GraphPad Software) was used to calculate half-maximal effective concentration (EC_50_) or dilution (ED_50_) for mAb or plasma, respectively.

#### Cell surface binding assay

Flow cytometry-based cell surface binding of mAbs with human betacoronavirus spikes was performed as described previously (*22*). A total of 4x10^6^ HEK293T cells were seeded into 10cm round cell culture dishes and incubated at 37°C for 24h. The plasmids encoding full-length spikes were then transfected into HEK293T cells. After incubation for 36-48h at 37°C, the cells were harvested and distributed into 96-well round-bottom tissue culture plates for individual cell staining. For each reaction, cells were washed by 200 μL FACS buffer (1xPBS, 2%FBS, 2mM EDTA) for 3 times. Primary antibody in 50 μL staining buffer at a concentration of 10μg/mL was used to stain cells for 1h on ice. After washing three times by 200 μL FACS buffer, R-phycoerythrin (PE)-conjugated mouse anti-human IgG Fc antibody (SouthernBiotech Cat# 9040-09) and Zombie-NIR viability dye (BioLegend Cat# 423105), which were diluted at a ratio of 1:200 and 1:1000 in 50 μL FACS buffer, respectively, were applied to the plates and incubated on ice for 45 min in the dark. Following three washes with 200 μL FACS buffer, the cells were resuspended and analyzed by flow cytometry (BD Lyrics cytometer). FlowJo 10 software was used to calculate the binding data Mean Fluorescence Intensity (MFI). Mock-transfected 293T cells were used as a negative control.

#### BioLayer interferometry binding assay

An Octet RH96 system (Sartorius) was used to determine the binding of monoclonal antibodies with stem-helix ferritin nanoparticles or synthetic stem-helix peptides. The baseline was obtained by flowing Octet buffer (PBS with 0.1% Tween) for 60s, then anti-human IgG Fc capture (AHC) biosensors (Sartorius Cat# 18-5063) were used to capture IgG for 60s. AHC sensors were then transferred into the wells containing Octet buffer for another 60s, followed by into wells containing diluted stem-helix ferritin nanoparticle for 120s and into Octet buffer for disassociation for 240s. For monoclonal antibody binding to peptides, the method used was similar to the one above, except that streptavidin biosensors (Sartorius Cat# 18-5020) were used to capture N-terminal biotinylated peptide and the sensors were transferred into wells containing diluted monoclonal antibodies for association for 120s. To identify critical residues for antibody binding, alanine scanning was performed on a series of N-terminal biotinylated 25-mer SARS-CoV-2 stem-helix peptides with one single alanine mutation at each amino acid position. Unlike the intrinsic monovalent binding affinity measured for Fab fragments, binding of IgG antibodies to nanoparticles or peptides may involve a mixture of 2:1 and 1:1 binding modes because of the multivalent nature of IgG. Accordingly, the dissociation constants reported here are designated as apparent affinity (K_D_^App^) to reflect the overall binding affinity of IgG antibodies to the tested nanoparticles or peptides.

#### Competition BLI

In-tandem epitope binning experiment by Octet RH96 system was conducted to determine the binding epitopes of the isolated rhesus S2 stem-helix mAbs compared with human S2 stem-helix mAbs of known epitopes. Briefly, after obtaining baseline by flowing Octet buffer for 30s, streptavidin biosensors (Sartorius Cat# 18-5020) were used to capture N-terminal biotinylated SARS-CoV-2 spike protein at a concentration of 100nM in Octet buffer for 5 min. Unbound spike protein was removed by transferring the sensors into Octet buffer for 30s.Then, the protein bound sensors were moved into saturating antibodies (human s2 stem-helix mAbs) in Octet buffer at a concentration of 100 μg/mL for 10 min followed by transfer into 100 μg/mL competitor antibodies (rhesus S2 stem-helix mAbs) in Octet buffer for 5 min to measure binding in the presence of saturating antibodies. As a control, spike protein bound biosensors were directly transferred into competitor antibody solution. The percent inhibition (%) in binding was calculated with the formula: [Percent binding inhibition (%) = 1- (competitor antibody binding response in presence of saturating antibody / binding response of the competitor antibody without saturating antibody).

#### Pseudovirus production

HIV-based lentivirus backbone plasmid pCMV-dR8.2 dvpr (Addgene Cat# 8455), pBOB-Luciferase (Addgene Cat# 170674) and plasmids encoding various different truncated spike proteins, including SARS-CoV-1, SARS-CoV-2, WIV1, Pang17, MERS-CoV and SARS-CoV-2 variants of concern (VOCs, Omicron BA.4/5 and XBB.1.5), were co-transfected into HEK293T cells by lipofectamine 2000 (Thermo Fisher Scientific Cat# 11668019) to produce pseudoviruses (*78*). The medium was removed before adding fresh medium 12-16 hours post transfection.Supernatants containing pseudoviruses were collected 48 hours post transfection, and aliquoted and stored at -80°C until further use. The viral titers were measured by a bright-glo luciferase assay system (Promega Cat# E2620).

#### Neutralization assay

Pseudovirus neutralization assay was performed as previously described (*54*). Briefly, after mixing 25 μL of pseudovirus with 25 μL serial dilutions of purified antibodies or plasma in the wells of 96-well half-area TC-treated plate (Corning Cat# 3688), the plate was incubated for 1h at 37°C, then 50 μL medium containing 1X10^4^ HeLa-hACE2 or hDPP4 cells and 20μg/mL Dextran directly added to each well. After incubation at 37°C for 42-48 h, luciferase activity was measured. Compared to the virus controls, the reduction in luciferase activity was used to measure neutralizing activity. Fifty percent inhibitory concentration or dilution (IC_50_ or ID_50_) were calculated using the dose-response-inhibition model with 5-parameter Hill slope equation in GraphPad Prism 8 (GraphPad Software).

#### Immunization in rhesus macaques and blood processing

Two groups of rhesus macaques (6 animals in each group), evenly distributed by gender between ages 5-6 years, were immunized with stem-helix ferritin nanoparticles (along with the SMNP adjuvant) in combination with SARS-CoV-2 BA.1 spike mRNA vaccine. Each rhesus macaque received 2 nanoparticle immunization at week 0 and week 4. For each immunization, 100 μg nanoparticle and 375 μg SMNP adjuvant were administered subcutaneously and distributed into two injection sites (left and right mid-thigh) for every rhesus macaque. At week 12, both groups were boosted with 100 μg SARS-CoV-2 BA.1 spike mRNA LNP at two injection sites (left and right mid-thigh). Serial bleeds were collected over the course of the immunization experiment to evaluate antibody and B cell responses with the schedule: wk -1, wk 2, wk 6, wk 12 and wk 14.

Peripheral blood was collected in sterile vacutainers (DB Vacutainer Cat #364606) containing acid citrate dextrose formula A (ACD-A) as an anticoagulant. Following collection, samples were centrifuged at 1000g for 10 minutes at 20°C in sterile 50 mL conical tubes. Plasma was carefully collected while avoiding disruption of the buffy coat and red blood cell pellet, followed by a second centrifugation at 1500g for 15 minutes at 20°C to remove residual cellular material. The cell-depleted plasma was aliquoted into 2 mL cryovials (Sarstedt Cat # 72.694.396) and stored at −80°C. The cell fraction was resuspended in an equal volume of Hanks’ Balanced Salt Solution (HBSS) without calcium or magnesium (HBSS−/−) (Gibco Cat # 14175-079) containing 2 mM EDTA (Invitrogen Cat #15575-020) and distributed into 50 mL conical tubes. Additional HBSS−/− with EDTA was added to each tube to adjust the total volume to 35 mL. The cell suspension was layered over 14 mL of 96% Ficoll-Paque Plus (Cytiva Cat # 17144003) and centrifuged at 725g for 20 minutes at 20°C with slow acceleration and braking. Mononuclear cells at the Ficoll interface were collected, transferred to fresh 50 mL conical tubes containing HBSS−/− with EDTA, and washed by centrifugation at 200g for 15 minutes at 20°C. Following removal of the supernatant, the cell pellet was resuspended in 40 mL of HBSS containing calcium and magnesium (HBSS+/+) (Gibco Cat # 24020-117) supplemented with 1% fetal bovine serum (FBS) (Cytiva Cat # SH300.71.03). The suspension was centrifuged at 200g for 15 minutes at 20°C, and the resulting supernatant was removed. This centrifugation step preferentially pelleted white blood cells (WBCs) while retaining most platelets in suspension. The mononuclear cell pellet was gently resuspended in the residual media, followed by the addition of HBSS+/+ with 1% FBS to a final volume of 10 mL. Cell number and viability were determined using ViaStain AOPI solution (Revvity Cat #CS2-0106-25ml) and a Cellometer Auto 2000 instrument (Revvity, Waltham, MA). The cells were then centrifuged at 300g for 10 minutes at 20°C, after which the supernatant was discarded, and the pellet was resuspended in CryoStor CS5 cryopreservation medium (Stemcell Technologies Cat # 07930) at a final concentration of 5–10 × 10⁶ cells/mL. The suspension was aliquoted into 1.8 mL cryovials (Thermo Scientific Cat # 374503), cryopreserved overnight at −80°C using a Corning CoolCell LX (Corning cat #432002) or FTS30 (Corning cat #432006) freezing container, and subsequently transferred to vapor-phase liquid nitrogen for long-term storage.

#### Antigen-specific B cell sorting and monoclonal antibody isolation

The strategy of antigen-specific memory B cell sorting was performed as described in previously reported papers (*39, 79, 80*). Pre-warmed RPMI1640 medium (Thermo Fisher Scientific Cat# 11875085) with 50% FBS was used to thaw the frozen PBMC samples. After centrifuging for 5 min at 400g and discarding the supernatant, the cells were resuspended in 5 mL FACS buffer (PBS with 2% FBS and 2 mM EDTA). Antibodies specific for cell surface markers and fluorescently labeled were prepared at 1:100 dilution as a mixture in FACS buffer. T-cell markers CD3 (APC-Cy7, BD Biosciences Cat# 557757), CD4 (APC-Cy7, BioLegend Cat# 317418), CD8 (APC-Cy7, BD Biosciences Cat# 557760), monocyte marker CD14 (APC-H7, BD Biosciences Cat# 561384) and IgM (PE, BioLegend Cat# 314508) were stained for negative selection. B-cell markers CD19 (PerCP-Cy5.5, BioLegend Cat# 302230), CD20 (PerCP-Cy5.5, BioLegend Cat# 302326) and IgG (BV786, BD Biosciences Cat# 564230) were stained for IgG^+^ B cell isolation. SARS-CoV-2 spike protein with Avi-tag was conjugated to both streptavidin-BV421 (BD Biosciences Cat# 563259) and -AF488 (Thermo Fisher Scientific Cat# S11223), while the MERS-CoV spike protein with Avi-tag was conjugated to streptavidin-AF647 (Thermo Fisher Scientific Cat# S21374). After incubating PBMC with the mixture of fluorescent antibodies for 15 min in the dark, spike protein probes were added to PBMC and incubated for 30 min. Then, FVS510 Live/Dead stain (Thermo Fisher Scientific Cat# L34966) diluted in 1:300 in FACS buffer was added to the samples and incubated for 15 min. After washing 3 times with FACS buffer, the PBMCs were resuspended by FACS buffer to a concentration of 20-40 X 10^6^/mL. After being filtered through cell strainer snap cap into FACS tubes (Fisher Scientific Cat# 08-771-23), spike protein-specific CD3^-^CD4^-^CD8^-^CD14^-^IgM^-^IgG^+^ CD19^+^CD20^+^ live memory B cells were sorted by BD FACSMelody sorter. For PBMCs of rhesus macaques DHIA, DHHP and K872 from week 6, SARS-CoV-2 spike double-positive B cells were sorted as single cells into 96-well plates on a cooling platform. For week 6 PBMCs of rhesus macaque RB6 and week 14 PBMCs of rhesus macaques DHIA and RB6, SARS-CoV-2 and MERS-CoV spike triple-positive B cells were sorted.

Reverse transcription PCR reaction was conducted to generate cDNA from the sorted cells by Superscript IV Reverse Transcriptase (Invitrogen Cat# 18090010), 10mM dNTPs (Invitrogen Cat# 18427088), random hexamers (Gene Link Cat# 26-4000-03), Ig gene-specific primers, 0.1M DTT, RNAseOUT (Invitrogen Cat# 10777019), and 10% Igepal (Sigma-Aldrich Cat# 18896). The cDNA was then used as template to perform two rounds of nested PCR reactions to amplify IgG heavy and light chain variable regions by Hot Start DNA Polymerases (QIAGEN Cat# 203643) and specific primer sets as described previously (*80*). After purification with SPRI beads (Beckman Coulter Cat# B23318), the heavy and light chain variable region PCR products were cloned into the expression vectors containing constant domains of human IgG1 heavy or kappa/lambda chains, respectively, by HiFi DNA assembly (New England Biolabs Cat# E2621L). After transforming into competent *E. coli* cells, single colonies were picked for sequencing, analysis on IMGT V-Quest online tool (http://www.imgt.org), and downstream plasmid production.

#### Expression and purification of monoclonal antibodies

Expi293F cells were co-transfected by paired heavy and light chain plasmids to produce monoclonal antibodies. In brief, 12μg heavy chain plasmid and 12 μg light chain plasmid were mixed into 3 mL Opti-MEM (Thermo Fisher Scientific Cat# 31985070), and then 24 μL FectoPRO reagent (Polyplus Cat# 116-001) was added and fully mixed. After incubation at RT for 15 min, the mixture was added to 30 mL Expi293F cells at a concentration of 2.8 X 10^6^ cells/mL, and then cultured in a shaker. After 24 hours post transfection, 300 μL 0.3 M sodium valproic acid solution and 275 μL 45% glucose solution were used to feed the transfected cells. Supernatants were harvested by centrifugation at 2500g for 30 min and filtering through 0.22μm membrane after 4 days post transfection. Protein A (Cytiva Cat# 17096302) and protein G Sepharose (Cytiva Cat# 17061805) mixed at a 1:1 ratio were added into the supernatant followed by rotating overnight at 4°C. After loading the supernatant into Econo-Pac column (BioRad Cat# 7321010) and washing the column by 1 column volume of PBS, antibodies were eluted by 10 mL of 0.2 M citric acid (pH 2.67) into tubes containing 1 mL 2M Tris Base solution. The antibodies were concentrated and exchanged buffer into PBS by 30K Amicon centrifugal tubes (Millipore Cat# UFC903024). Finally, the antibodies were concentrated into smaller volumes for further use.

#### Expression and purification of Fabs

To generate Fabs, a stop codon was introduced into the IgG1 heavy chain plasmid after amino acids “KSC” of the CH1 domain. Just as the abovementioned method to express and purify monoclonal antibodies, the heavy chain with stop codon and light chain were co-transfected into Expi293F cells. After harvesting the supernatant, the Fabs were purified by CaptureSelect CH1-XL Affinity Matrix (Thermo Fisher Scientific Cat#1943462250). The Fabs were eluted by 0.2 M citric acid (pH 2.67) into tubes containing 2M Tris Base solution. After concentration, the Fabs were further purified by SEC using a Superdex 200 Increase 10/300 GL column (Cytiva Cat# 28990944). Selected fractions were pooled and concentrated again for further use.

#### Crystallization and structural determination

RB6-Stem13.01 (13 mg/mL) with 10× (molar ratio, same as below) SARS-CoV-2 S2 stem-helix peptide (1140-PLQPELDSFKEELDKYFKNHTSPDV-1164), DHIA-Stem7.02 (13 mg/mL) with 10× SARS-CoV-2 S2 stem-helix peptide, RB6-Stem20.01 (11 mg/mL) with 10× SARS-CoV-2 S2 stem-helix peptide, RB6-Stem7.01 (13 mg/mL) with 10× SARS-CoV-2 S2 stem-helix peptide, DHHP-Stem20.01 (13 mg/mL) with 10× SARS-CoV-2 S2 stem-helix peptide, DHIA-Stem30.06 (12 mg/mL) with 10× SARS-CoV-2 S2 stem-helix peptide, DHIA-Stem3.02 (13 mg/mL) with 10× HCoV-OC43 S2 stem-helix peptide (1224-TSIPNLPDFKEELDQWFKNQTSVAP-1249), DHIA-Stem15.02 (13 mg/mL) with 10× MERS-CoV S2 stem-helix peptide (1220-PLLGNSTGIDFQDELDEFFKNVSTSIP-1247), and RB6-Stem13.01 (13 mg/mL) with 10× MERS-CoV S2 stem-helix peptide, were screened for crystallization using the 384 conditions of the JCSG Core Suite (Qiagen) on our robotic CrystalMation system (Rigaku) at Scripps Research. Crystallization trials were set up by the vapor diffusion method in sitting drops containing 0.1 μL of protein and 0.1 μL of reservoir solution. Diffraction-quality crystals were obtained in the following conditions:

1. RB6-Stem13.01/SARS-CoV-2 S2 stem-helix peptide: 0.2 M potassium nitrate, 10% (v/v) ethylene glycol, and 20% (w/v) polyethylene glycol 3350 at 20°C. Crystals appeared on day 3 and were harvested on day 15.
2. DHIA-Stem7.02/SARS-CoV-2 S2 stem-helix peptide: 0.1 M sodium acetate pH 4.6, 0.2 M ammonium sulfate, 10 %(v/v) ethylene glycol, and 25% (w/v) polyethylene glycol 4000 at 20°C. Crystals appeared on day 3 and were harvested on day 10.
3. RB6-Stem20.01/SARS-CoV-2 S2 stem-helix peptide: 2 M ammonium sulfate and 5% (v/v) 2-propanol at 20°C. Crystals appeared on day 3 and were harvested on day 13 by soaking in reservoir solution supplemented with 15% (v/v) ethylene glycol as cryoprotectant.
4. RB6-Stem7.01/SARS-CoV-2 S2 stem-helix peptide: 24% (w/v) polyethylene glycol 3350 and 0.237 M potassium acetate at 20°C. Crystals appeared on day 3 and were harvested on day 6.
5. DHIA-Stem3.02/HCoV-OC43 S2 stem-helix peptide: 0.1 M HEPES pH 7.5 and 70% (v/v) 2-methyl-2,4-pentanediol at 20°C. Crystals appeared on day 3 and were harvested on day 5 by soaking in reservoir solution supplemented with 15% (v/v) ethylene glycol as cryoprotectant.
6. DHHP-Stem20.01/SARS-CoV-2 S2 stem-helix peptide: 0.1 M CHES pH 9.5 and 40% (v/v) polyethylene glycol 600 at 20°C. Crystals appeared on day 7 and were harvested on day 14.
7. DHIA-Stem30.06/SARS-CoV-2 S2 stem-helix peptide: 0.2 M ammonium formate, 10% (v/v) ethylene glycol, 20% (w/v) polyethylene glycol 3350. at 20°C. Crystals appeared on day 3 and were harvested on day 14.
8. DHIA-Stem15.02/MERS-CoV S2 stem-helix peptide: 0.1 M sodium citrate pH 5.6, 0.2 M ammonium acetate, and 40% (w/v) polyethylene glycol 4000 at 20°C. Crystals appeared on day 7 and were harvested on day 14.
9. RB6-Stem13.01/MERS-CoV S2 stem-helix peptide: 0.2 M potassium formate and 20% (w/v) polyethylene glycol 3350 at 20°C. Crystals appeared on day 3 and were harvested on day 7 by soaking in reservoir solution supplemented with 15% (v/v) ethylene glycol as cryoprotectant.

The crystals were then flash-cooled and stored in liquid nitrogen until data collection. Diffraction data were collected at cryogenic temperature (100 K) at National Synchrotron Light Source II (NSLS-II) beamlines 17-ID-1, 17-ID-2, Stanford Synchrotron Radiation Lightsource (SSRL) beamlines 12-1 (12–2), and Advanced Light Source (ALS) beamline 5.0.3 with beam wavelengths of 0.92010 (0.92009) Å, 0.97934 Å, 0.97946 Å, and 0.97648 Å respectively. Diffraction data were processed with HKL2000 (*81*). Structures were solved by molecular replacement using PHASER (*82*) using SARS-CoV-2 stem-helix peptide in complex with antibody CHM-16 (*28*). Iterative model building and refinement were carried out in Coot (*83*) and PHENIX (*84*), respectively. Epitope and paratope residues, as well as their interactions, were identified by accessing PISA at the European Bioinformatics Institute (http://www.ebi.ac.uk/pdbe/prot_int/pistart.html) (*85*).

#### Melting temperature measure by differential scanning calorimetry (DSC)

Thermal stability and melting temperature of stem-helix ferritin nanoparticles were measured on a MicroCal VP-Capillary calorimeter (Malvern Panalytical) at a concentration 1mg/mL for each sample in PBS buffer at a scanning rate of 90 °C/hour from 20°C to 120°C. Data were analyzed by the VP-Capillary DSC automated data analysis software.

#### Site-specific glycan analysis by LC-MS

50 µg aliquots of each S2 stem-helix nanoparticle sample were denatured for 1 h in 50 mM Tris/HCl, pH 8.0 containing 6 M urea and 5 mM dithiothreitol (DTT), alkylated with 20 mM iodoacetamide for 1 h in the dark, and incubated for a further 1 h with 20 mM DTT to quench residual iodoacetamide. Samples were buffer exchanged into 50 mM Tris/HCl, pH 8.0 using Vivaspin columns (3 kDa) and aliquots digested separately overnight with trypsin (Mass Spectrometry Grade, Promega), chymotrypsin (Promega), or alpha-lytic protease (Sigma Aldrich) at a ratio of 1:30 (w/w). Peptides were dried, extracted using C18 ZipTips (Merck Millipore), dried again, and re-suspended in 0.1% formic acid.

Peptides were analyzed by nanoLC-ESI MS on an Ultimate 3000 HPLC (Thermo Fisher Scientific) coupled to an Orbitrap Eclipse Tribrid mass spectrometer (Thermo Fisher Scientific), using stepped higher energy collision-induced dissociation (HCD) fragmentation. Separation was performed on an EasySpray PepMap RSLC C18 column (75 µm × 75 cm) with an in-line PepMap Neo Trap Cartridge, at 300 nL/min using a 280-min gradient (4–32% acetonitrile in 0.1% formic acid over 260 min, followed by 20 min alternating between 76% and 4% acetonitrile in 0.1% formic acid to ensure complete elution). The spray voltage was 2.5 kV, the heated capillary 55 °C, and the ion transfer tube 275 °C. Spectra were acquired over 375–1500 m/z with precursor and fragment detection in the Orbitrap (MS1 = 120,000; MS2 = 30,000), standard AGC target and automatic injection times. Stepped HCD collision energies of 15, 25 and 45% were applied and the MS2 for each energy combined.

Glycopeptide fragmentation data were extracted using Byos (Version 5.5; Protein Metrics Inc.) and searched against the Protein Metrics 305 N-glycan library with a precursor mass tolerance of 4 ppm, fragment tolerance of 10 ppm, and a 1% false discovery rate. Assignments were evaluated manually and scored as true-positive when the expected b and y fragment ions were observed alongside oxonium ions corresponding to the assigned glycan. Relative glycan abundance and the unoccupied proportion at each site yielding data were determined by comparing extracted ion chromatographic areas for glycopeptides sharing an identical peptide sequence, summing all charge states.

#### Negative stain EM data collection and processing for stem-helix ferritin nanoparticles

S2 stem-helix ferritin nanoparticle protein was diluted to 0.02 mg/mL in 1x Tris-buffered saline, 3 uL applied to a 400mesh Cu grid, blotted with filter paper, and stained with 2% uranyl formate. Micrographs were collected on a 120kV Thermo Fisher Tecnai Spirit microscope with a FEI Eagle CCD (4k) camera (52,000 magnification, 2.06 Å pixel size) or a 200kV Thermo Fisher Scientific Talos with a Thermo Fisher Scientific CETA 4K CMOS camera (73,000X magnification, 1.981 Å pixel size) using Leginon automated image collection software (*86*). Particles were picked using DogPicker (*87*) and 2D classification was done using iterative multi-variate statistical analysis (MSA)/multi-reference alignment (MRA) (*88*).

#### In vivo virus challenge in mouse model

SARS-CoV-2 challenge: 20-week-old female BALB/c mice (strain 047) from Envigo (now Inotiv) were intraperitoneally injected with 300mg of antibody and 12 hours later intranasally infected with 1000 PFU of SARS-CoV-2 MA10 (*89*) diluted in sterile PBS. Mice were weighed daily and respiratory function was assessed at days 2 and 5 post infection via whole body plethysmography (*90*). At day 5 post-infection, mice were euthanized via isoflurane overdose and lungs assessed for gross pathology before tissue collection. Plaque assays were performed on VeroE6 cells using homogenized and serially diluted lung tissue, the limit of detection was 100 PFU per lung.

MERS-CoV challenge: 20-week-old male and female hDPP4 288/330 mice (*91*) were intraperitoneally injected with 300mg of antibody and 12 hours later intranasally infected with 10^5^ PFU of MERS maM35c4 (*92*). Mice were weighed daily and respiratory function was measured at 3 days post infection. At day 5 post-infection, mice were euthanized via isoflurane overdose and lungs assessed for gross pathology before tissue collection. Plaque assays were performed on Vero CCL81 cells (ATCC) using homogenized and serially diluted lung tissue, the limit of detection was 10 PFU per lung.

#### Immunogenetic analysis of rhesus S2 stem-helix mAbs

mmunogenetic analyses were performed on isolated rhesus S2 stem-helix mAbs from four vaccinated rhesus macaques. Immunoglobulin heavy- and light-chain variable region sequences were analyzed using IgBLAST with the IMGT rhesus macaque immunoglobulin germline reference database to assign V, D, and J genes, identify CDR3 sequences, and determine somatic hypermutation (SHM) frequencies. Clonal lineage assignment was performed using paired heavy- and light-chain sequence information. Antibodies were assigned to the same lineage if they shared identical heavy-chain V and J genes, identical light-chain V and J genes, identical heavy- and light-chain CDR3 lengths, and similar heavy- and light-chain CDR3 amino acid sequences. Candidate lineage assignments were subsequently confirmed by manual inspection of paired heavy- and light-chain sequence features to ensure consistent clonal relationships. Germline gene usage, SHM frequencies, heavy- and light-chain CDR3 length distributions, and clonal lineage relationships were analyzed for the isolated rhesus S2 stem-helix mAbs. Multiple sequence alignments of selected CDR3 amino acid sequences were generated using the MUSCLE algorithm (*93, 94*) implemented in the R package (*95*).

#### Statistical analysis

Statistical analysis was performed using GraphPad Prism 8, GraphPad Software, San Diego, California, USA, and R (v4.4.1). Descriptive data are presented as the median ± standard deviation or geometric mean ± standard deviation, as indicated in the figure legends. KDAPP, kon, koff, ED50 titters and percent weight change were compared using the Mann-Whitney test. Data were considered statistically significant when p < 0.05.

### Supplementary Figures and Tables

**Fig. S1.**
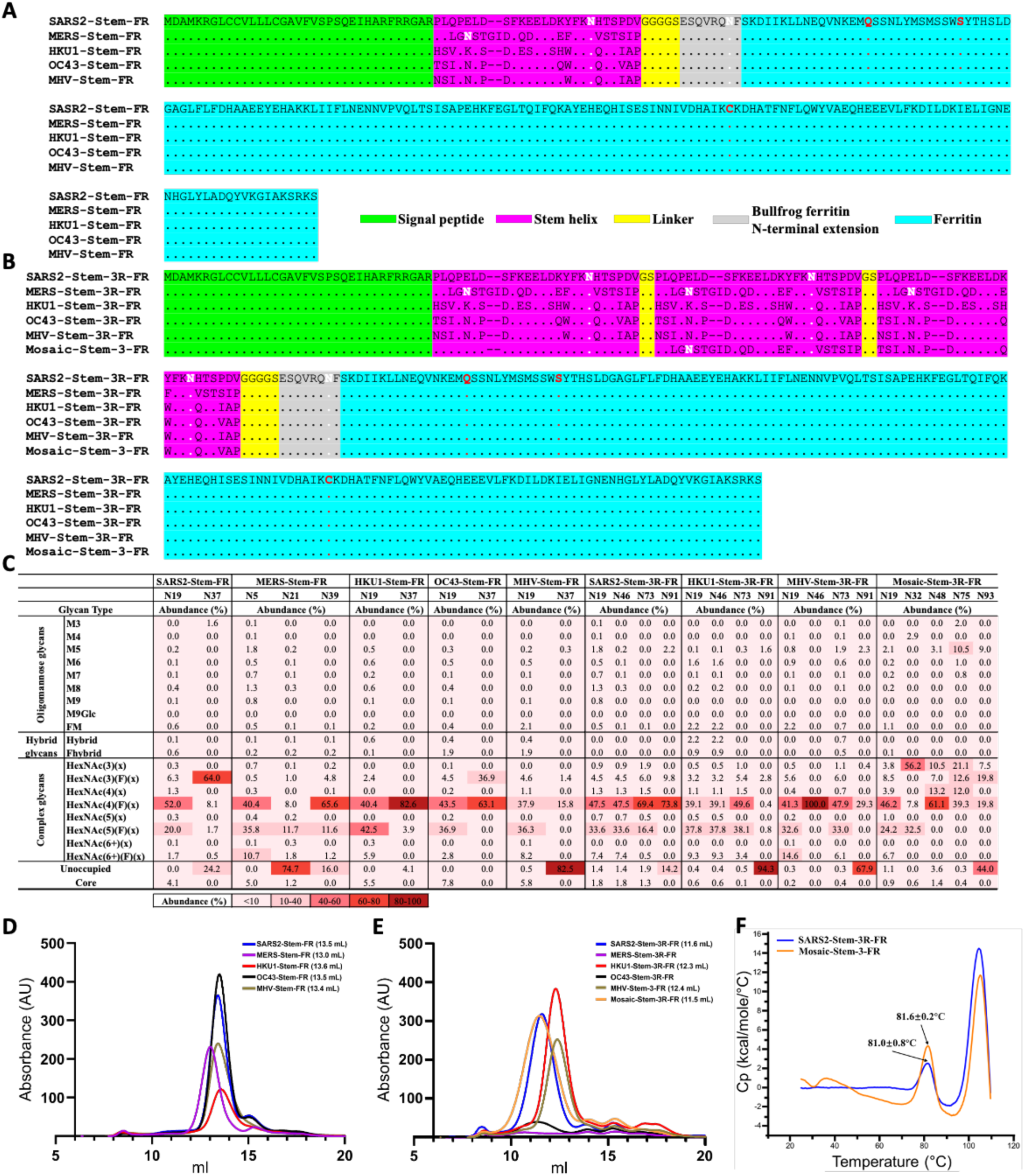
Sequence features and biophysical properties of stem-helix ferritin nanoparticles. (**A-B**) Amino acid sequences of stem-helix ferritin nanoparticles. The signal peptide, stem-helix peptide, linker, bullfrog ferritin N-terminal extension, and ferritin were shown in green, purple, yellow, grey, and cyan, respectively. Predicted N-linked glycosylation sites were highlighted in white. The bullfrog ferritin N-terminal extension was incorporated to provide outward-projecting and evenly distributed antigen attachment sites on the ferritin nanoparticle surface, and three mutations (highlighted in red, N19Q, C31S, and S111C) were introduced into the *Helicobacter pylori* ferritin scaffold to improve its functionality, as previously described (*36, 37*). Dots (.) represented identical amino acids, whereas dashes (-) indicated sequence insertions or deletions. The stem-helix peptide sequences shown here were derived from SARS-CoV-2 (SARS2), MERS-CoV (MERS), HCoV-HKU1 (HKU1), HCoV-OC43 (OC43), and murine hepatitis virus (MHV). SARS-CoV-1 shared an identical stem-helix sequence with SARS-CoV-2. Stem-FR displayed 24 copies of the stem-helix peptide per nanoparticle, whereas Stem-3R-FR displayed 72 copies. Mosaic-Stem-3-FR sequentially included stem-helix peptides from SARS-CoV-2, MERS-CoV, and HCoV-OC43, with 24 copies of each peptide displayed on the nanoparticle surface. FR, ferritin. (**C**) Site-specific N-linked glycan composition of the stem-helix ferritin nanoparticles. Glycans were classified as oligomannose glycans (M3–M9, M9Glc, and FM), hybrid glycans (including FHybrid), complex glycans, unoccupied (no glycan), or core glycans (truncated N-glycan structures smaller than M3). Values indicated the relative abundance (%) of each glycan species at the indicated glycosylation site. GNL preferentially bound high-mannose N-glycans (M5-M9) (*96, 97*). Glycosylation site numbering started from the first amino acid after the signal peptide in each nanoparticle construct. M9Glc, glucosylated oligomannose glycan; FM, fucosylated oligomannose glycan; FHybrid, fucosylated hybrid glycan. (**D-E**) Size-exclusion chromatography of stem-helix ferritin nanoparticles. Peak elution positions (mL) were indicated in parentheses. (**F**) Differential scanning calorimetry profiles of SARS2-Stem-3R-FR and Mosaic-Stem-3-FR. The average melting temperature (T_m_) ± SD from three independent measurements was shown for each nanoparticle.

**Fig S2.**
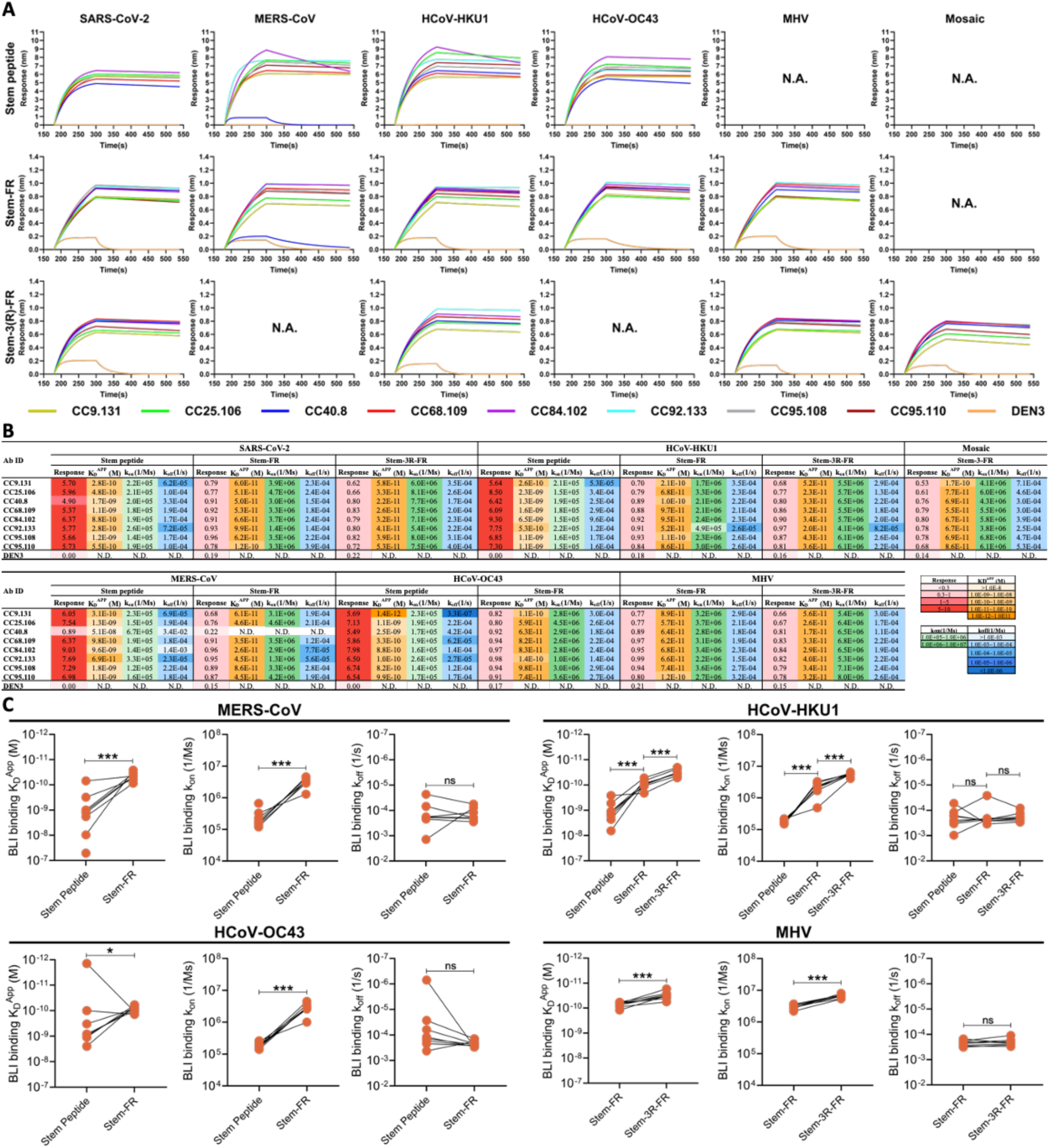
Antigenicity of stem-helix ferritin nanoparticles measured by biolayer interferometry (BLI). (**A**) BLI sensorgrams showing the binding of human S2 stem-helix bnAbs to the indicated synthetic betacoronavirus stem-helix peptides and stem-helix ferritin nanoparticles. Individual human S2 stem-helix bnAbs (IgG) were distinguished by color, and the negative control mAb DEN3 was shown in orange. (**B**) Maximum binding responses and kinetic parameters corresponding to the BLI sensorgrams shown in panel A. Binding kinetic parameters including apparent dissociation constant (K_D_^APP^), association rate constant (k_on_), and dissociation rate constant (k_off_) were calculated only for antibody-antigen interactions with a maximum binding response >0.3 nm. (**C**) Comparison of antibody binding kinetic parameters among monomeric stem-helix peptides, Stem-FR, and Stem-3R-FR within the same betacoronavirus. N.A., not available; N.D., not determined. *P* values were calculated by a Mann-Whitney test. ns, not significant; *p < 0.05; **p < 0.01; ***p < 0.001.

**Fig S3.**
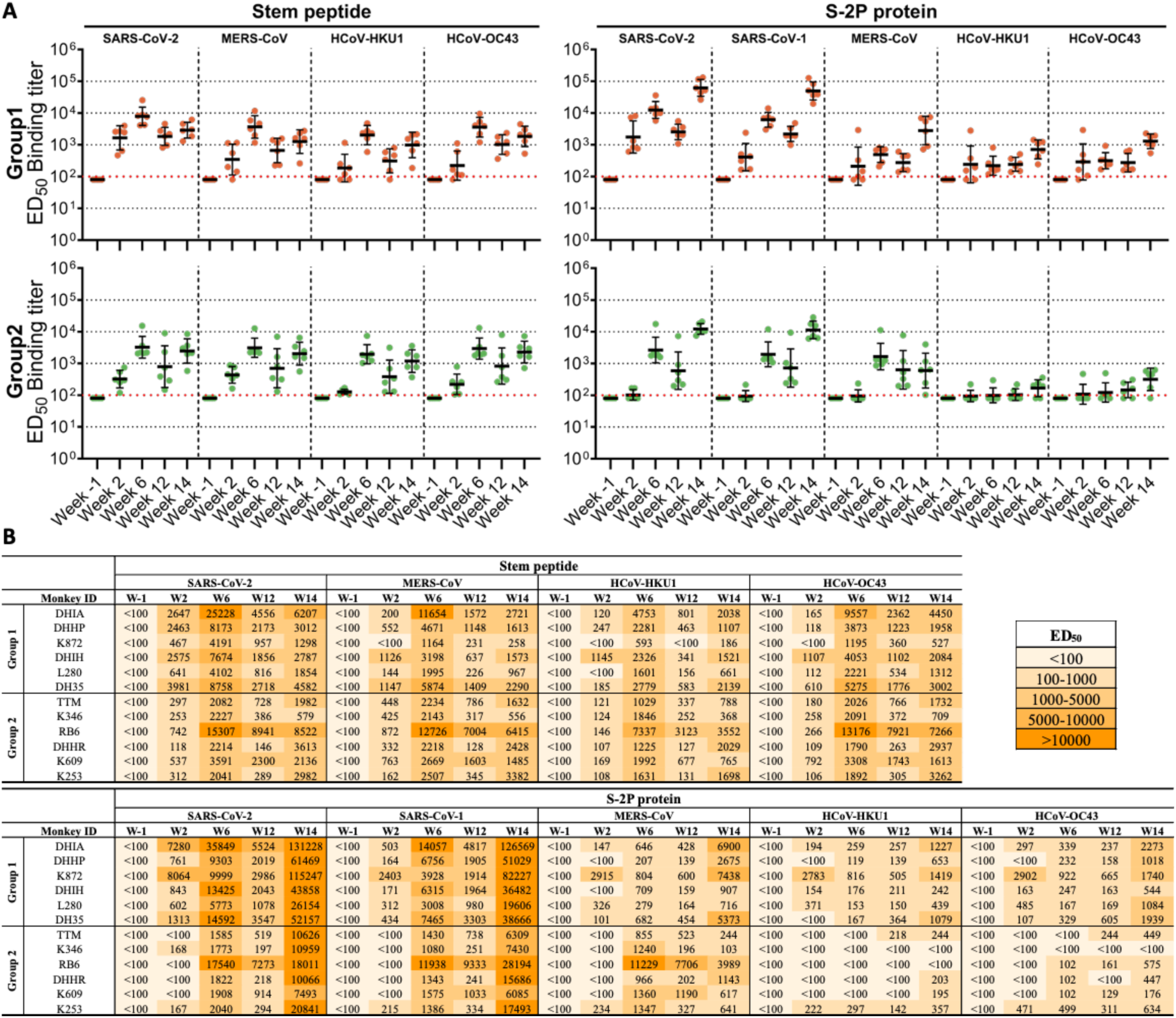
Plasma antibody binding in immunized macaques to human betacoronavirus S2 stem-helix peptides and S-2P proteins as a function of time. (**A**) Plasma binding antibody ED_50_ titers against stem-helix peptides (left) and S-2P proteins (right) from SARS-CoV-2, SARS-CoV-1, MERS-CoV, HCoV-HKU1, and HCoV-OC43 measured by ELISA. Plasma samples were collected from the immunized rhesus macaques shown in Fig. 2A. Macaques in Group 1 and Group 2 received two doses of SARS2-Stem-3R-FR or Mosaic-Stem-3-FR, respectively, followed by a single BA.1 spike mRNA booster. SARS-CoV-1 shared an identical stem-helix sequence with SARS-CoV-2. Each symbol represented one macaque, and the bars indicated geometric mean ED_50_ values ± SD. ED_50_ values <100, below the red dashed line, were considered negative for binding. ED_50_ values <100 were assigned a value of 80 for visualization. (**B**) Individual plasma binding antibody ED_50_ titers corresponding to the data shown in panel A. Cell colors ranged from light to dark orange according to the ED_50_ values, with darker shades indicating stronger binding. ED_50_, half-maximal effective dilution.

**Fig S4.**
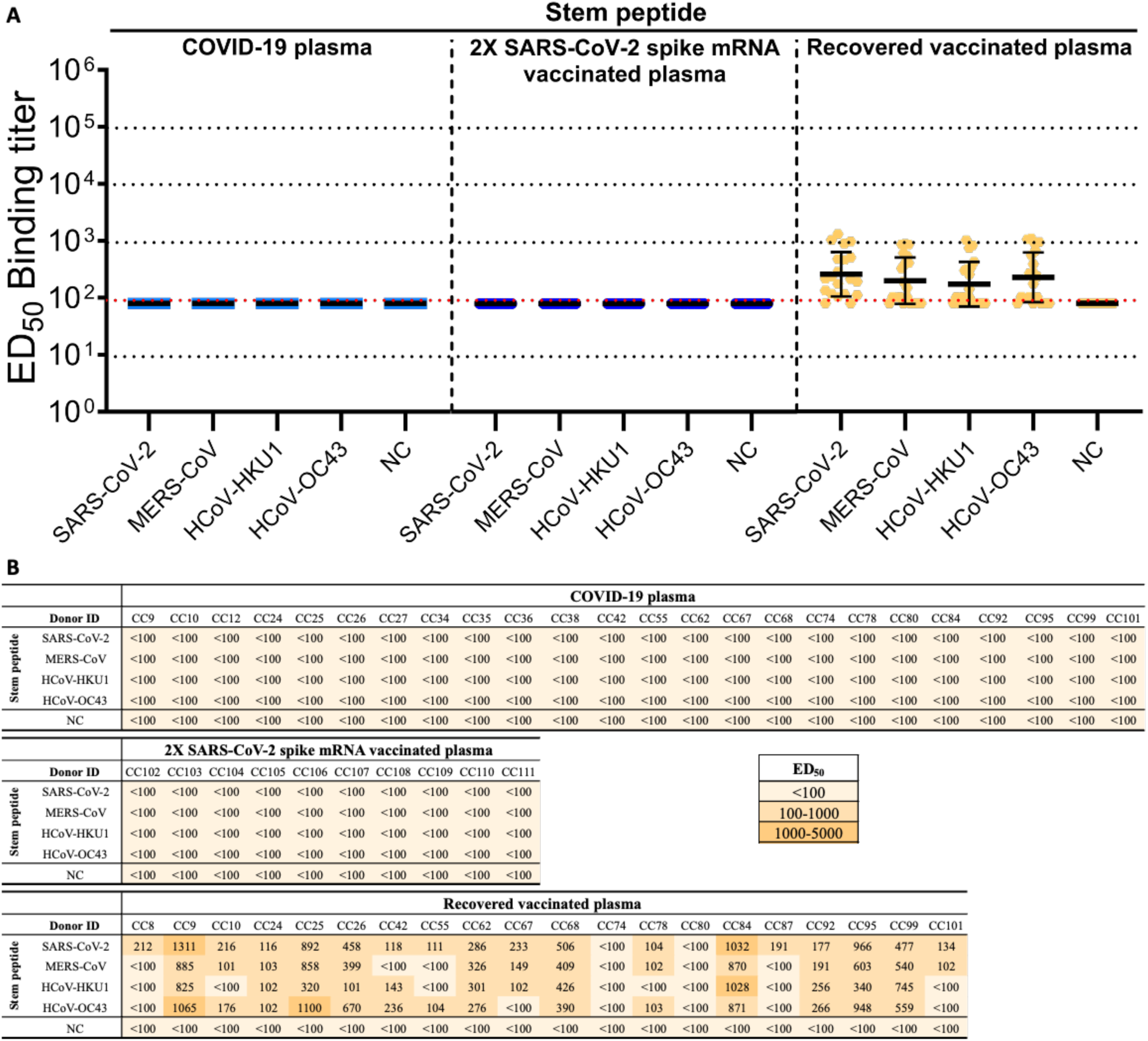
Plasma antibody binding in different human cohorts to human betacoronavirus S2 stem-helix peptides. (**A**) Plasma binding antibody ED_50_ titers against S2 stem-helix peptides from SARS-CoV-2, MERS-CoV, HCoV-HKU1, and HCoV-OC43 measured by ELISA. Plasma samples were obtained from three human cohorts: COVID-19 convalescent individuals, SARS-CoV-2 mRNA vaccine recipients, and recovered-vaccinated individuals. Recovered-vaccinated individuals referred to donors who recovered from SARS-CoV-2 infection and subsequently received SARS-CoV-2 mRNA vaccinationation. SARS-CoV-1 shared an identical stem-helix sequence with SARS-CoV-2. For the negative control (NC), ELISA plates were coated with streptavidin alone without addition of biotinylated stem-helix peptide, and all remaining steps were performed as described in the Methods section. Each symbol represented one donor, and the bars indicated geometric mean ED_50_ values ± SD. ED_50_ values <100, below the red dashed line, were considered negative for binding. ED_50_ values <100 were assigned a value of 80 for visualization. (**B**) Individual plasma binding antibody ED_50_ titers corresponding to the data shown in panel A. Cell colors ranged from light to dark orange according to the ED_50_ values, with darker shades indicating stronger binding. CC, COVID-19 cohort; 2X, two doses of SARS-CoV-2 mRNA vaccination; ED_50_, half-maximal effective dilution.

**Fig S5.**
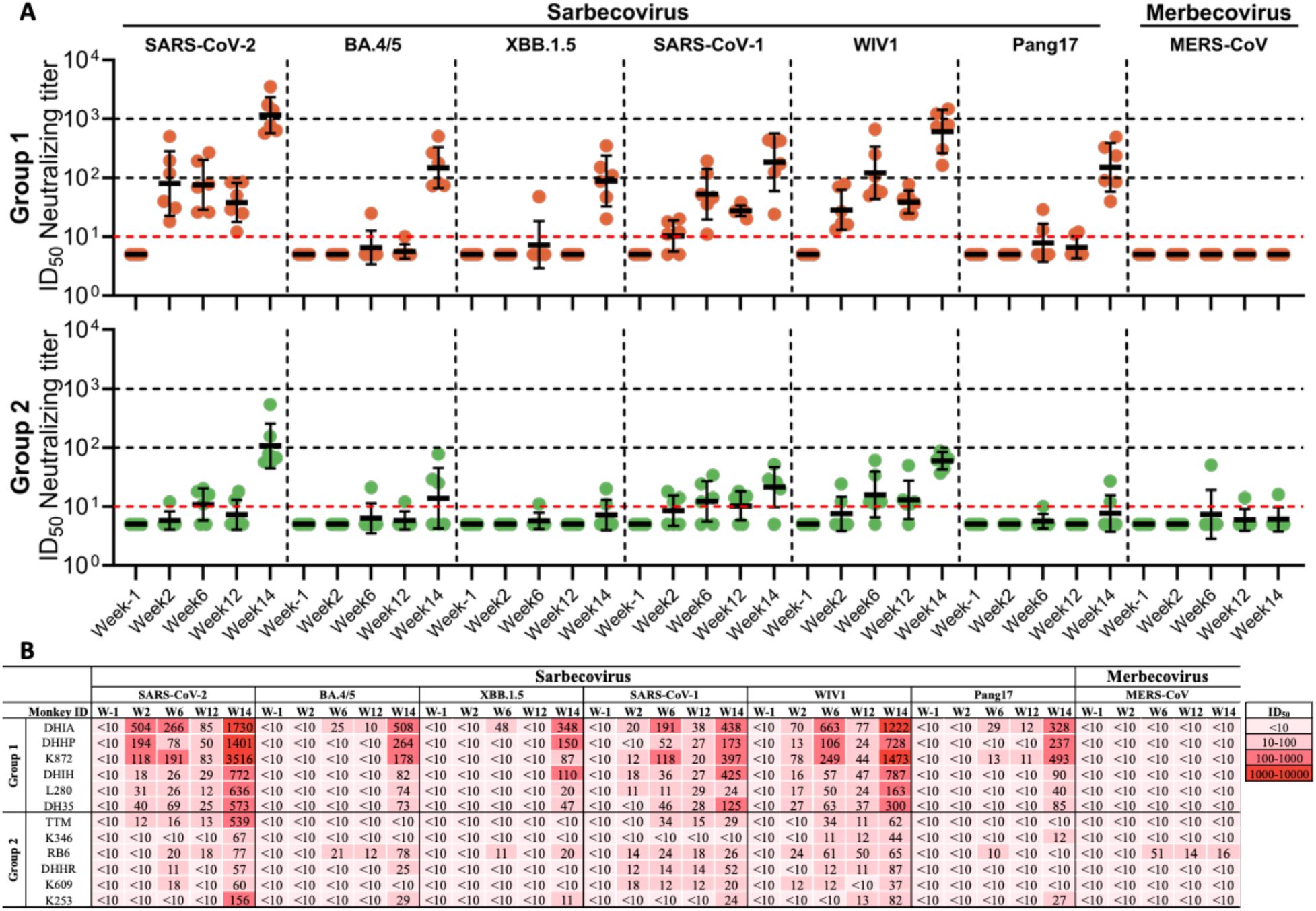
Plasma neutralizing antibody responses in immunized macaques against representative human betacoronavirus pseudoviruses as a function of time. (**A**) Plasma neutralizing antibody ID_50_ titers against representative human betacoronavirus pseudoviruses. SARS-CoV-2, its Omicron variants BA.4/5 and XBB.1.5, SARS-CoV-1, WIV1, and Pang17 were sarbecoviruses, whereas MERS-CoV was a merbecovirus. Plasma samples were collected at the indicated time points from the immunized macaques shown in Fig. 2A. Macaques in Group 1 and Group 2 received two doses of SARS2-Stem-3R-FR or Mosaic-Stem-3-FR, respectively, followed by a single BA.1 spike mRNA booster. Each symbol represented one macaque, and the bars indicated geometric mean ID_50_ values ± SD. ID_50_ values <10, below the red dashed line, were considered negative for neutralization. ID_50_ values <10 were assigned a value of 5 for visualization. (**B**) Individual plasma neutralizing antibody ID_50_ titers corresponding to the data shown in panel A. Cell colors ranged from light to dark red according to the ID_50_ values, with darker shades indicating higher neutralizing titers. ID_50_, 50% inhibitory dilution.

**Fig S6.**
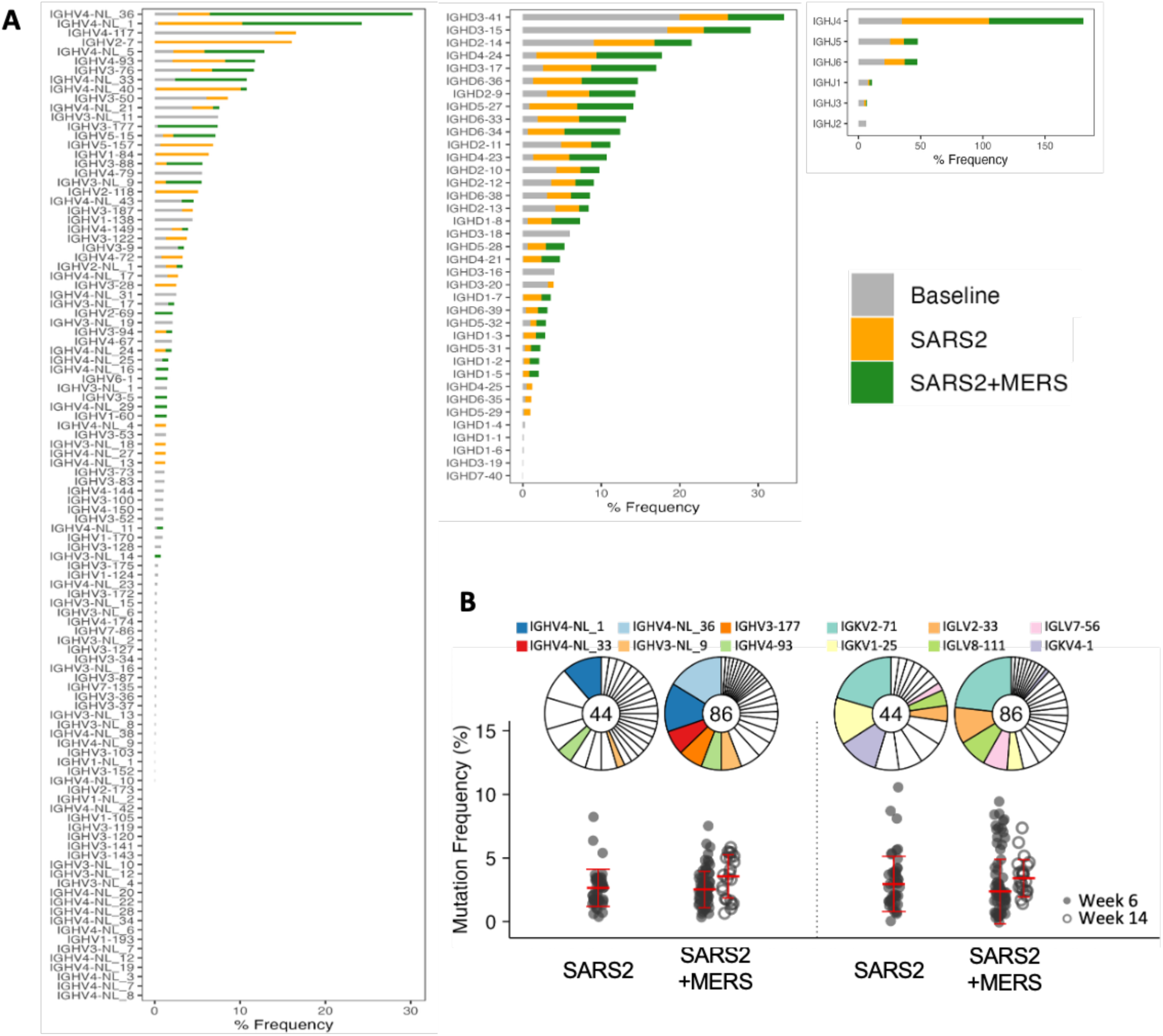
Germline gene usage and somatic hypermutations in 130 isolated rhesus S2 stem-helix mAbs. (**A**) Germline gene usage distribution of heavy chains among isolated rhesus S2 stem-helix mAbs grouped by binding specificity to spike proteins. SARS-CoV-2 spike-specific binding mAbs (binding to SARS-CoV-2 but not MERS-CoV spike, SARS2, n = 78) and mAbs with cross-binding to both SARS-CoV-2 and MERS-CoV spikes (SARS2+MERS, n = 52) were shown in orange and green, respectively. All mAbs in the SARS2 group bound the SARS-CoV-2 S2 stem-helix peptide, and all mAbs in the SARS2+MERS group also cross-bound the SARS-CoV-2 and MERS-CoV S2 stem-helix peptides. Baseline germline frequencies of rhesus heavy chain genes (IGHV, IGHD and IGHJ) were shown in grey. (**B**) Germline gene usage distribution and somatic hypermutation levels among isolated rhesus S2 stem-helix mAbs grouped by binding specificity to S2 stem-helix peptides after homologous nanoparticle boost (week 6, closed dots) and BA.1 spike boos (week 14, open dots). Dot plots showed the percentage of nucleotide somatic hypermutations (SHMs) in the heavy and light chain variable regions. Pie chart segments were ordered counterclockwise by V gene usage frequency, from highest to lowest. Median ± SD was shown. The SARS2 group comprised 44 mAbs that bound the SARS-CoV-2 but not MERS-CoV S2 stem-helix peptide. The SARS2+MERS group comprised 86 mAbs that cross-bound both the SARS-CoV-2 and MERS-CoV S2 stem-helix peptides, including all 52 mAbs in the SARS2+MERS group shown in panel A.

**Fig S7.**
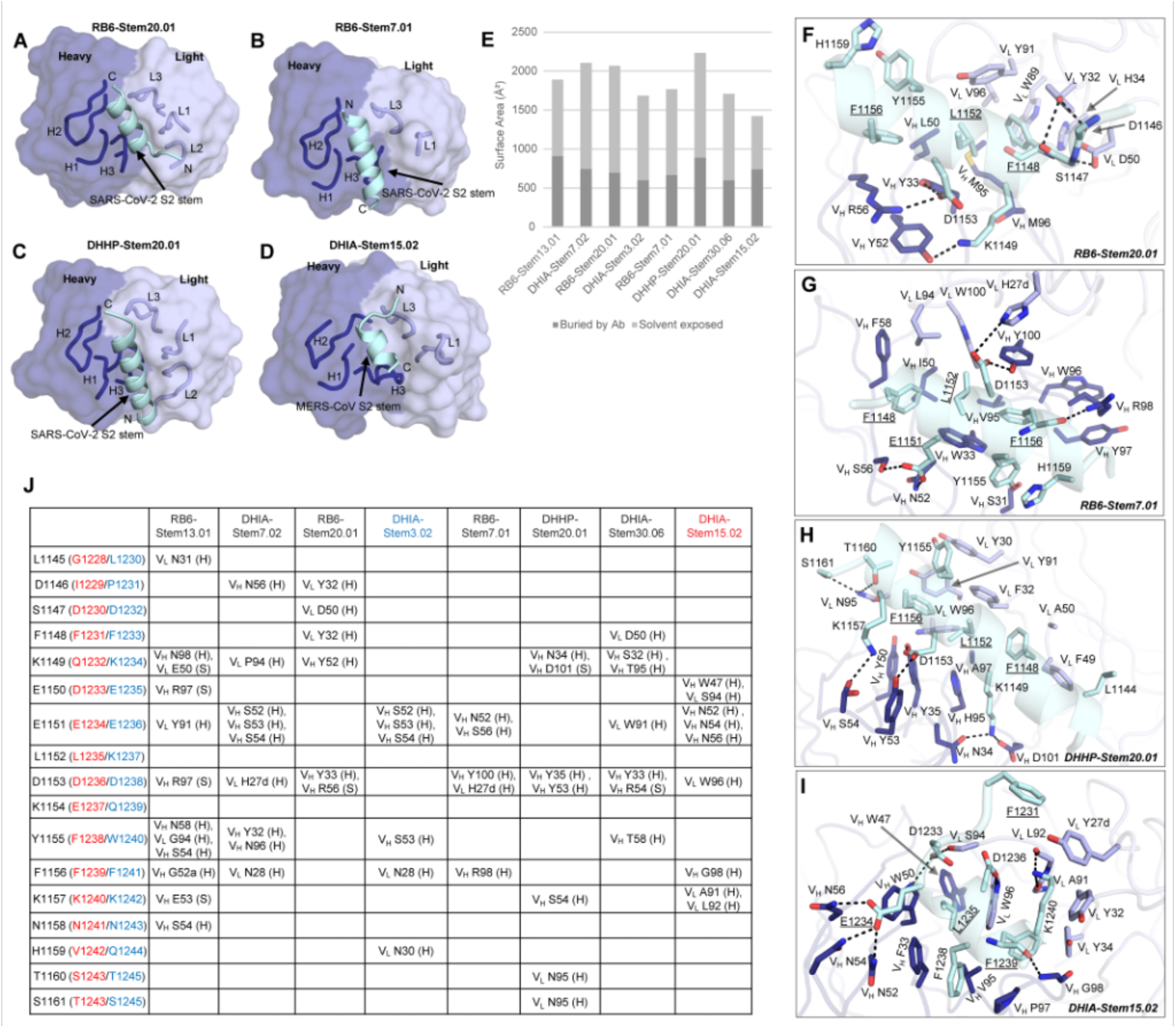
Crystal structures and detailed atomic interactions of rhesus antibodies in complex with S2 stem-helix peptides. The color scheme and numbering were as in Figure 5. (**A-D**) Overall view of the crystal structures of complexes of the RB6-Stem20.01-SARS-CoV-2 S2 stem-helix peptide, RB6-Stem7.01-SARS-CoV-2 S2 stem-helix peptide, DHHP-Stem20.01-SARS-CoV-2 S2 stem-helix peptide, and DHIA-Stem15.02-MERS-CoV S2 stem-helix peptide structures at 2.28 Å, 1.98 Å, 1.73 Å, and 1.68 Å resolutions, respectively. CDRH1, H2, H3, L1, L2, L3 were shown in ribbon representation and labeled. The N- and C-terminus of the peptides were labeled N and C, respectively. (**E**) The surface area of S2 stem-helix peptides buried by the eight antibodies. Solvent exposed and buried areas were calculated with Proteins, Interfaces, Structures, and Assemblies (PISA) (*85*). (**F-I**) Detailed atomic interactions between the S2 stem-helix peptides and rhesus antibodies: (**F**) RB6-Stem20.01 with SARS-CoV-2 S2 stem-helix peptide, (**G**) RB6-Stem7.01 with SARS-CoV-2 S2 stem-helix peptide, (**H**) DHHP-Stem20.01 with SARS-CoV-2 S2 stem-helix peptide, and (**I**) DHIA-Stem15.02 with MERS-CoV S2 stem-helix peptide. (**J**) Hydrogen bonds and salt bridges identified between the S2 stem helices and the eight antibodies using the PISA program. “H” represented a hydrogen bond and “S” was a salt bridge/hydrogen bond. Kabat numbering was used for the antibodies. Numbering of the peptides was based on SARS-CoV-2(red and blue numbers in brackets corresponding to MERS-CoV and HCoV-OC43 numbering, respectively). Interactions were identified by accessing PISA at the European Bioinformatics Institute (http://www.ebi.ac.uk/pdbe/prot_int/pistart.html)(98).

**Fig S8.**
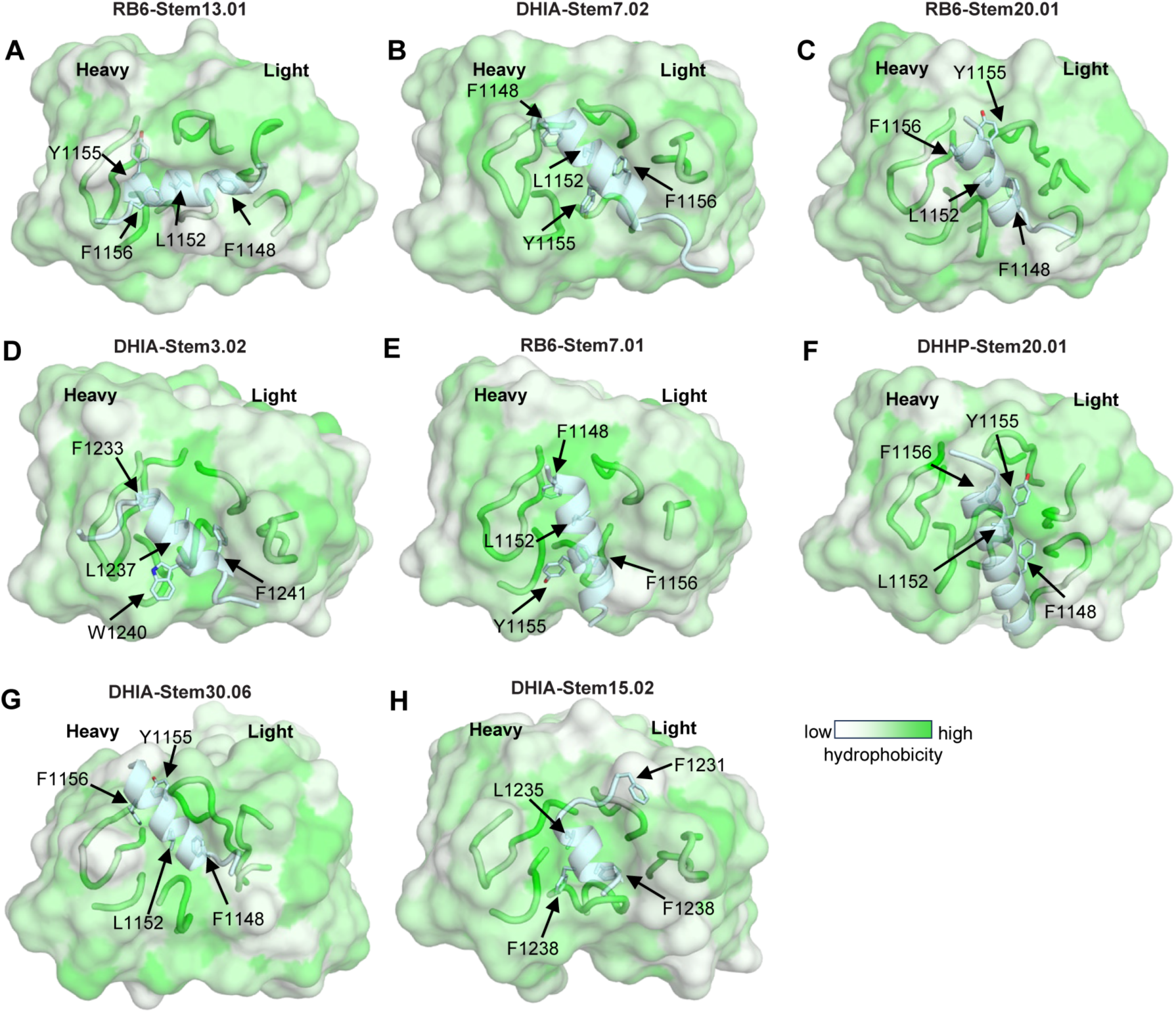
Stem-helix peptides bound in hydrophobic grooves formed by heavy and light chains of rhesus S2 stem-helix mAbs. The surfaces of the eight antibodies are color-coded by hydrophobicity [calculated using Color h (https://pymolwiki.org/index.php/Color_h)]. Four aromatic and hydrophobic residues of the S2 stem-helix peptides were shown.

**Fig S9.**
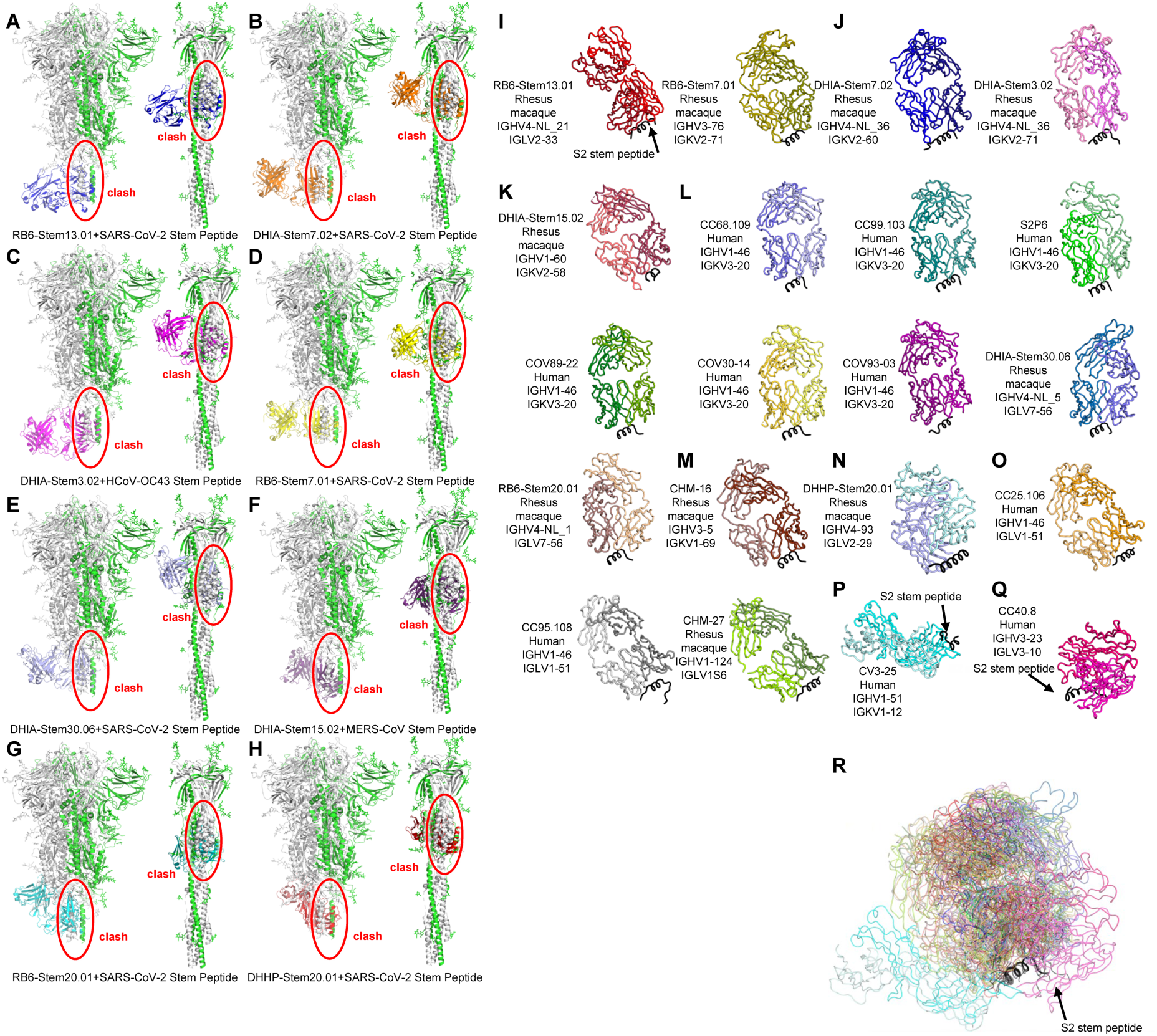
Steric clashes of rhesus S2 stem-helix mAbs with spike protein in the pre- and post-fusion states, and structural alignment of 20 human and rhesus S2 stem-helix mAbs. (A-H) Crystal structures of eight rhesus antibody-S2 stem-helix peptide complexes superimposed on SARS-CoV-2 spike structure in its pre-fusion state (PDB: 6XR8) and post-fusion state (PDB: 6XRA). Superimposed protomers were shown in green, while the other two protomers of the spike trimer were in grey. Steric clashes were indicated by red circles. (**I-R**) Crystal structures of the rhesus and human S2 stem-helix mAb Fabs bound to stem-helix peptides. All S2 stem helices were shown in black in the same orientation. Source and V_H_/V_L_ germline genes of each antibody were listed. For clarity, the heavy chains of all antibodies were shown in a darker color than the light chains. (**I**) RB6-Stem13.01-peptide complex and RB6-Stem7.01-peptide complex (this study).(**J**) DHIA-Stem7.02-peptide complex and DHIA-Stem3.02-peptide complex (this study). These two antibodies shared a nearly identical angle of binding to the S2 stem-helix peptide. (**K**) DHIA-Stem15.02-peptide complex (this study). (**L**) S2 stem-helix peptides in complex with human antibodies CC68.109 (PDB: 8DGX), CC99.103 (PDB: 8DGV), S2P6 (PDB: 7RNJ), COV89-22 (PDB: 8DTX), COV30-14 (PDB: 8DTR), COV93-03 (PDB: 8DTT), and rhesus antibodies RB6-Stem20.01 (this study) and DHIA-Stem30.06 (this study). These eight antibodies shared a nearly identical angle of binding to the S2 stem-helix peptide. (**M**) CHM-16-peptide complex (PDB: 8TMY). (**N**) DHHP-Stem20.01-peptide complex (this study). (**O**) S2 stem-helix peptides in complex with human antibodies CC25.106 (PDB: 8DGU), CC95.108 (PDB: 8DGW), and rhesus antibody CHM-27 (PDB: 8TMZ). These three antibodies shared a nearly identical angle of binding to the stem-helix peptide. (**P**) Human antibody CV3-25-peptide complex (PDB: 7RAQ). (**Q**) Human antibody CC40.8-peptide complex (PDB: 7SJS). (**R**) Structural alignment of the 20 antibody-peptide complexes superimposed on their S2 stem helices illustrating divergence in the antibody approach to the stem-helix peptide. S2 stem helices were shown in black and antibody colors were the same as in the individual structures above.

**Fig S10.**
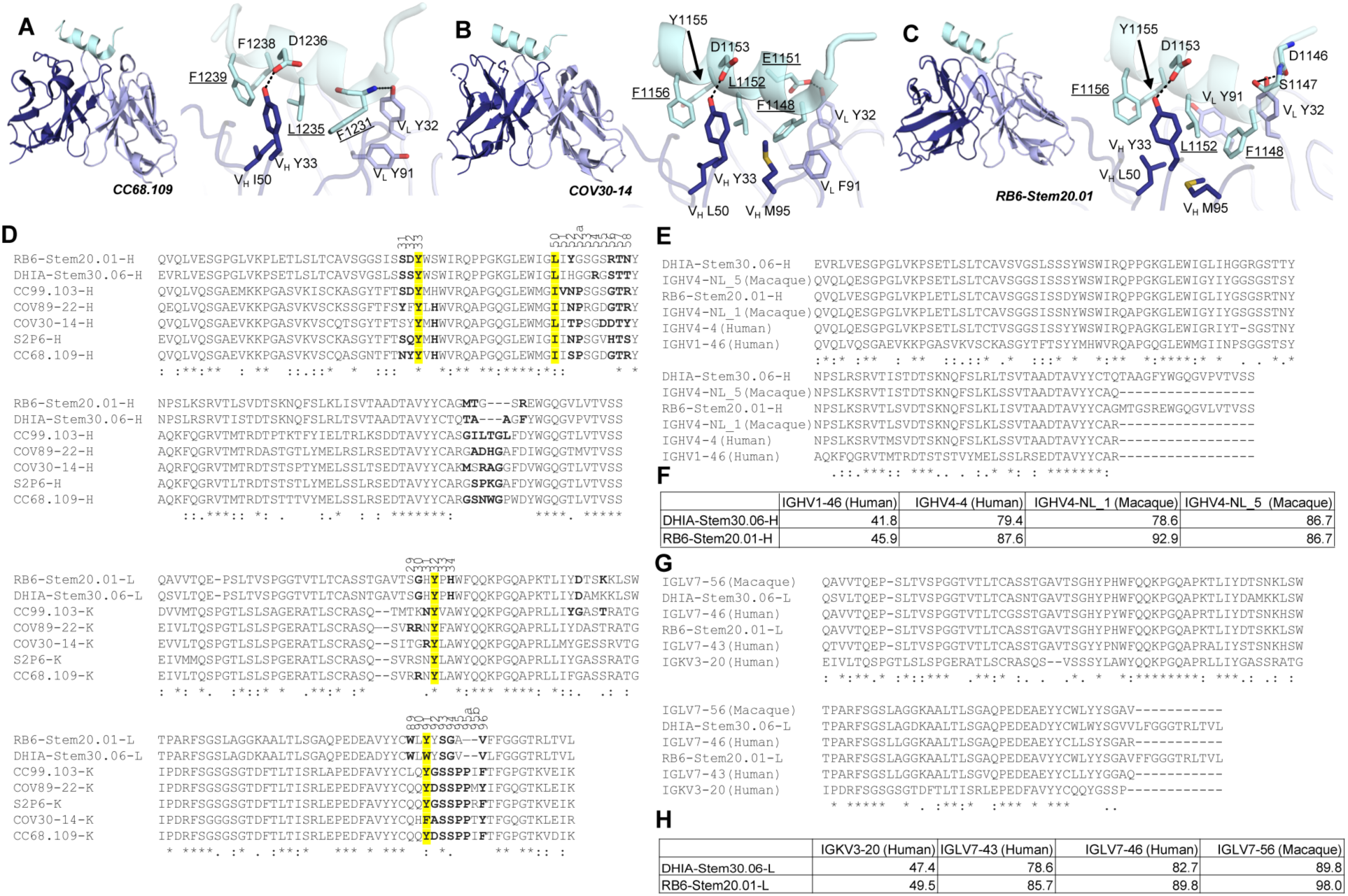
Structural and sequence alignment of RB6-Stem20.01-peptide and DHIA-Stem30.06-peptide with five human IGHV1-46/IGKV3-20 antibodies in complex with S2 stem-helix peptides. The color scheme and numbering were the same as in Figure 5. Crystal structures of: (**A**) human antibody CC68.109 with MERS-CoV S2 stem-helix peptide (PDB: 8DGX); (**B**) human antibody COV30-14 with SARS-CoV-2 S2 stem-helix peptide (PDB: 8DTR); (**C**) rhesus antibody RB6-Stem20.01 with SARS-CoV-2 S2 stem-helix peptide. Detailed interactions of conserved residues in antibodies and S2 stem-helix peptides were shown on the right of each antibody panel. Conserved identical residues between betacoronaviruses were underlined. Hydrogen bonds were represented by black dashed lines. (**D**) Sequence alignment of rhesus RB6-Stem20.01 and DHIA-Stem30.06 with five human IGHV1-46/IGKV3-20 antibodies. Residues that were involved in antigen interaction (BSA > 0 Å^2^ as calculated by PISA(*98*)) were in bold. Conserved or similar paratope residues that were nearly identical in all antibodies were highlighted by yellow boxes. Residues identical in all aligned sequences were labeled by an asterisk (*), whereas a colon (:) and a period (.) indicated strongly similar and less similar sequences, respectively. The sequence alignment was performed with Clustal Omega(*99*). Residue numbers of CDR loops (Kabat numbering) were shown on top of the sequences. (**E-H**) Sequence alignment of RB6-Stem20.01 and DHIA-Stem30.06 with their putative rhesus germline gene sequences, and comparison to the most similar human germlines including IGHV1-46/IGKV3-20. (**E-F**) Sequence alignment of two rhesus antibodies DHIA-Stem30.06 and RB6-Stem20.01 heavy chains with their putative heavy chain germline genes, most similar human germline genes, and the human IGHV1-46 germline gene. (**G-H**) Sequence alignment of two rhesus antibodies DHIA-Stem30.06 and RB6-Stem20.01 light chains with their putative light chain germline genes, most similar human germline genes, and the human IGKV3-20 germline gene. Putative germline sequences analyses were conducted by searching the IMGT database(*100*) with IgBLAST (*101*). The sequence alignments were performed with Clustal Omega(*99*), and amino-acid sequence identity percentages were shown in the tables below the sequences.

**Fig S11.**
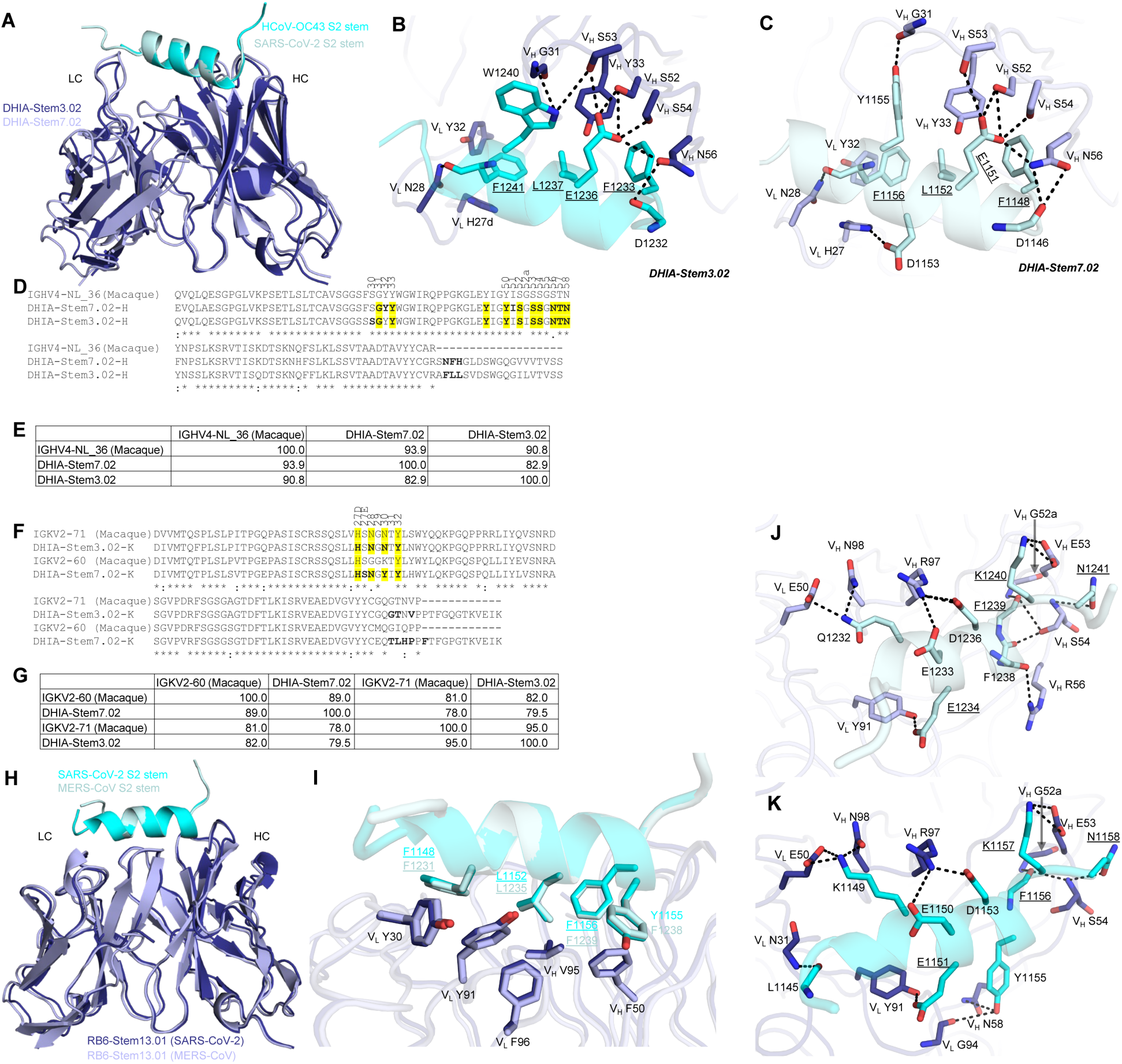
Structural and sequence comparison of rhesus antibodies DHIA-Stem7.02 and DHIA-Stem3.02, and structural comparison of rhesus antibody RB6-Stem13.01 bound to SARS-CoV-2 and MERS-CoV stem-helix peptides. (**A**) Superimposition of the DHIA-Stem7.02-SARS-CoV-2 S2 stem-helix peptide complex and the DHIA-Stem3.02-HCoV-OC43 S2 stem-helix peptide complex, with the S2 stem-helix peptide as reference. The HCoV-OC43 S2 stem-helix peptide was shown in cyan, whereas the DHIA-Stem3.02 was in blue. The SARS-CoV-2 S2 stem-helix peptide was shown in pale cyan, whereas the DHIA-Stem7.02 was in lavender. For clarity, only the variable domains of the Fabs were shown. (**B-C**) Detailed interactions of conserved paratope residues of (**B**) DHIA-Stem7.02 and (**C**) DHIA-Stem3.02 with HCoV-OC43 S2 stem-helix peptide and SARS-CoV-2 S2 stem-helix peptide. Conserved identical residues between betacoronaviruses were underlined. Hydrogen bonds were represented by black dashed lines. (**D-E**) Sequence alignment of the DHIA-Stem7.02 and DHIA-Stem3.02 heavy chains with their putative heavy chain germline genes. (**F-G)** Sequence alignment of the DHIA-Stem7.02 and DHIA-Stem3.02 light chains with their putative light chain germline gene. Putative germline sequence analyses of the heavy and light chains were conducted by searching the IMGT database(*100*) with IgBLAST(*101*). Residues that were involved in interacting with the antigens (BSA > 0 Å^2^ as calculated by PISA(*98*)) were in bold. Paratope residues that were identical in DHIA-Stem7.02 and DHIA-Stem3.02 were highlighted by yellow boxes. Residues identical in all aligned sequences were labeled by an asterisk (*), whereas a colon (:) and a period (.) indicated strongly similar and less similar sequences, respectively. The sequence alignment was performed with Clustal Omega(*99*), where amino-acid sequence identity percentages were shown in the tables below the sequences. Residue numbers of CDR loops (Kabat numbering) were shown on top of the sequences. (**H-K**) The SARS-CoV-2 S2 stem-helix peptide was shown in cyan, whereas the Fab was in blue. The MERS-CoV S2 stem-helix peptide was shown in pale cyan, whereas the Fab was in lavender. **(H)** Superimposition of the RB6-Stem13.01-SARS-CoV-2 S2 stem-helix peptide complex and the RB6-Stem13.01-MERS-CoV S2 stem-helix peptide complex, with the S2 stem-helix peptide as reference. For clarity, only the variable domains of the Fabs were shown. **(I)** Hydrophobic interactions between RB6-Stem13.01 with MERS-CoV S2 stem-helix peptide and SARS-CoV-2 S2 stem-helix peptide. Four aromatic and hydrophobic residues of S2 stem-helix peptides were shown. **(J-K)** Detailed interactions of paratope residues of RB6-Stem13.01 with (**J**) MERS-CoV S2 stem-helix peptide and (**K**) SARS-CoV-2 S2 stem-helix peptide. Conserved identical residues between betacoronaviruses were underlined. Hydrogen bonds and salt bridges were represented by black dashed lines.

**Table S1.**
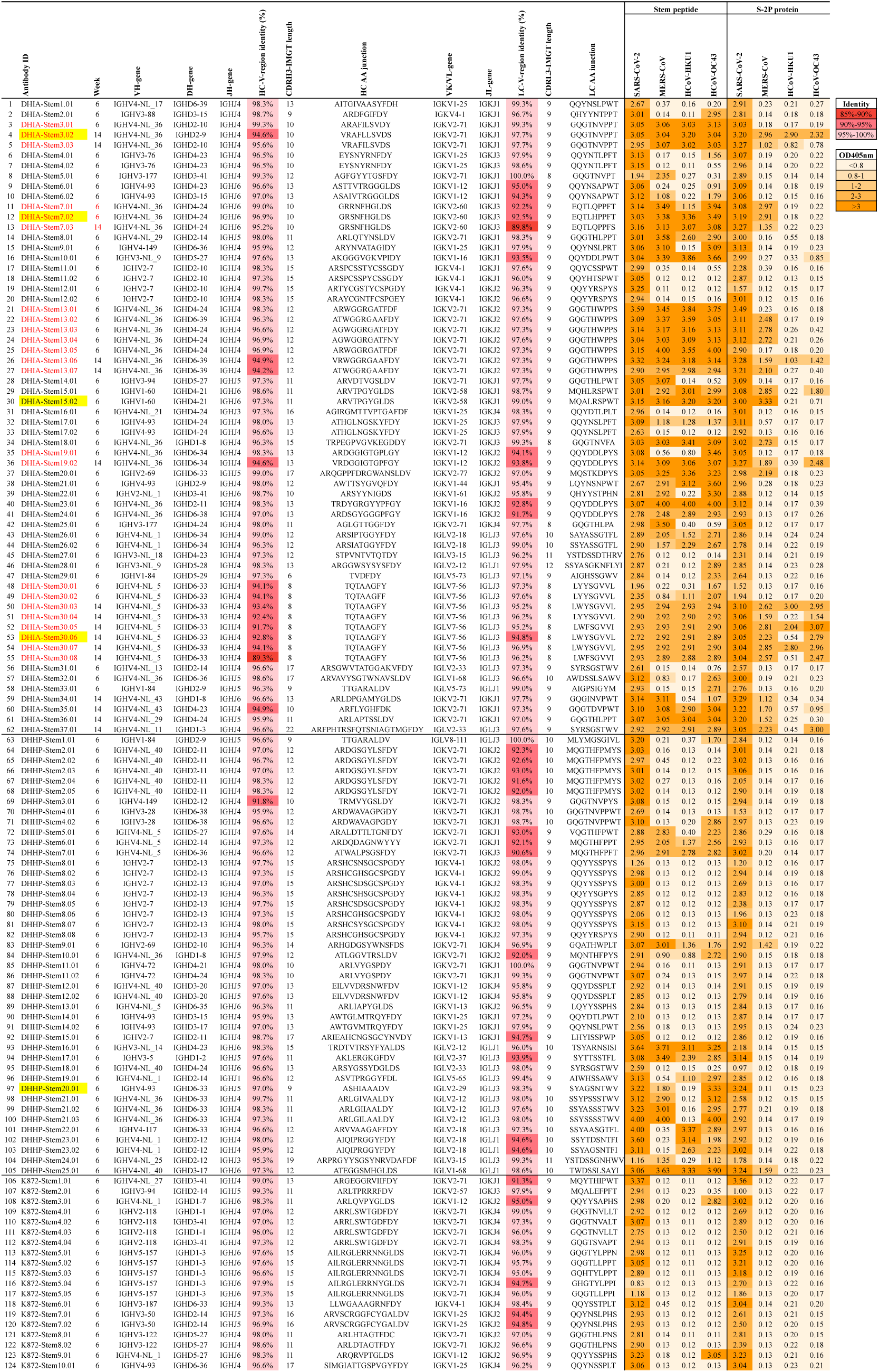
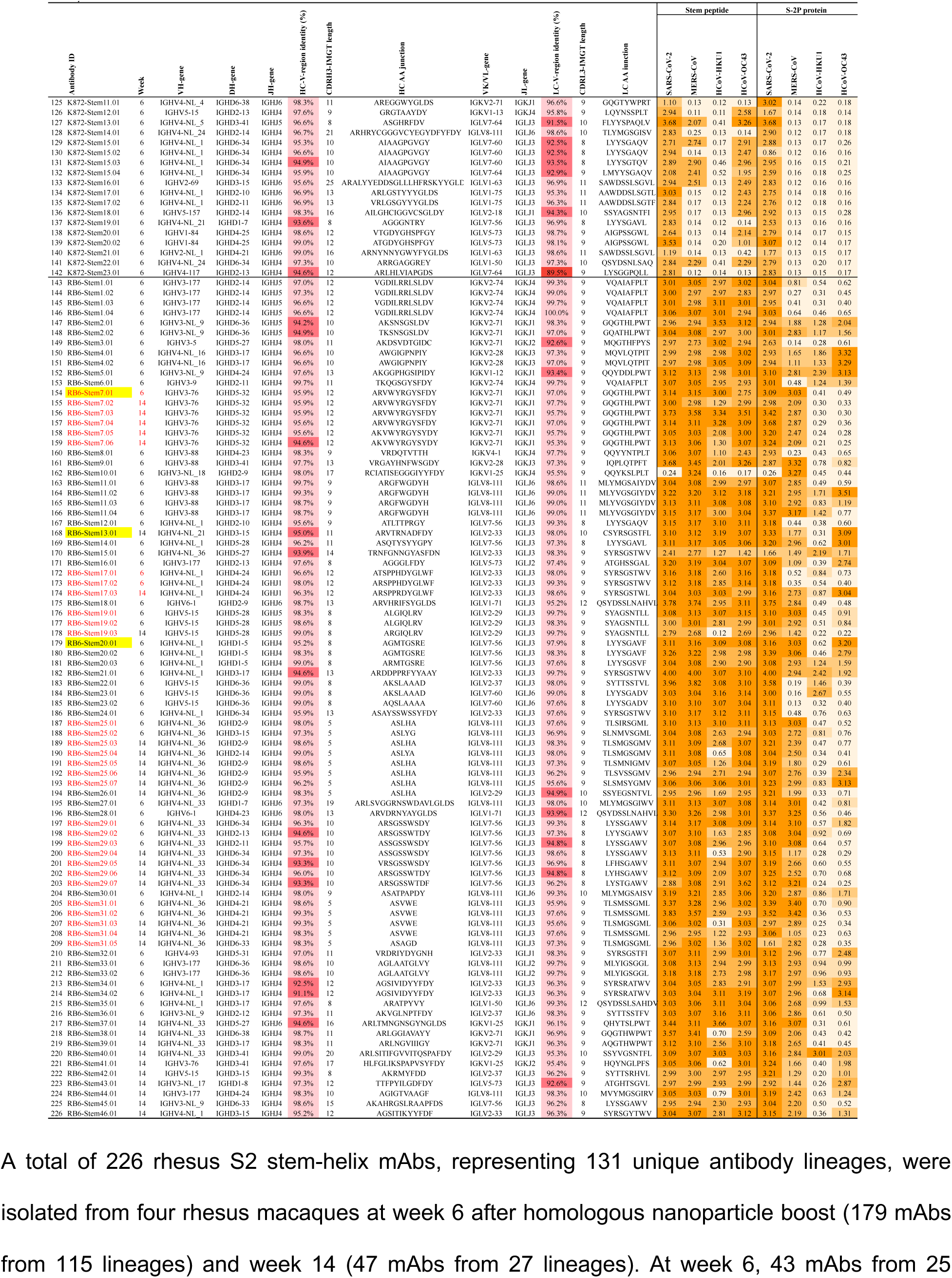
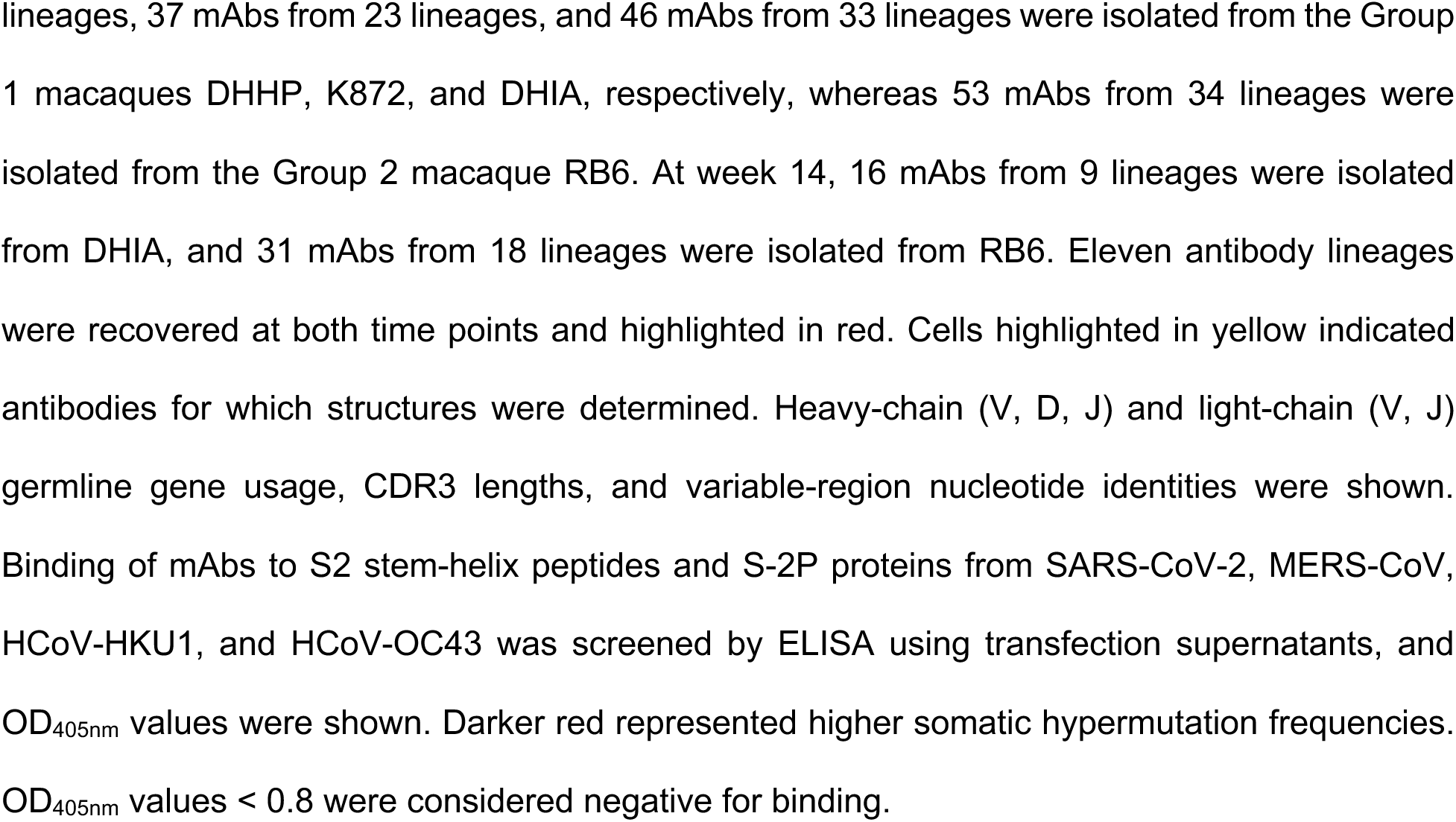
Immunogenetic and binding properties of isolated rhesus S2 stem-helix mAbs.

**Table S2.**
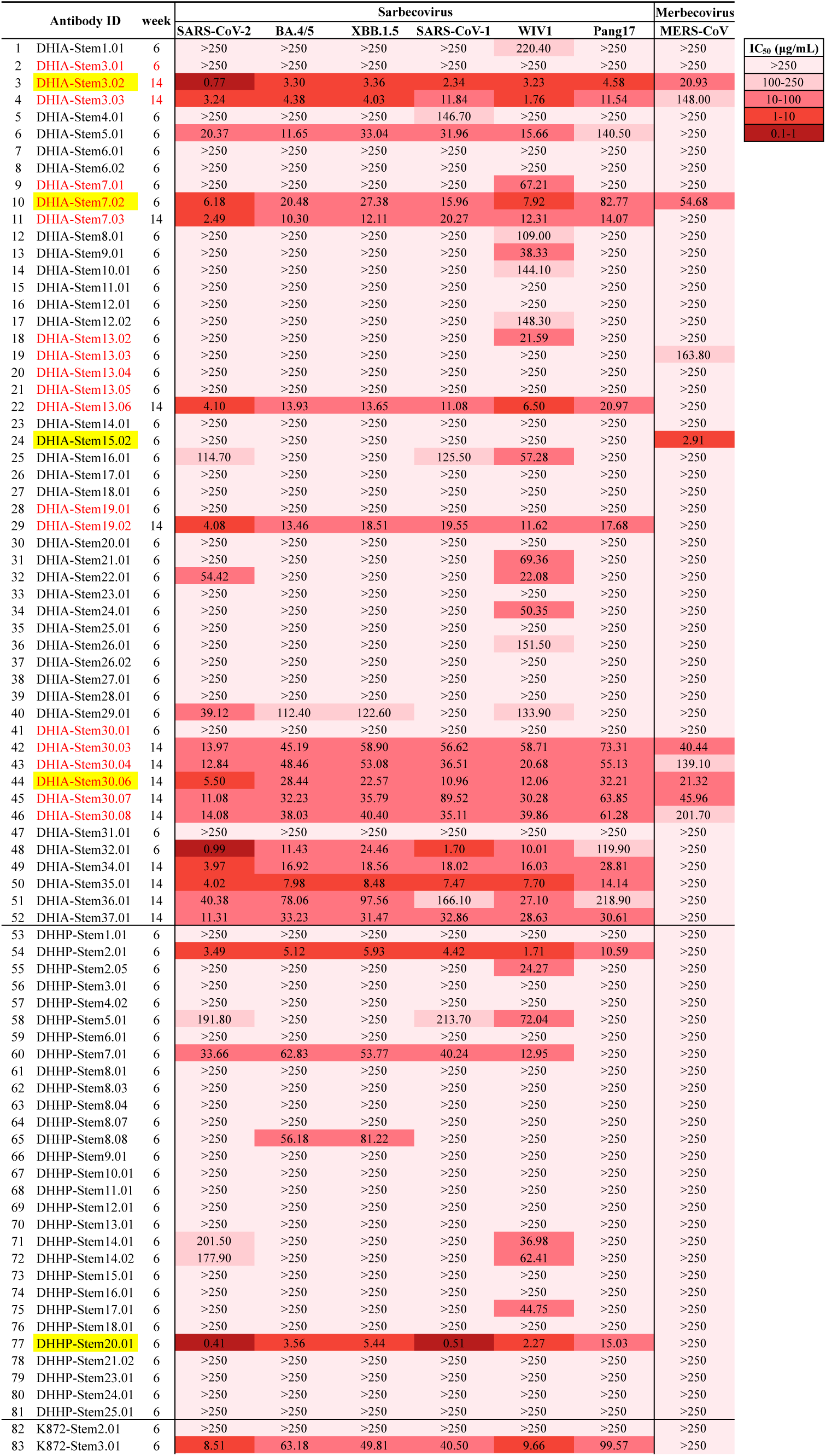
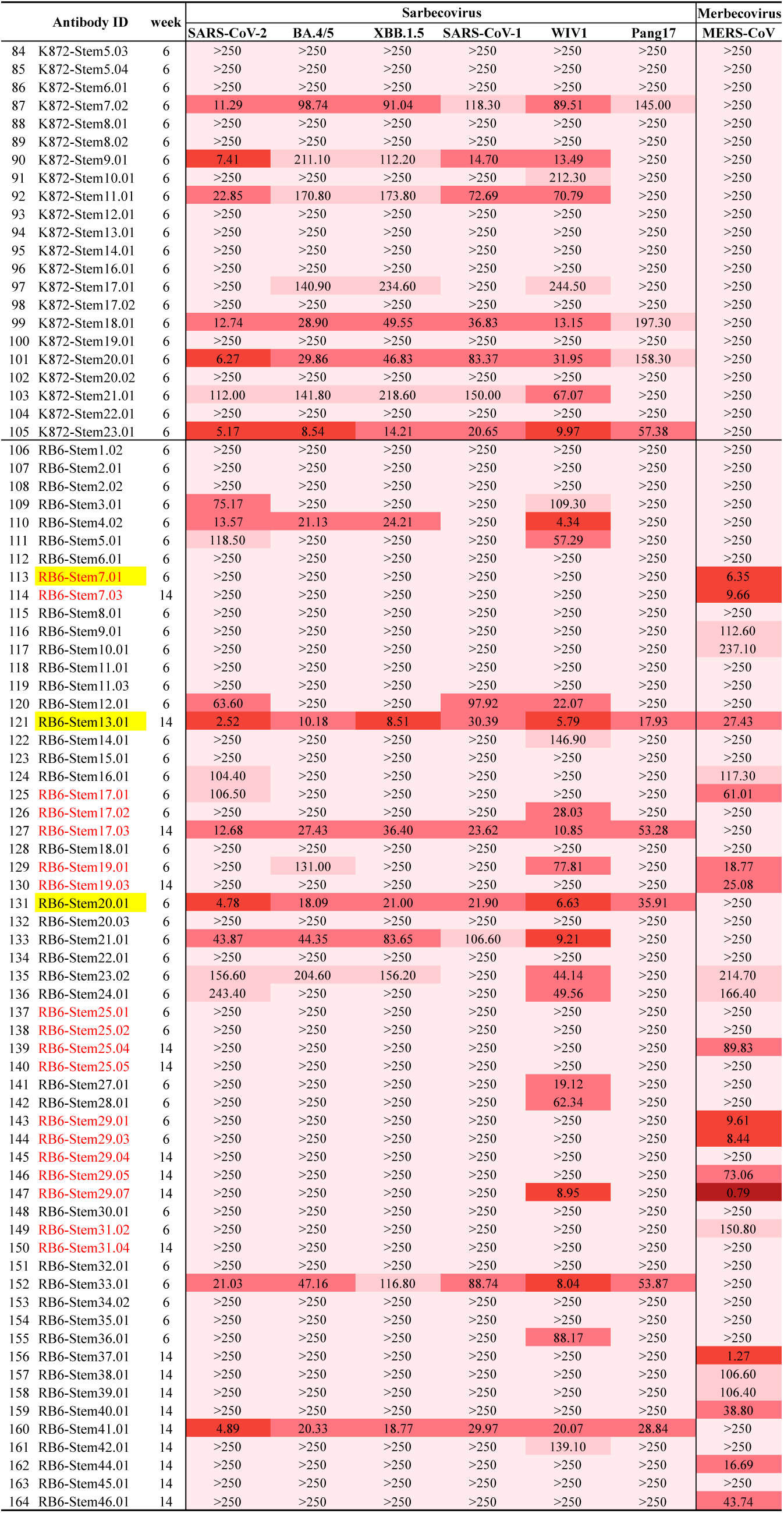
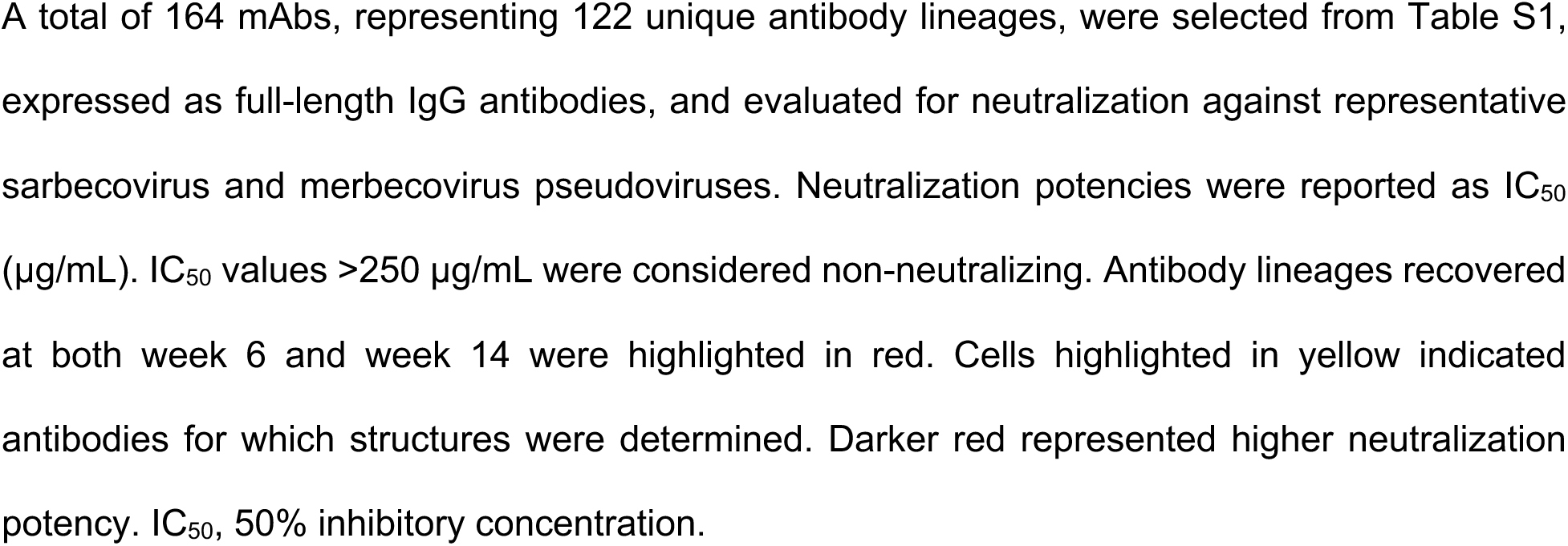
Neutralization properties of selected rhesus S2 stem-helix mAbs.

**Table S3.**
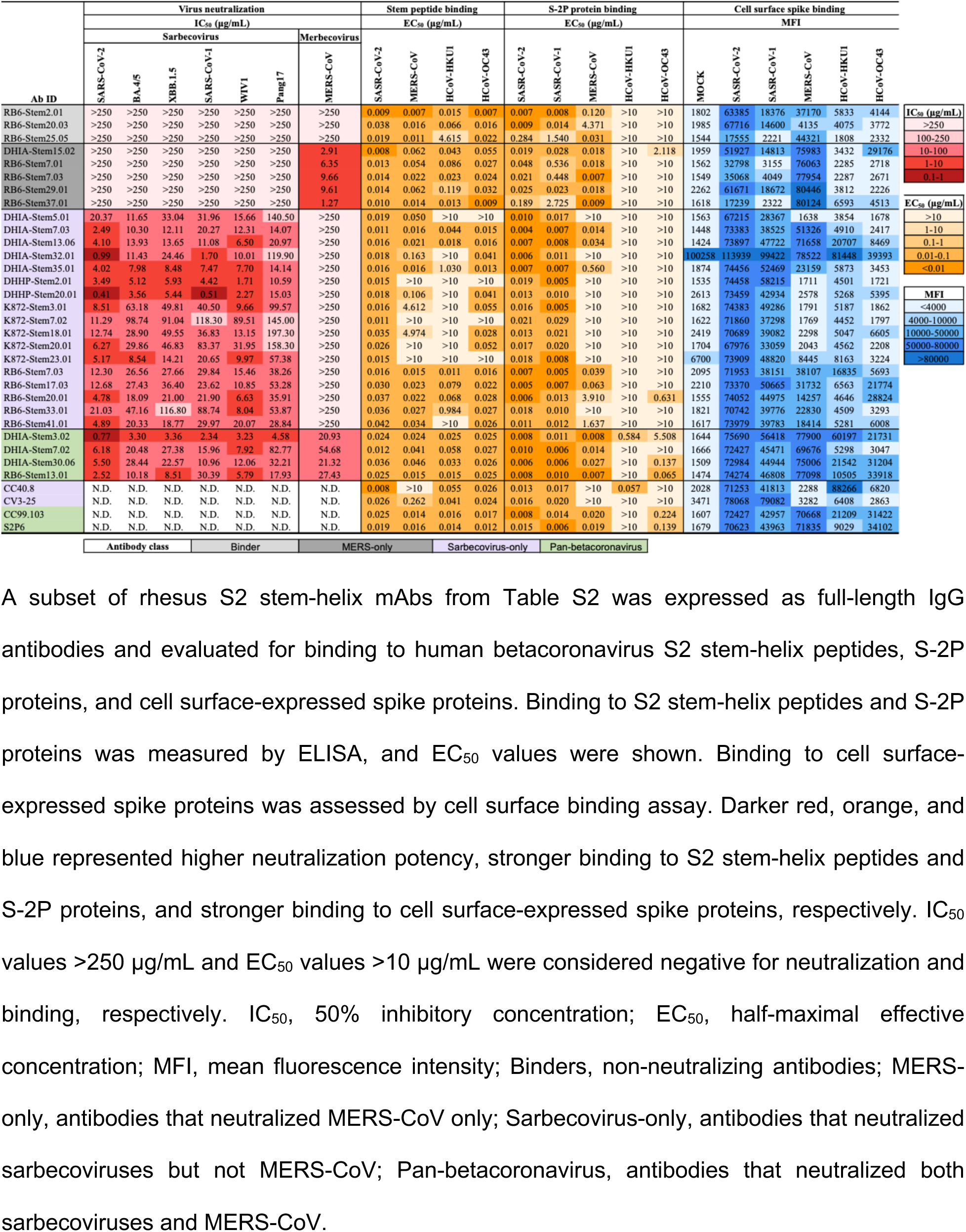
Binding characterization of selected rhesus S2 stem-helix mAbs.

**Table S4.**
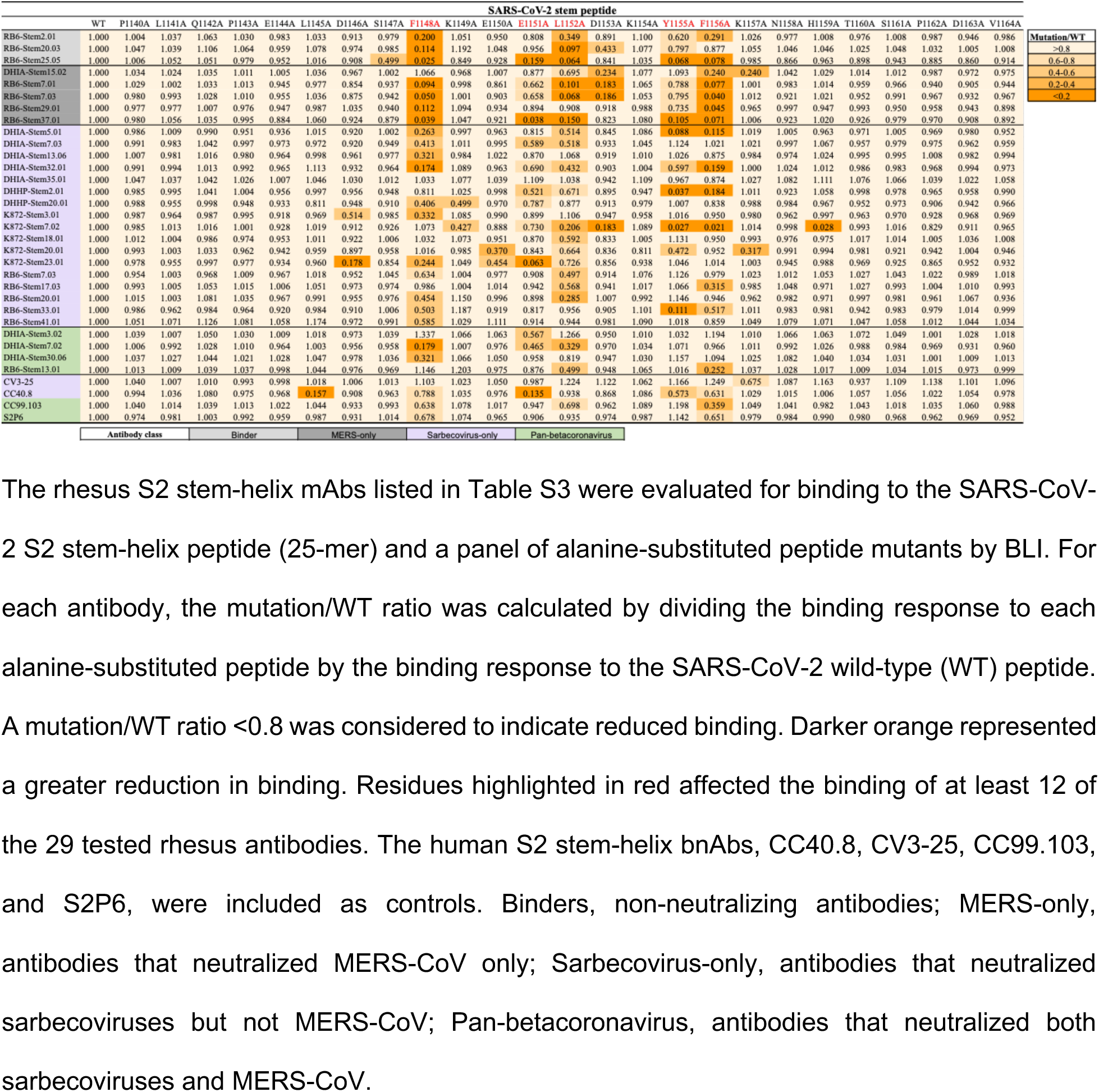
BLI-based epitope mapping of selected rhesus S2 stem-helix mAbs.

**Table S5.**
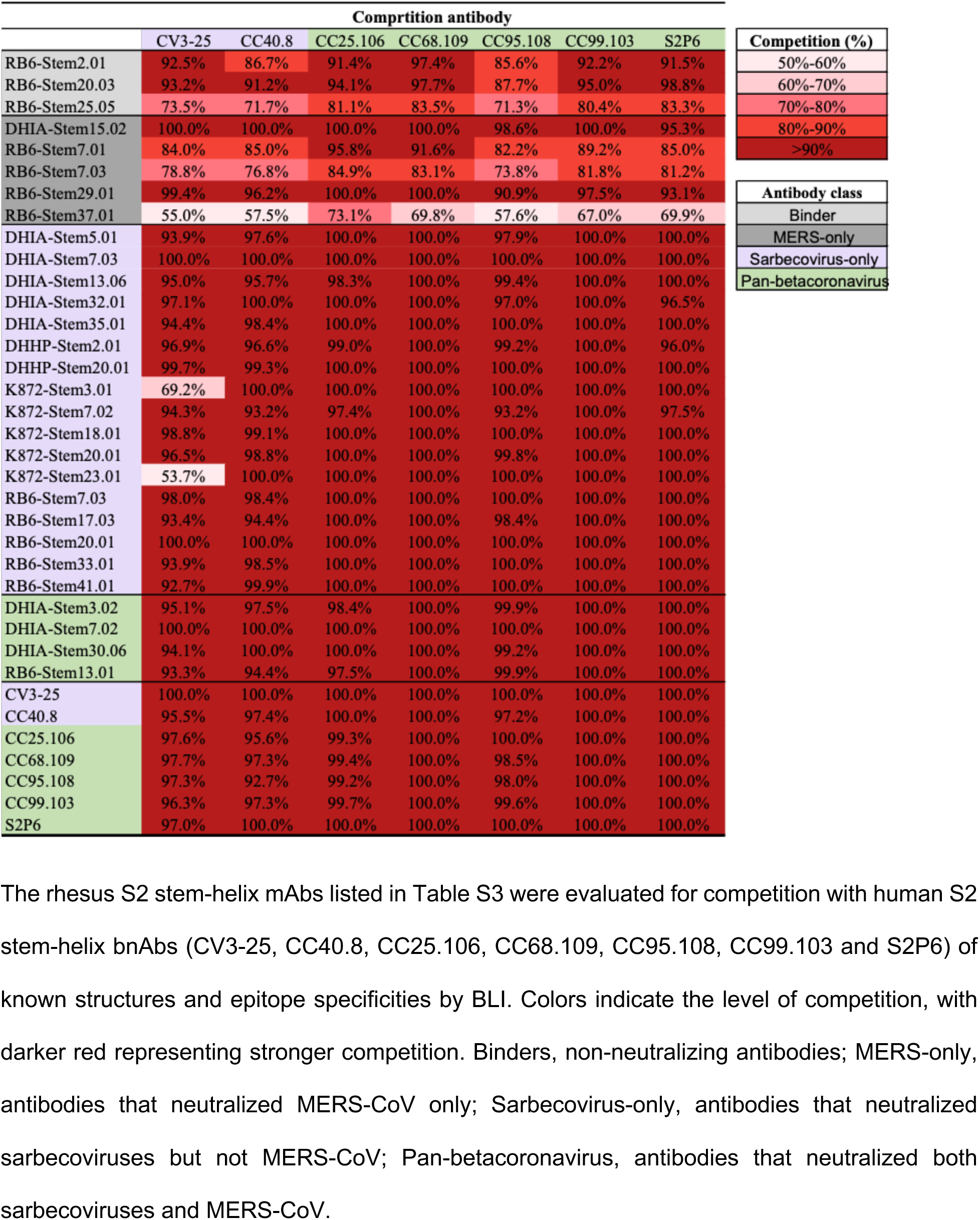
BLI competition epitope binning of selected rhesus S2 stem-helix mAbs.

**Table S6.**
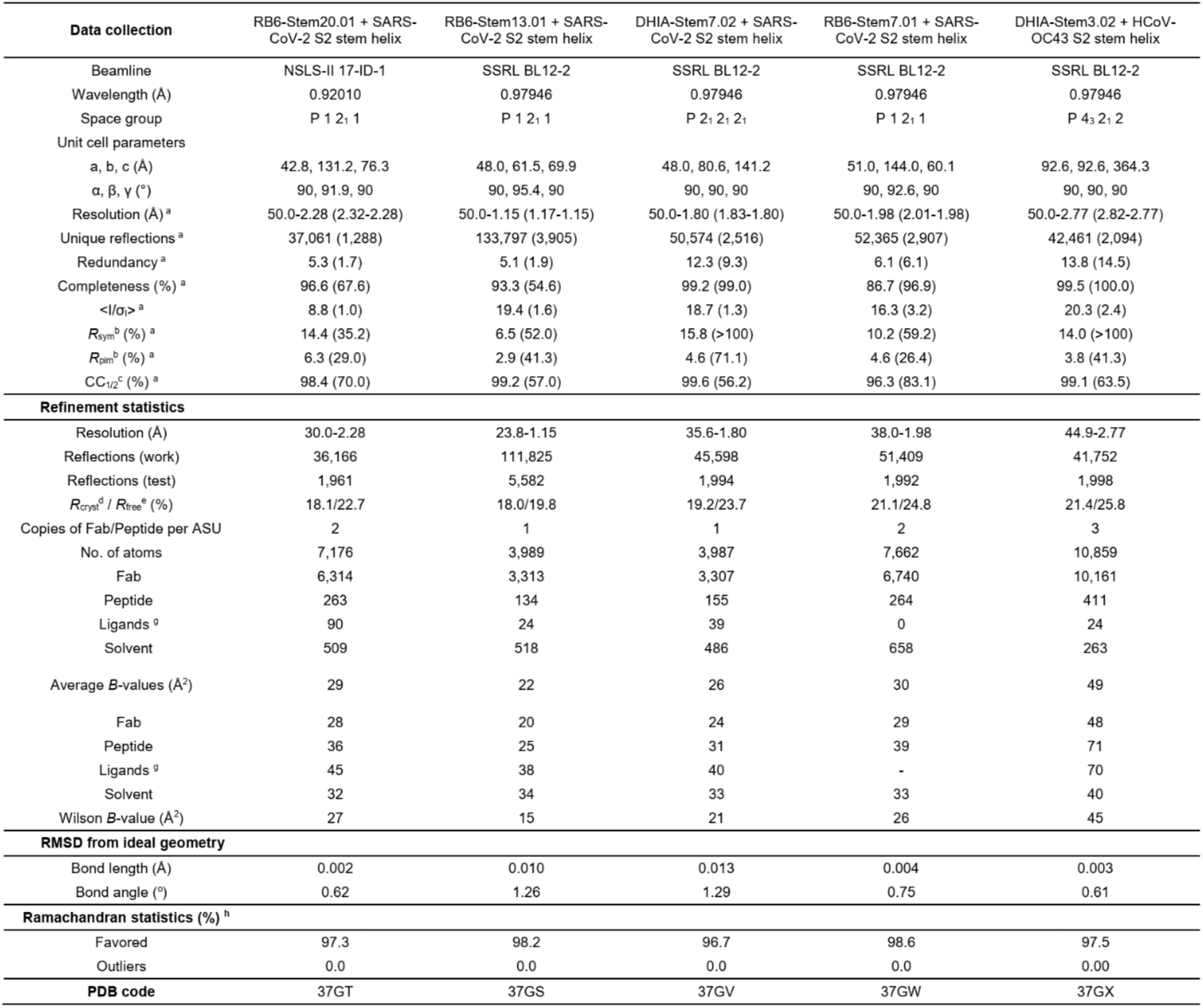
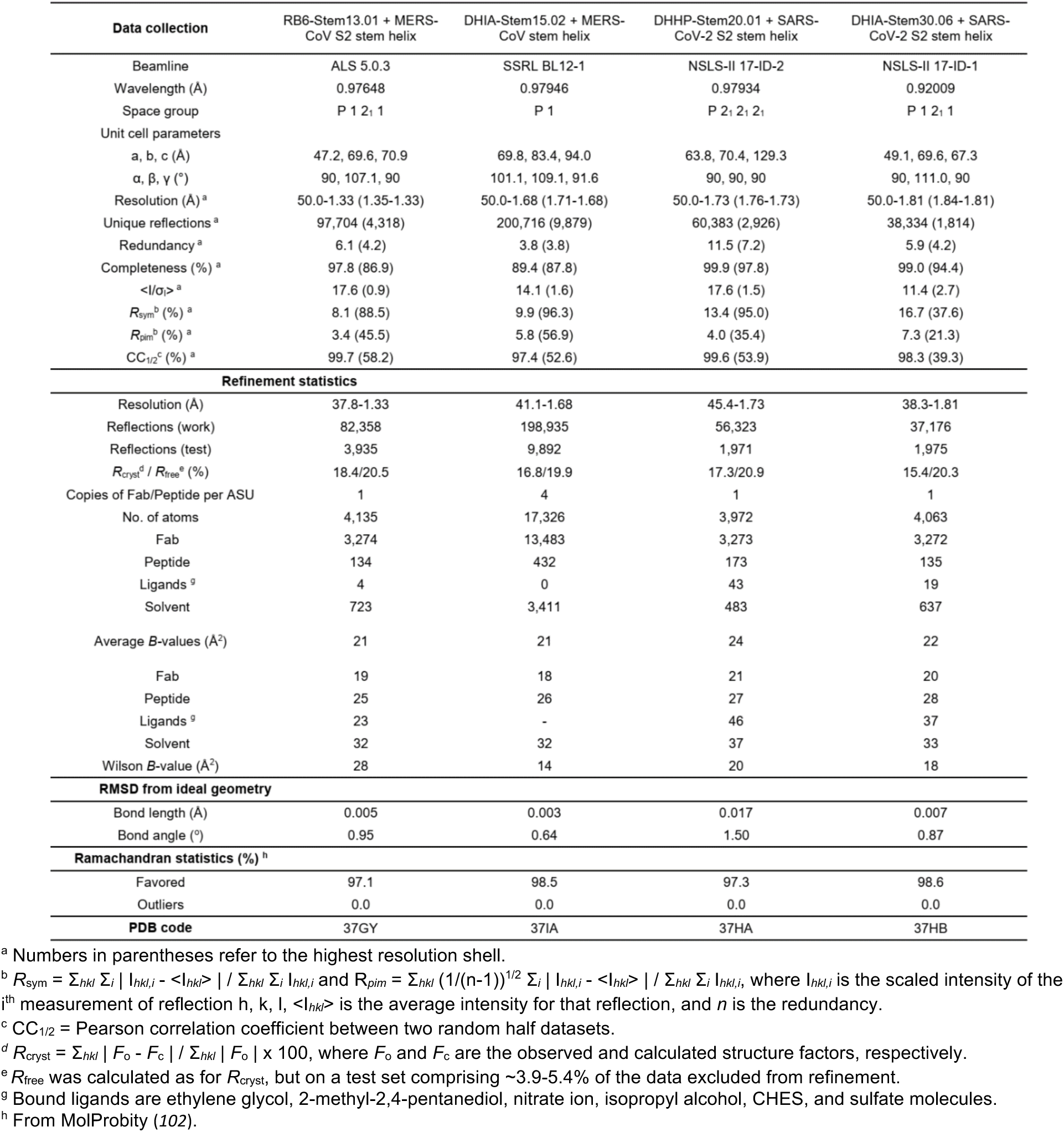
X-ray data collection and refinement statistics.

## REFERENCES AND NOTES

1. C. Drosten et al., Identification of a novel coronavirus in patients with severe acute respiratory syndrome. The New England journal of medicine 348, 1967–1976 (2003).

2. A. M. Zaki, S. van Boheemen, T. M. Bestebroer, A. D. Osterhaus, R. A. Fouchier, Isolation of a novel coronavirus from a man with pneumonia in Saudi Arabia. The New England journal of medicine 367, 1814–1820 (2012).

3. J. F.-W. Chan et al., A familial cluster of pneumonia associated with the 2019 novel coronavirus indicating person-to-person transmission: a study of a family cluster. The Lancet 395, 514–523 (2020).

4. P. Zhou et al., A pneumonia outbreak associated with a new coronavirus of probable bat origin. Nature 579, 270–273 (2020).

5. S. Cankat, M. U. Demael, L. Swadling, In search of a pan-coronavirus vaccine: next-generation vaccine design and immune mechanisms. Cellular & Molecular Immunology 21, 103–118 (2024).

6. Y. Chen et al., Broadly neutralizing antibodies to SARS-CoV-2 and other human coronaviruses. Nature Reviews Immunology 23, 189–199 (2023).

7. J. L. Gordon et al., Development of broadly protective coronavirus vaccines: A joint NIAID-CEPI workshop report. Vaccine 54, 126909 (2025).

8. L. R. Baden et al., Efficacy and Safety of the mRNA-1273 SARS-CoV-2 Vaccine. The New England journal of medicine 384, 403–416 (2020).

9. F. P. Polack et al., Safety and Efficacy of the BNT162b2 mRNA Covid-19 Vaccine. The New England journal of medicine 383, 2603–2615 (2020).

10. Z. Wang et al., mRNA vaccine-elicited antibodies to SARS-CoV-2 and circulating variants. Nature 592, 616–622 (2021).

11. P. B. Gilbert et al., Title: Immune Correlates Analysis of the mRNA-1273 COVID-19 Vaccine Efficacy Trial. Science 375, 43–50 (2021).

12. A. J. Greaney et al., Antibodies elicited by mRNA-1273 vaccination bind more broadly to the receptor binding domain than do those from SARS-CoV-2 infection. Science translational medicine 13, eabi9915 (2021).

13. C. O. Barnes et al., SARS-CoV-2 neutralizing antibody structures inform therapeutic strategies. Nature 588, 682–687 (2020).

14. C. K. Wibmer et al., SARS-CoV-2 501Y.V2 escapes neutralization by South African COVID-19 donor plasma. Nature medicine 27, 622–625 (2021).

15. M. Yuan et al., Structural and functional ramifications of antigenic drift in recent SARS-CoV-2 variants. Science 373, 818–823 (2021).

16. A. A. Cohen et al., Mosaic nanoparticles elicit cross-reactive immune responses to zoonotic coronaviruses in mice. Science 371, 735–741 (2021).

17. A. C. Walls et al., Elicitation of broadly protective sarbecovirus immunity by receptor-binding domain nanoparticle vaccines. Cell 184, 5432–5447.e5416 (2021).

18. Y. Liang et al., Design of a mutation-integrated trimeric RBD with broad protection against SARS-CoV-2. Cell Discovery 8, 17 (2022).

19. D. R. Martinez et al., Chimeric spike mRNA vaccines protect against Sarbecovirus challenge in mice. Science 373, 991–998 (2021).

20. C. Dacon et al., Broadly neutralizing antibodies target the coronavirus fusion peptide. Science 377, 728–735 (2022).

21. M. M. Sauer et al., Structural basis for broad coronavirus neutralization. Nature structural & molecular biology 28, 478–486 (2021).

22. G. Song et al., Cross-reactive serum and memory B-cell responses to spike protein in SARS-CoV-2 and endemic coronavirus infection. Nature communications 12, 2938 (2021).

23. S. Changrob et al., Common cold embecovirus imprinting primes broadly neutralizing antibody responses to SARS-CoV-2 S2. Journal of Experimental Medicine 222, (2025).

24. K. W. Ng et al., SARS-CoV-2 S2–targeted vaccination elicits broadly neutralizing antibodies. Science translational medicine 14, eabn3715 (2022).

25. P. Zhou et al., Broadly neutralizing anti-S2 antibodies protect against all three human betacoronaviruses that cause deadly disease. Immunity, (2023).

26. C.-L. Hsieh et al., Stabilized coronavirus spike stem elicits a broadly protective antibody. Cell reports 37, (2021).

27. P. Zhou et al., A human antibody reveals a conserved site on beta-coronavirus spike proteins and confers protection against SARS-CoV-2 infection. *Science translational medicine*, eabi9215 (2022).

28. K. Dueker et al., Heterologous betacoronavirus spike immunization in non-human primates elicits antibodies that neutralize both sarbeco- and merbecoviruses. Cell reports 45, (2026).

29. D. Pinto et al., Broad betacoronavirus neutralization by a stem helix-specific human antibody. Science 373, 1109–1116 (2021).

30. C. Dacon et al., Rare, convergent antibodies targeting the stem helix broadly neutralize diverse betacoronaviruses. Cell host & microbe, (2022).

31. N. K. Hurlburt et al., Structural definition of a pan-sarbecovirus neutralizing epitope on the spike S2 subunit. Commun Biol 5, 342 (2022).

32. W. Li et al., Structural basis and mode of action for two broadly neutralizing antibodies against SARS-CoV-2 emerging variants of concern. Cell reports 38, (2022).

33. A. B. Kapingidza et al., Engineered immunogens to elicit antibodies against conserved coronavirus epitopes. Nature communications 14, 7897 (2023).

34. W. Shi et al., Vaccine-elicited murine antibody WS6 neutralizes diverse beta-coronaviruses by recognizing a helical stem supersite of vulnerability. Structure 30, 1233–1244.e1237 (2022).

35. C. Wang et al., A conserved immunogenic and vulnerable site on the coronavirus spike protein delineated by cross-reactive monoclonal antibodies. Nature communications 12, 1715 (2021).

36. H. D. Kamp et al., Design of a broadly reactive Lyme disease vaccine. NPJ Vaccines 5, 33 (2020).

37. M. Kanekiyo et al., Rational Design of an Epstein-Barr Virus Vaccine Targeting the Receptor-Binding Site. Cell 162, 1090–1100 (2015).

38. M. Kanekiyo et al., Mosaic nanoparticle display of diverse influenza virus hemagglutinins elicits broad B cell responses. Nature immunology 20, 362–372 (2019).

39. W.-t. He et al., Broadly neutralizing antibodies to SARS-related viruses can be readily induced in rhesus macaques. Science translational medicine 14, eabl9605 (2022).

40. M. Silva, et al., A particulate saponin/TLR agonist vaccine adjuvant alters lymph flow and modulates adaptive immunity. Sci Immunol 6, eabf1152 (2021).

41. L. Stamatatos et al., mRNA vaccination boosts cross-variant neutralizing antibodies elicited by SARS-CoV-2 infection. Science 372, 1413–1418 (2021).

42. W.-t. He et al., Targeted isolation of diverse human protective broadly neutralizing antibodies against SARS-like viruses. Nature immunology 23, 960–970 (2022).

43. R. R. Goel, et al., Distinct antibody and memory B cell responses in SARS-CoV-2 naïve and recovered individuals following mRNA vaccination. Sci Immunol 6, (2021).

44. R. Suryawanshi, M. Ott, SARS-CoV-2 hybrid immunity: silver bullet or silver lining? Nature Reviews Immunology 22, 591–592 (2022).

45. E. Andreano et al., Hybrid immunity improves B cells and antibodies against SARS-CoV-2 variants. Nature 600, 530–535 (2021).

46. M. M. Corcoran et al., Production of individualized V gene databases reveals high levels of immunoglobulin genetic diversity. Nature communications 7, 13642 (2016).

47. Y. Cai et al., Distinct conformational states of SARS-CoV-2 spike protein. Science 369, 1586–1592 (2020).

48. P. Zhou et al., A human antibody reveals a conserved site on beta-coronavirus spike proteins and confers protection against SARS-CoV-2 infection. Science Translational Medicine 14, eabi9215 (2022).

49. P. Zhou et al., Broadly neutralizing anti-S2 antibodies protect against all three human betacoronaviruses that cause deadly disease. Immunity 56, 669–686.e667 (2023).

50. M. W. Grunst et al., Structure and inhibition of SARS-CoV-2 spike refolding in membranes. Science 385, 757–765 (2024).

51. C. Akıl, J. Xu, J. Shen, P. Zhang, Unveiling the structural spectrum of SARS-CoV-2 fusion by in situ cryo-ET. Nature communications 16, 5150 (2025).

52. C. Dacon et al., Rare, convergent antibodies targeting the stem helix broadly neutralize diverse betacoronaviruses. Cell Host & Microbe 31, 97–111.e112 (2023).

53. D. Pinto et al., Broad betacoronavirus neutralization by a stem helix–specific human antibody. Science 373, 1109–1116 (2021).

54. T. F. Rogers et al., Isolation of potent SARS-CoV-2 neutralizing antibodies and protection from disease in a small animal model. Science 369, 956–963 (2020).

55. D. F. Robbiani et al., Convergent antibody responses to SARS-CoV-2 in convalescent individuals. Nature 584, 437–442 (2020).

56. M. Yuan et al., Structural basis of a shared antibody response to SARS-CoV-2. Science 369, 1119-1123 (2020).

57. P. J. M. Brouwer et al., Potent neutralizing antibodies from COVID-19 patients define multiple targets of vulnerability. Science 369, 643–650 (2020).

58. G. Song et al., Broadly neutralizing antibodies targeting a conserved silent face of spike RBD resist extreme SARS-CoV-2 antigenic drift. Cell reports, (2025).

59. A. Iwasaki, Exploiting Mucosal Immunity for Antiviral Vaccines. Annual review of immunology 34, 575–608 (2016).

60. M. W. Russell, Z. Moldoveanu, P. L. Ogra, J. Mestecky, Mucosal Immunity in COVID-19: A Neglected but Critical Aspect of SARS-CoV-2 Infection. Frontiers in immunology 11, 611337 (2020).

61. W. Hong et al., Mucosal immunity and vaccine development. Signal Transduction and Targeted Therapy 11, 301 (2026).

62. P. C. H. Tang, Z. Ling, M. Lee, N. J. C. King, S. Mahalingam, Combating respiratory diseases with mucosal vaccines. Journal of virology 100, e0014626 (2026).

63. T. Mao et al., Unadjuvanted intranasal spike vaccine elicits protective mucosal immunity against sarbecoviruses. Science 378, eabo2523 (2022).

64. H. Kiyono, P. B. Ernst, Nasal vaccines for respiratory infections. Nature 641, 321–330 (2025).

65. H. Shin, A. Iwasaki, A vaccine strategy that protects against genital herpes by establishing local memory T cells. Nature 491, 463–467 (2012).

66. S. Uematsu, Programming systemic and mucosal immunity through co-adjuvant-based prime-boost vaccination. Current Opinion in Virology 76, 101525 (2026).

67. N. Pardi, F. Krammer, mRNA vaccines for infectious diseases - advances, challenges and opportunities. Nat Rev Drug Discov 23, 838–861 (2024).

68. A. Toniolo, G. Maccari, G. Camussi, mRNA Technology and Mucosal Immunization. Vaccines (Basel) 12, (2024).

69. P. D. Kwong, J. R. Mascola, HIV-1 Vaccines Based on Antibody Identification, B Cell Ontogeny, and Epitope Structure. Immunity 48, 855–871 (2018).

70. K. Xu et al., Epitope-based vaccine design yields fusion peptide-directed antibodies that neutralize diverse strains of HIV-1. Nature medicine 24, 857–867 (2018).

71. F. Krammer, The human antibody response to influenza A virus infection and vaccination. Nature Reviews Immunology 19, 383–397 (2019).

72. A. Impagliazzo et al., A stable trimeric influenza hemagglutinin stem as a broadly protective immunogen. Science 349, 1301–1306 (2015).

73. D. R. Burton, L. Hangartner, Broadly Neutralizing Antibodies to HIV and Their Role in Vaccine Design. Annual review of immunology 34, 635–659 (2016).

74. S. J. Krebs et al., Longitudinal Analysis Reveals Early Development of Three MPER-Directed Neutralizing Antibody Lineages from an HIV-1-Infected Individual. Immunity 50, 677–691.e613 (2019).

75. S. Henikoff, J. G. Henikoff, Amino acid substitution matrices from protein blocks. Proceedings of the National Academy of Sciences 89, 10915–10919 (1992).

## REFERENCES

76. J. Pallesen et al., Immunogenicity and structures of a rationally designed prefusion MERS-CoV spike antigen. Proceedings of the National Academy of Sciences of the United States of America 114, E7348–E7357 (2017).

77. D. Wrapp et al., Cryo-EM structure of the 2019-nCoV spike in the prefusion conformation. Science 367, 1260–1263 (2020).

78. X. Zhou et al., Diverse immunoglobulin gene usage and convergent epitope targeting in neutralizing antibody responses to SARS-CoV-2. Cell reports 35, 109109 (2021).

79. T. Tiller et al., Efficient generation of monoclonal antibodies from single human B cells by single cell RT-PCR and expression vector cloning. Journal of immunological methods 329, 112–124 (2008).

80. J. Gorman et al., Isolation and Structure of an Antibody that Fully Neutralizes Isolate SIVmac239 Reveals Functional Similarity of SIV and HIV Glycan Shields. Immunity 51, 724–734.e724 (2019).

81. Z. Otwinowski, W. Minor, Processing of X-ray diffraction data collected in oscillation mode. Methods Enzymol 276, 307–326 (1997).

82. A. J. McCoy et al., Phaser crystallographic software. J Appl Crystallogr 40, 658–674 (2007).

83. P. Emsley, B. Lohkamp, W. G. Scott, K. Cowtan, Features and development of Coot. Acta Crystallogr D Biol Crystallogr 66, 486–501 (2010).

84. P. D. Adams et al., PHENIX: a comprehensive Python-based system for macromolecular structure solution. Acta Crystallogr D Biol Crystallogr 66, 213–221 (2010).

85. E. Krissinel, K. Henrick, Inference of macromolecular assemblies from crystalline state. Journal of molecular biology 372, 774–797 (2007).

86. C. Suloway et al., Automated molecular microscopy: the new Leginon system. J Struct Biol 151, 41–60 (2005).

87. N. R. Voss, C. K. Yoshioka, M. Radermacher, C. S. Potter, B. Carragher, DoG Picker and TiltPicker: software tools to facilitate particle selection in single particle electron microscopy. J Struct Biol 166, 205–213 (2009).

88. T. Ogura, K. Iwasaki, C. Sato, Topology representing network enables highly accurate classification of protein images taken by cryo electron-microscope without masking. J Struct Biol 143, 185–200 (2003).

89. S. R. Leist et al., A Mouse-Adapted SARS-CoV-2 Induces Acute Lung Injury and Mortality in Standard Laboratory Mice. Cell 183, 1070–1085 e1012 (2020).

90. V. D. Menachery, L. E. Gralinski, R. S. Baric, M. T. Ferris, New Metrics for Evaluating Viral Respiratory Pathogenesis. PloS one 10, e0131451 (2015).

91. A. S. Cockrell et al., A mouse model for MERS coronavirus-induced acute respiratory distress syndrome. Nat Microbiol 2, 16226 (2016).

92. M. G. Douglas, J. F. Kocher, T. Scobey, R. S. Baric, A. S. Cockrell, Adaptive evolution influences the infectious dose of MERS-CoV necessary to achieve severe respiratory disease. Virology 517, 98–107 (2018).

93. R. C. Edgar, MUSCLE: a multiple sequence alignment method with reduced time and space complexity. BMC Bioinformatics 5, 113 (2004).

94. R. C. Edgar, MUSCLE: multiple sequence alignment with high accuracy and high throughput. Nucleic acids research 32, 1792–1797 (2004).

95. U. Bodenhofer, E. Bonatesta, C. Horejš-Kainrath, S. Hochreiter, msa: an R package for multiple sequence alignment. Bioinformatics 31, 3997–3999 (2015).

96. E. Stansell, R. C. Desrosiers, Fundamental difference in the content of high-mannose carbohydrate in the HIV-1 and HIV-2 lineages. Journal of virology 84, 8998–9009 (2010).

97. E. Fouquaert et al., Related lectins from snowdrop and maize differ in their carbohydrate-binding specificity. Biochemical and biophysical research communications 380, 260–265 (2009).

98. E. Krissinel, K. Henrick, Inference of macromolecular assemblies from crystalline state. J. Mol. Biol. 372, 774–797 (2007).

99. F. Sievers et al., Fast, scalable generation of high-quality protein multiple sequence alignments using Clustal Omega. Molecular Systems Biology 7, 539 (2011).

100. M.-P. Lefranc, IMGT, the international ImMunoGeneTics database ®. Nucleic Acids Research 31, 307–310 (2003).

101. J. Ye, N. Ma, T. L. Madden, J. M. Ostell, IgBLAST: an immunoglobulin variable domain sequence analysis tool. Nucleic Acids Research 41, W34–W40 (2013).

102. C. J. Williams et al., MolProbity: More and better reference data for improved all-atom structure validation. Protein science : a publication of the Protein Society 27, 293–315 (2018).

